# Variable patterns of retrotransposition in different HeLa strains provide mechanistic insights into SINE RNA mobilization processes

**DOI:** 10.1101/2024.05.03.592410

**Authors:** John B. Moldovan, Huira C. Kopera, Ying Liu, Marta Garcia-Canadas, Purificacion Catalina, Paola E. Leone, Laura Sanchez, Jacob O. Kitzman, Jeffrey M. Kidd, Jose Luis Garcia-Perez, John V. Moran

## Abstract

*Alu* elements are non-autonomous Short INterspersed Elements (SINEs) derived from the *7SL RNA* gene that are present at over one million copies in human genomic DNA. *Alu* mobilizes by a mechanism known as retrotransposition, which requires the Long INterspersed Element-1 (LINE-1 or L1) *ORF2*-encoded protein (ORF2p). Here, we demonstrate that HeLa strains differ in their capacity to support *Alu* retrotransposition. Human *Alu* elements retrotranspose efficiently in HeLa-HA and HeLa-CCL2 (*Alu*-permissive) strains, but not in HeLa-JVM or HeLa-H1 (*Alu*-nonpermissive) strains. A similar pattern of retrotransposition was observed for other *7SL RNA*-derived SINEs and *tRNA*-derived SINEs. In contrast, mammalian LINE-1s, a zebrafish LINE, a human *SINE-VNTR*-*Alu* (*SVA*) element, and an *L1 ORF1*-containing messenger RNA can retrotranspose in all four HeLa strains. Using an *in vitro* reverse transcriptase-based assay, we show that *Alu* RNAs associate with ORF2p and are converted into cDNAs in both *Alu*-permissive and *Alu*-nonpermissive HeLa strains, suggesting that *7SL*- and *tRNA*-derived SINE RNAs use strategies to ‘hijack′ L1 ORF2p that are distinct from those used by *SVA* elements and *ORF1*-containing mRNAs. These data further suggest ORF2p associates with the *Alu* RNA poly(A) tract in both *Alu*-permissive and *Alu*-nonpermissive HeLa strains, but that *Alu* retrotransposition is blocked after this critical step in *Alu*-nonpermissive HeLa strains.

## INTRODUCTION

*Alu* retrotransposon sequences are a class of primate specific non-autonomous Short INterspersed Elements (SINEs) that are present at over one million copies in the human genome (1,2). *Alu* elements evolved from the 5′ *Alu* domain of the *7SL RNA* gene (3) and can mobilize throughout the genome by a replicative process known as retrotransposition (4,5). *Alu* RNAs do not encode proteins and require enzymatic activities encoded by the second open reading frame (*ORF2*) of Long INterspersed Element-1 (LINE-1 or L1) retrotransposons to mediate their retrotransposition (4). *Alu* retrotransposition events are responsible for rare cases of genetic disease (1,6) and it is estimated that a *de novo Alu* retrotransposition event occurs in approximately 1 of 40 live human births (7,8), making it the most active retrotransposon in the extant human population.

*Alu* elements are classified into several subfamilies based upon their sequence and evolutionary age (1). *Alu*Y elements (*e.g.*, *Alu*Ya5 and *Alu*Yb8) represent the youngest and currently active human *Alu* subfamily. *Alu*Y elements are approximately 280 nucleotides (nts) in length and consist of two *7SL Alu* domains that are arranged in a head-to-tail manner, which are separated by a short adenosine (A)-rich sequence (1,3). The 3′ end of *Alu* elements typically contain a homopolymeric adenosine (poly[A]) tract that varies in length; *Alu* elements from younger subfamilies (*e.g.*, *Alu*Y) generally contain longer 3′ poly(A) tracts than *Alu* elements from older subfamilies (*e.g.*, *Alu*S and *Alu*J) (9). Modern dimeric *Alu* elements likely evolved from fusion events between monomeric *7SL Alu* domain RNAs known as Fossil *Alu* Monomers (FAMs), which descended directly from the *7SL RNA* gene (10,11). Although FAMs are not thought to be currently active in humans, an engineered *Alu*J left monomer consensus sequence can efficiently retrotranspose in cultured cell-based retrotransposition assays (12).

A round of *Alu* retrotransposition begins with transcription of a genomic *Alu* sequence by RNA polymerase (pol) III. *Alu* transcription is dependent on the RNA pol III A/B box promoter present within its left monomer (13) as well as genomic DNA sequences upstream of the *Alu* element (14–16). *Alu* transcription termination requires an RNA pol III terminator sequence of at least four consecutive thymidine (T) residues located in genomic DNA, downstream of the *Alu* 3′ poly(A) tract (17). Following transcription, *Alu* RNAs form a RiboNucleoprotein Particle (RNP) complex, containing the Signal Recognition Particle proteins 9 and 14 (SRP 9/14) and likely other host proteins, which enable *Alu* RNAs to localize to ribosomes (4,12,18,19). It is hypothesized that ribosomal localization allows the *Alu* RNA 3′ poly(A) tract to compete with the L1 RNA 3′ poly(A) tract for the co-translational binding of L1 *ORF2*-encoded protein (ORF2p) (12,18,20). Upon binding ORF2p, and perhaps other host factors, the resultant *Alu* RNA/ORF2p RNPs then gain nuclear access, where a new *Alu* copy is integrated into the genome by Target site-Primed Reverse Transcription (TPRT) (4,21).

Using a strategy similar to that used to monitor LINE-1 retrotransposition (22), Dewannieux and colleagues developed a cultured cell *Alu* retrotransposition assay (4), which has been instrumental in elucidating several aspects of *Alu* biology (12,23,24). The use of this assay, as well as similarly designed *trans*-complementation-based retrotransposition assays, demonstrated that L1 ORF2p expression is required to mediate *Alu* retrotransposition (4), as well as the mobilization of other SINE RNAs and messenger RNAs (mRNAs), with the latter leading to the formation of processed pseudogenes (25–30).

Here, we used genetic, molecular biological, and biochemical assays to gain insight into the mobilization mechanism of *7SL*-derived SINEs, *tRNA*-derived SINEs, and cellular mRNAs by the L1 retrotransposition machinery in different HeLa cell strains. We demonstrate that two *Alu*-permissive HeLa strains (HeLa-HA, HeLa-CCL2) support robust *Alu* retrotransposition in *trans*, whereas two *Alu*-nonpermissive HeLa strains (HeLa-JVM, HeLa-H1) do not support *Alu* retrotransposition. The retrotransposition of rodent *7SL*- and *tRNA*-derived SINEs, as well as a *tRNA*-derived canine SINE, also is dramatically decreased in *Alu*-nonpermissive HeLa strains. By comparison, all four HeLa strains support the retrotransposition of vertebrate LINEs, a *SINE-VNTR*-*Alu* element (*SVA*) non-autonomous retrotransposon, and a mammalian *L1 ORF1*-containing mRNA (*ORF1mneoI*), which structurally resembles a rat “Half of An L1” (*HAL1*) non-autonomous retrotransposon (26,31). Whole genome sequencing and karyotype analyses further indicate that *Alu*-permissive and *Alu*-nonpermissive HeLa strains differ significantly in DNA copy number. Thus, these data strongly suggest that *7SL*- and *tRNA*-derived SINEs retrotranspose by a similar mechanism that is fundamentally distinct from the mobility mechanisms used by *SVA* elements and/or *L1 ORF1*-containing mRNAs.

## MATERIALS & METHODS

### Cell Culture Conditions

All HeLa strains were grown in high-glucose DMEM (Gibco) supplemented with 10% Fetal Bovine Serum (FBS) (Gibco), 100 U/mL penicillin-streptomycin (Invitrogen), and 0.29 mg/mL L-glutamine (Gibco). The HeLa-JVM strain originally was obtained from the laboratory of Dr. Maxine Singer (Bethesda, MD, US). The HeLa-HA strain was a kind gift from the laboratory of Dr. Astrid Roy-Engel (New Orleans, LA, US). The HeLa-CCL2 and HeLa-H1 strains were obtained from the American Type Culture Collection (ATCC). All HeLa strains were maintained at 37°C with 7% CO_2_. We regularly screened cells for the absence of *Mycoplasma spp.* and used Short Tandem Repeat (STR) analyses to confirm the identity of cultured cells.

#### Plasmids

Oligonucleotide sequences used in this study and cloning strategies are available upon request. All human L1 plasmids contain the indicated fragments of L1.3 DNA (a retrotransposition-competent human L1, GenBank accession no. L19088 (32)) cloned into pCEP4 (Invitrogen) unless indicated otherwise. All LINE and cDNA plasmids contain the CMV promoter (upstream) and the SV40 polyadenylation signal (downstream) unless noted otherwise. Plasmid DNAs were prepared using a Midiprep Plasmid DNA Kit (Qiagen).

<u>pJM101/L1.3</u>: is a pCEP4-based plasmid that contains an active human L1 (L1.3) equipped with a *mneoI* retrotransposition indicator cassette (22,32,33).

<u>pJM105/L1.3</u>: is derivative of pJM101/L1.3 that contains a D702A (GAC to GCU) missense mutation in the Reverse Transcriptase (RT) active site of L1.3 ORF2 (26).

<u>pL1.3*neo*^Tet^</u>: is a derivative of pJM101/L1.3 where the *mneoI* reporter cassette was replaced by the *neo*^Tet^ retrotransposition indicator cassette (34). The *neo*^Tet^ retrotransposition indicator cassette has been described previously and it was a generous gift from Dr. Thierry Heidmann (Villejuif, France).

<u>p*Aluneo*^Tet^</u>: expresses an *Alu*Y element that is marked with the *neo*^Tet^ retrotransposition reporter cassette. The reporter was subcloned upstream of a 44 nucleotide (nt) long poly(A) tract (4,34). *Alu* expression is augmented by the *7SL RNA* enhancer located upstream of the *Alu* sequence and an RNA polymerase III terminator located downstream of the *Alu* poly(A) sequence. This construct was provided by Dr. Thierry Heidmann (Villejuif, France).

<u>pAJL</u>: expresses a highly active consensus *Alu*J element, consisting of only the *Alu*J left hand monomer, and is tagged with the *neo*^Tet^ retrotransposition reporter cassette (12).

<u>pSINEC_Cf</u>: expresses a canine consensus *tRNA*-derived SINE tagged with the *neo*^Tet^ retrotransposition reporter cassette (28).

<u>Q19-33-*Alu*Ya5</u>: is a pBluescript II KS(−) (Agilent) based vector that expresses an active *Alu*Ya5 element, which was isolated from a mutagenic insertion into the *NF1* gene (6), and is tagged with the *neo*^Tet^ reporter cassette. *Alu* expression is driven by the *7SL* promoter/enhancer located upstream of *Alu* and a 33 nt long poly(A) tract that was subcloned downstream of the *neo*^Tet^ retrotransposition reporter cassette. To facilitate *Alu* expression, a *BC1* RNA polymerase III terminator (35) was subcloned downstream of the *Alu* poly(A) sequence.

<u>Q19-33-Sx3</u>: is a derivative of Q19-33-*Alu*Ya5 that contains an active *Alu*Sx3 core sequence replacing *Alu*Ya5. The *Alu*Sx3 element was PCR amplified using genomic DNA from HeLa cells and primers *Alu*Sx3FWD (5′-*<u>ACCGGT</u>*GGCCGGGCGCGGTGGCTC-3′) and *Alu*Sx3Rev (5′-*<u>GTATAC</u>*GAGACGGAGTCTCGCTCTGTC-3′).

<u>Q19-33-*B1*-Mur4</u>: is a derivative of Q19-33-*Alu*Ya5 that contains an active mouse *B1*_Mur4 element replacing *Alu*Ya5. The *B1* element was isolated from a recent mutagenic insertion in mice (36).

<u>Q19-33-*B2*-Mm1a</u>: is a derivative of Q19-33-*Alu*Ya5 that contains an active mouse *B2*_Mm1a consensus sequence replacing *Alu*Ya5. The *B2* element was isolated from genomic DNA from a C57BL mice (genomic coordinates in mm10: chr6 (+): 149008364-149008544).

*<u>BC200</u>*<u>con1</u>: was derived from p*Aluneo*^Tet^ and expresses a *BC200 RNA* consensus sequence (*BC200*con1) (37) that is marked with the *neo*^Tet^ reporter cassette. The reporter was subcloned upstream of a 44 nt long poly(A) tract (4,34). *BC200*con1 RNA expression is augmented by the *7SL RNA* enhancer located in front of the *BC200*con1 sequence and an RNA polymerase III terminator after the 44 nt poly(A) sequence.

<u>pAD26</u>: Was a kind gift from Dr. Annette Damert (Göttingen, Germany); it is a pCEP4-based plasmid that expresses the highly active human *SVA*_E H8_43 element tagged with the *NEO* retrotransposition indicator cassette (38).

<u>pCEP/GFP</u>: is a pCEP4-based plasmid that contains the humanized enhanced green fluorescent protein (*hrGFP*) coding sequence from *phrGFP-C* (Agilent) under the control of the pCEP4 CMV promoter and SV40 polyadenylation signal (39).

<u>pCEPsmL1</u>: is a pCEP4-based plasmid that expresses a codon optimized full-length active mouse element from the L1MdTf_I subfamily and is tagged with the *mneoI* indicator cassette (40); the construct was a generous gift from Dr. Jef Boeke (New York City, NY, US).

<u>pTMF3</u>: is a derivative of pJM101/L1.3 except that it was modified to contain a T7 *gene10* epitope-tag on the carboxyl-terminus of ORF1p and a 3XFLAG epitope-tag on the carboxyl-terminus of ORF2p (41). The 3′-UTR contains the *mneoI* retrotransposition indicator cassette.

<u>pTMF3Δneo</u>: is a derivative of pTMF3 that lacks the *mneoI* indicator cassette (41).

<u>pTMO2F3</u>: is like pTMF3 except that the T7-tagged *ORF1* has been deleted and the 3′-UTR does not contain the *mneoI* retrotransposition indicator cassette (41).

<u>pTMO2F3(D702A)</u>: is derivative of TFO2F3 that contains a D702A (GAC to GCU) missense mutation in the RT active site of L1.3 ORF2 (41).

<u>p5UT-hORF2</u>: is a pCEP4-based “driver” plasmid that expresses human *L1 ORF2* (from L1.3); *L1 ORF2* expression is augmented by a CMV promoter and the native L1 5′UTR located upstream of the *L1 ORF2* sequence and the SV40 polyadenylation signal located at the 3′ end of the *L1 ORF2* sequence (39).

<u>pJBM560</u>: is a derivative of pJM101/L1.3 that contains the L1 5’UTR, *L1 ORF1*, and a version of the L1 3’UTR containing the *mneoI* retrotransposition indicator cassette.

<u>pJBM560^AA^</u>: is a derivative of pJBM560 except that *ORF1* contains the mutations R261A and R262A in the RNA binding domain of *L1 ORF1p* (22,42). The 3′-UTR contains the *mneoI* retrotransposition indicator cassette.

<u>pJBM561</u>: is a derivative of pJBM560 that lacks the *mneoI* retrotransposition cassette.

<u>pEGFP*mneoI*</u>: is a pCEP4 based vector containing the enhanced green fluorescent protein *(EGFP) ORF* sequence from plasmid pEGFP-C3 (Clontech) followed by the *mneoI* retrotransposition indicator cassette. The *EFGP* and *mneoI* sequences are cloned in between the pCEP4 CMV promoter and SV40 polyadenylation sequences.

<u>pZfL2-2</u>: is a pCEP4 based plasmid that contains the *ORF* and 3′UTR (235 bp long) from an active zebrafish ZfL2-2 element (ZL15, accession no. A*B2*11150); a copy of the *mneoI* indicator cassette was cloned in the 3′UTR, 106 bp downstream of the *ORF* stop codon (43).

<u>pTG_F_21</u>: is a pCEP4 based plasmid that contains a 8.8-kb fragment containing a full length mouse *TG_F_21* L1 element that contains the *mneoI* indicator cassette cloned downstream of the 3′UTR of *TG_F_21* (44).

<u>pA2</u>: is a derivative of plasmid Q19-33-*Alu*Ya5 where an *L1 ORF2* expression cassette was inserted downstream of the *BC1* RNA polymerase III terminator, in the opposite transcriptional orientation as the tagged *Alu*Ya5 cassette. *L1 ORF2* expression is augmented by a CMV promoter and the native L1 5′ UTR located upstream of the *ORF2* sequence, and the SV40 polyadenylation signal located at the 3′ end of the *L1 ORF2* sequence.

#### LINE Retrotransposition Assays

The cultured cell retrotransposition assay was carried out essentially as described (22,45,46). For retrotransposition assays with LINE constructs tagged with *mneoI* or *neo*^Tet^, cells were seeded at approximately 2×10^4^ cells/well (unless otherwise noted) in a 6-well plate (Corning). Within 24 hours, each well was transfected with 1 μg of plasmid DNA (0.5 μg L1 plasmid + 0.5 μg pCEP/GFP) using 4 μL of FuGENE 6 transfection reagent (Promega); the following day, the transfection media was removed, and fresh media was added to each well. To control for transfection efficiency, cells from replicate plates were trypsinized 72 hours after transfection and subjected to flow cytometry analyses to monitor GFP expression using an Accuri™ C6 flow cytometer (BD Biosciences). Three days post-transfection, cells were grown in the presence of media containing G418 (Gibco) (400μg/mL) for 10-12 days; as needed, media was replaced with fresh media approximately every other day. After approximately 12 days of selection, cells were washed with 1x Phosphate Buffered Saline (1xPBS), fixed, and then stained with crystal violet to visualize colonies. The number of G418-resistant colonies for each experimental condition was determined by dividing the number of G418-resistant colonies enumerated in a single well of a 6-well plate by the normalized GFP expression (normalized GFP expression = measured %GFP expression value for each experimental condition divided by the highest %GFP expression value).

#### SINE Retrotransposition Assays

For SINE *trans*-based retrotransposition assays (4,28,46,47), approximately 2-3×10^5^ cells were plated per well in 6-well plates (Corning) and transfected with 0.25 μg of “driver” plasmid (pTMO2F3, p5UThORF2, or other *L1 ORF2p* plasmids) + 0.5 μg of “reporter” plasmid (p*Aluneo*^Tet^ or other SINE reporter plasmids) + 0.25 μg of pCEP/GFP using 3 μL FuGENE® HD (Promega); the following day, the transfection media was removed and fresh media was added to each well. Three days post-transfection, cells were grown in the presence of G418 (400μg/mL) to select for *Alu* retrotransposition events; as needed, media was replaced with fresh media during selection. After approximately 12 days of selection, cells were washed with 1x PBS, fixed, and then stained with crystal violet to visualize colonies. To control for transfection efficiency, cells from replicate plates were trypsinized 72 hours after transfection and subjected to flow cytometry analyses to monitor GFP expression using an Accuri™ C6 flow cytometer (BD Biosciences). The number of G418-resistant colonies for each experimental condition was determined by dividing the number of G418-resistant colonies enumerated in a single well of a 6-well plate by the normalized GFP expression (normalized GFP expression = measured %GFP expression v*alu*e for each experimental condition divided by the highest %GFP expression v*alu*e).

When indicated, some *trans*-based retrotransposition assays were conducted in T-75 flasks (Corning). Briefly, approximately 5-9×10^5^ cells were plated per flask (Corning) and transfected with 1 μg of “driver” plasmid (pTMO2F3, p5UThORF2 or other “driver”) + 3 μg of “reporter” plasmid (p*Aluneo*^Tet^, or other “reporters”) using 12 μL FuGENE® 6 (Promega); the following day, the transfection media was removed, and fresh media was added to each well. Transfection efficiency determination, G418 selection, fixation, and foci staining were carried out as described above. Similarly, some *trans*-retrotransposition assays were carried out in 100mm tissue culture plates (Corning). In these instances, we plated 2×10^5^ cells per plate and we transfected cells using 4μg of DNA (3 μg “reporter” + 1 μg “driver”) and 12μL of FuGENE® 6. Transfection conditions, G418 selection, fixation, and staining were carried out as described above.

#### PCR-Based Retrotransposition Assays

Approximately 2-3×10^5^ HeLa cells were plated per well in 6-well plates (Corning). For *Alu* assays, HeLa cells were co-transfected with 0.5 μg of “driver” plasmid (pTMO2F3) + 0.5 μg of p*Aluneo*^Tet^ using 3 μL FuGENE® HD (Promega); the following day, the transfection media was removed, and fresh media was added to each well. For L1 assays, HeLa cells were transfected with 1 μg of pTMF3. After approximately 7 days, genomic DNA was extracted and cut with the restriction enzymes *Nhe*I and *Eco*NI, which cut within the p*Aluneo*^Tet^ intron, to reduce amplification of plasmid DNA. PCR was carried out on 200-300 ng of restricted genomic DNA using OneTaq 2X Master Mix (NEB) with primers complementary to sequences within the neo cassette that flanked the intron [Neotet1s: 5′-AGTCCCTTCCCGCTTCAGTGACAAC-3′) and (Neotet1as: 5′-CCTCGGCCTCTGAGCTATTC-3′)] using the following thermal cycler conditions: an initial cycle of 94°C for 30 seconds, followed by 35 cycles of 15 seconds at 94°C, 30 seconds at 56°C, and 4 seconds at 68°C with a final cycle of 68°C for 5 minutes. PCR products were visualized on 1% agarose gels stained with ethidium bromide.

#### *SVA* Retrotransposition Assays

*SVA trans-*retrotransposition assays were carried out essentially as described (30,38) Briefly, approximately 2-3×10^5^ HeLa cells were plated per well in 6-well plates (Corning) and transfected with 0.5 μg of “driver” plasmid + 0.5 μg of *SVA “*reporter” plasmid (pAD26) using 3 μL FuGENE® HD (Promega); the following day, the transfection media was removed and fresh media was added to each well. Three days post-transfection, cells were grown in the presence of G418 (400μg/mL) to select for *SVA* retrotransposition events; during selection, the old media was replaced with fresh media containing G418. After approximately 12 days of selection, cells were washed with 1x PBS, fixed, and then stained with crystal violet to visualize colonies. G418-resistant colonies were then enumerated and an average of raw colony counts was reported for pAD26 retrotransposition.

#### *ORF1mneoI* Retrotransposition Assays

For ORF1*mneoI trans*-complementation based assays (26,48), approximately 2-3×10^5^ HeLa cells were plated per well in 6-well plates (Corning) and transfected with 0.25 μg of “driver” plasmid (pTMO2F3 or p5UThORF2) + 0.5 μg of “reporter” plasmid (pJBM560 and derivatives thereof) + 0.25 μg of pCEP/GFP using 3 μL FuGENE® HD (Promega); the following day, the transfection media was removed and fresh media was added to each well. Three days post-transfection, cells were grown in the presence of G418 (400μg/mL) to select for retrotransposition events; during selection, the old media was replaced with fresh media containing G418. After approximately 12 days of selection, cells were washed with 1x PBS, fixed, and then stained with crystal violet to visualize colonies. To control for transfection efficiency, cells from replicate plates were trypsinized 72 hours after transfection and subjected to flow cytometry analyses to monitor GFP expression using an Accuri™ C6 flow cytometer (BD Biosciences). The number of G418-resitant colonies for each experimental condition was determined by dividing the number of G418-resistant colonies enumerated in a single well of a 6-well plate by the normalized GFP expression (normalized GFP expression = measured %GFP expression v*alu*e for each experimental condition divided by the highest %GFP expression v*alu*e). When indicated, ORF1*mneoI* retrotransposition assays were conducted in T-75 flasks or 100mm plates (Corning). To do that, approximately 1×10^6^ cells were plated and transfected with 1 μg of “driver” plasmid (pTMO2F3, p5UThORF2 or other) + 3 μg of “reporter” plasmid (pJBM560) using 12 μL FuGENE® 6 (Promega). Transfection efficiency determination, G418 selection, fixation, and foci staining were carried out as described above.

#### Next Generation DNA Sequencing

Genomic DNA was isolated from HeLa strains using a Qiagen DNeasy Blood and Tissue kit and then submitted to the University of Michigan DNA Sequencing Core facility for library preparation and Illumina sequencing. Briefly, HeLa genomic DNA was sheared to ∼320 bp and subjected to 2 x 150 paired-end sequencing on an Illumina HiSeq 4000, which yielded approximately 8-15X coverage for each HeLa strain. Copy number profiles were created using fastCN (49). Profiles were normalized to have a genome-wide average of 2 and thus represent relative copy numbers across the genome.

#### HeLa Karyotype Analyses

Karyotype analyses were performed as described previously (50), blindly, and using conventional protocols (51–53). We analyzed a total of twenty-five metaphase spreads for HeLa-HA and HeLa-JVM cell lines using the MetaSystem software (Izasa, Spain).

#### HeLa Spectral Karyotyping (SKY) Analyses

SKY-FISH analyses were performed as described previously (50). To identify markers, we used our karyotype analyses and data from a study that cytogenetically characterized HeLa CCL2 cells from ATCC (54). Briefly, we analyzed 21 and 16 metaphase spreads from HeLa-JVM and HeLa-HA cells, respectively, using an Applied Spectral Imaging kit (Applied Spectral Imaging, Migdal Ha′Emek, Israel), according to the protocol provided by the manufacturer. To capture images, we used an SD300 Spectra Cube (Applied Spectral Imaging), and a Zeiss Axioplan microscope equipped with a custom-designed optical filter (SKY-1; Chroma Technology, Brattleboro, VT). Because SKY is somewhat limited in the determination of breakpoints and in the identification of intrachromosomal changes (duplications, deletions, and inversions), breakpoints on SKY-painted chromosomes were determined by comparing DAPI banding with G-banding karyotype of both HeLa sublines.

#### HLA Genomic Typing

Typing analyses were conducted blindly using genomic DNA isolated from HeLa-HA and HeLa-JVM cells. Genomic DNA was isolated using a DNeasy Blood & Tissue Kit (Qiagen), following the manufacturer′s instructions. Next, we determined the HLA-genomic type of both cell lines using the Dynal kit for HLA-A*, B*, Cw*, DR*B1*\* and DQ*B1*\* (Dynal). As described (55), HLA alleles were assigned using the PMP software.

#### Microsatellite Marker Analysis

We analyzed Short Tandem Repeats (STRs) in chromosomes 6 and 15 using a Multiplex PCR blind strategy and genomic DNA isolated from HeLa-JVM and HeLa-HA strains (using a DNeasy Blood & Tissue Kit (Qiagen) and following the manufacturer′s instructions). We selected 8 primer pairs for the analysis of STRs located in chromosome 6 (7 in 6p21 and 1 in 6q21) and 5 markers for chromosome 15 (primer sequences in Sup Table 1). We then used a blind Multiplex PCR strategy by combining several markers, either four or five, using different staining and 1-4 pmol of each primer (forward and reverse, see Sup Table 1). The following combinations were used: Mx1: D6S276-311-291 and C.1.2.C; Mx2: D6S105-265-273 and C.1.2.5; Mx3: D15S146-126-209-1028-153. Amplicon size, labeling and sequence of the primers are shown in Sup Table 1. PCR reactions were carried out in 15 µl using the Taq DNA Polymerase kit (Roche) and the following conditions: 50 ng gDNA; 1 μl Primer Mix (5 μM each primer); 1.50 μl 10x PCR reaction buffer including 15 mM MgCl_2_ (Roche); 1.50 μl dNTPs mix (250 μM each dNTP); 0.12 μl Taq DNA polymerase (5 U/μl, Roche) and DNA-free distilled deionized water up to 15 μl. We used a PTC-100 Cycler (MJ Research, Inc.) and the following amplification program: 95°C, 12 minutes; 10 cycles of (94°C, 30 seconds; 55°C, 30 seconds; 72°C, 30 seconds); 20 cycles of (89°C, 30 seconds; 55°C, 30 seconds; 72°C, 30 seconds); and 60°C for 45 minutes. PCR products were diluted (1:3) using loading buffer (ThermoFisher), heat-denatured for 3 minutes at 95°C, and analyzed using an automatic sequencer ABI PRISM 3130xl Genetic analyzer (PE Applied Biosystem). Data analysis and the assignment of the size of each allele were carried out using ABI PRISM GeneMapper^TM^ (PE Applied Biosystem).

#### Immunoprecipitation and L1 Element Amplification Protocol (LEAP)

The LEAP assay has been described previously (56,57). Briefly, approximately 6×10^6^ HeLa cells were seeded onto 15 cm dishes (Corning); the next day, cells were transfected with a total of 10 μg plasmid DNA using 30 μL FuGENE HD (Promega). The following day, the transfection media was removed, and fresh media was added to each tissue culture dish. Approximately 48 hours after transfection, cells were washed with 1x PBS and scraped into 15 mL centrifuge tubes, centrifuged at 1000×g for 5 minutes, and then cell pellets were frozen at -80°C. Cell pellets then were thawed and lysed (1 mL of lysis buffer per ∼300 mg cell pellet) in lysis buffer (150 mM KCl, 2.5 mM MgCl_2_, 20 mM Tris-HCl [pH 7.5], 1 mM DTT, 0.1% NP-40, and 1× protease inhibitor EDTA-free mixture [Roche]) on ice for 30 minutes. Whole cell lysates then were centrifuged at 12,000×g for 15 minutes at 4°C. Next, 0.5 mL of the cleared lysate (∼2.5 mg total protein) was combined with anti-Flag (Sigma, 1804) coated Dynabeads Protein G (Invitrogen 10003D) and incubated with rotation for 3 hours at 4°C to precipitate 3XFLAG tagged *L1 ORF2p*. Beads were then washed four times with cold lysis buffer and immune complexes were eluted with 75 μL of 3XFLAG peptide (Sigma) (200 mg/mL) diluted in cold lysis buffer with rotation for 1 hour at 4°C. The eluates were then aliquoted into 1.5 mL microcentrifuge tubes, flash frozen in a dry ice/ethanol bath, and stored at -80°C. For the LEAP reaction, 1-3 μL of 3XFLAG eluate was combined and incubated with the LEAP reaction mixture [50 mM Tris-HCl [pH 7.5], 50 mM KCl, 5 mM MgCl2, 10 mM DTT, 20 U RNasin [Promega], and 0.05% Tween-20) containing HPLC-purified RACE12T Primer (sense: 5′-GCGAGCACAGAATTAATACGACTGGTTTTTTTTTTTT-3′) (0.4 μM) and dNTPs (0.2 mM)] at 37°C for 1 hour. One microliter of the LEAP reaction then was amplified in a 25 μL PCR reaction using Q5 Hot Start High-Fidelity 2X Master Mix (NEB, M0494S) with primers specific for sequences within the SV40 promoter of the *neo*^Tet^ and *mneoI* reporter cassettes (NeoSV40_fwd: 5′-GGGGCGGGACTATGGTTG-3′) and the 5′ end of the LEAP primer (3′LEAPOuter: 5′-GCGAGCACAGAATTAATACGACT-3′). Thermal cycler conditions: an initial cycle of 98°C for 30 seconds, followed by 35-40 cycles of 10 seconds at 98°C, 30 seconds at 63°C, and 8 seconds at 72°C with a final cycle of 72°C for 2 minutes. LEAP reaction products (10-20 μL) were visualized on 2% agarose gels using Ethidium bromide.

#### Cloning and Sequencing of LEAP Products

Visible LEAP bands were excised from agarose gels and cDNAs were purified using the Monarch DNA Gel Extraction Kit (NEB). Purified DNAs from excised gel slices were cloned into a pCR4Blunt-TOPO vector using a Zero Blunt TOPO PCR Cloning Kit for Sequencing (ThermoFisher Scientific) and subjected to Sanger DNA sequencing using M13 forward and M13 reverse primers (Eurofins Genomics).

#### M-MLV RT-PCR

RNA was purified from 30 μL of anti-FLAG IP eluates using a RNeasy Micro RNA extraction Kit (Qiagen, 74004) with on column DNaseI treatment and the RNA was eluted using 14 μL of DNA-free distilled deionized H_2_O. Next, first strand cDNA synthesis was performed using a SuperScript III First Strand Synthesis kit (Invitrogen, 18080051) using the RACE12T primer (50 μM). The cDNA reactions were diluted with H_2_O (1:5) and 2 μL of the diluted cDNA reaction was used in PCR reactions (25 μL) using the Q5 Hot Start High-Fidelity 2X Master Mix (NEB, M0494S) with NeoSV40_fwd and 3′LEAPOuter primers. Thermal cycler conditions: an initial cycle of 98°C for 30 seconds, followed by 35 cycles of 10 seconds at 98°C, 30 seconds at 63°C, and 10 seconds at 72°C with a final cycle of 72°C for 2 minutes. M-MLV RT-PCR reaction products (10-20 μL) were visualized on 2% agarose gels using ethidium bromide.

#### Western Blotting and Primary Antibodies

Standard western blotting procedures were used for protein analysis. Blots were analyzed using an Odyssey CLx (LI-COR) and western blot quantification was performed using the Image Studio software (version 3.1.4, LI-COR). The αhrGFP (240241; 1:5,000 dilution) was obtained from Agilent. αS6 (2217; 1:1,000 dilution) was obtained from Cell Signaling Technology. The αSRP9 (66068; 1:1,000 dilution) and αSRP14 (11525; 1:1,000 dilution) antibodies were obtained from Proteintech. The αFLAG (3165; 1:10,000 dilution), and αTubulin (T9026; 1:10,000 dilution) antibodies were obtained from Sigma.

#### Characterization of *Alu* Insertions by Inverse PCR

Inverse PCR was performed as previously described (26,58,59). Briefly, 2×10^6^ HeLa-JVM cells were plated in 100 mm plates, and 2×10^4^ HeLa-HA cells were plated into single wells of a six well plate. After ∼7 hours, HeLa-JVM cells were co-transfected with 30 μL of FuGENE 6 (Promega) and 7.5 μg pA2 plasmid DNA and 2.5 μg pJBM561 plasmid DNA per 100 mm plate; HeLa-HA cells were transfected with 3 μL of FuGENE 6 (Promega) and 1 μg pA2 plasmid DNA per well of a six well plate. The following day, transfection media was removed, and fresh media was added to each plate or well. Three days post-transfection, cells were subjected to G418 selection (400 μg/mL). During selection, the old media was replaced with fresh media containing G418 as needed. After 12 days, individual G418^R^ colonies were isolated and expanded. Genomic DNA was isolated from expanded G418^R^ foci using a DNeasy Blood & Tissue Kit (Qiagen). Five micrograms of genomic DNA derived from G418^R^ colonies were digested to completion with *Ssp*I or *Apo*I (NEB) in a total reaction volume of 50 μL. The restricted DNAs were ligated using 8 μL (3200U) of T4 DNA ligase (NEB) in a volume of 600 μL at 16°C overnight. The ligated DNA was precipitated with ethanol, washed, and dissolved in 30 μL of DNA-free distilled deionized H_2_O. Five microliters of ligated DNAs were used in the primary PCR reaction in a 50 μL reaction volume containing 125 μM of each dNTP, 300 nM of primers (NEO210AS: 5′-GACCGCTTCCTCGTGCTTTACG-3′) and (NEO742S: 5′-TCATTCAGGGCACCGGACAG-3′), 1× buffer 3, and 5 U of enzyme mix using the Expand Long Template PCR system (Roche). Samples were amplified with the following thermocycler conditions: one cycle of 95°C for 2 minutes, then 30 cycles of 94°C for 10 seconds, 65°C for 30 seconds, and 68°C for 15 minutes, followed by a final extension step at 68°C for 30 minutes. The resultant products (5 μL) were used in a second PCR reaction using the same conditions with nested primers (NEO148AS: 5′-CGAGTTCTTCTGAGGGGATCGG-3′) and (NEO1808S: 5′-GCGTGCAATCCATCTTGTTCAATG-3′). PCR products from the second amplification were purified using a gel extraction kit (Qiagen). The purified fragments were cloned into pGEM-T easy (Promega) and sequenced at the University of Michigan DNA Sequencing Core facility. The *Alu* insertion structure was inferred directly from Sanger sequence information (*i.e.,* distal 5′ and 3′ ends, Target Site Duplications (TSDs), poly(A) sequences, flanking genomic DNA sequences, *etc*.); the sequences flanking the *Alu* insertions were used as probes in BLAT searches at the UCSC genome browser (http://genome.ucsc.edu, assembly GRCh37/hg19), to determine the insertion site location (60).

## RESULTS

### HeLa strains differ in their ability to support *Alu* retrotransposition

To monitor *Alu* retrotransposition, we used a previously described cultured cell retrotransposition assay (4) (Fig 1A). Briefly, we co-transfected HeLa cells with a “driver” plasmid (pTMO2F3) that expresses a version of human *L1 ORF2p* containing a carboxyl-terminal 3X-FLAG epitope tag and an *Alu* “reporter” plasmid (p*Aluneo*^Tet^) that expresses an active human *Alu*Y element tagged in its 3′ end with a retrotransposition indicator cassette (*neo*^Tet^) (4,34). *Alu* expression is augmented by a 66 bp human *7SL RNA* gene enhancer, which is located directly upstream of the first *Alu* nucleotide. The *neo*^Tet^ indicator cassette is designed to be compatible with RNA pol III expression (34) and consists of a backward copy of the neomycin phosphotransferase gene, which is interrupted by a *Tetrahymena* self-splicing group I intron that is in the same transcriptional orientation as the *Alu* sequence. A 44 bp poly(A) tract is located after the *neo*^Tet^ indicator cassette, which is immediately followed by an RNA pol III terminator sequence (Fig 1A). An SV40 promoter sequence drives expression of the *neo* gene after it is integrated into genomic DNA by retrotransposition, thereby conferring G418-resistance to cells. Thus, the number of G418-resistant foci is a quantitative indicator of the efficiency of *Alu* retrotransposition in cultured cells.

**Figure 1:**
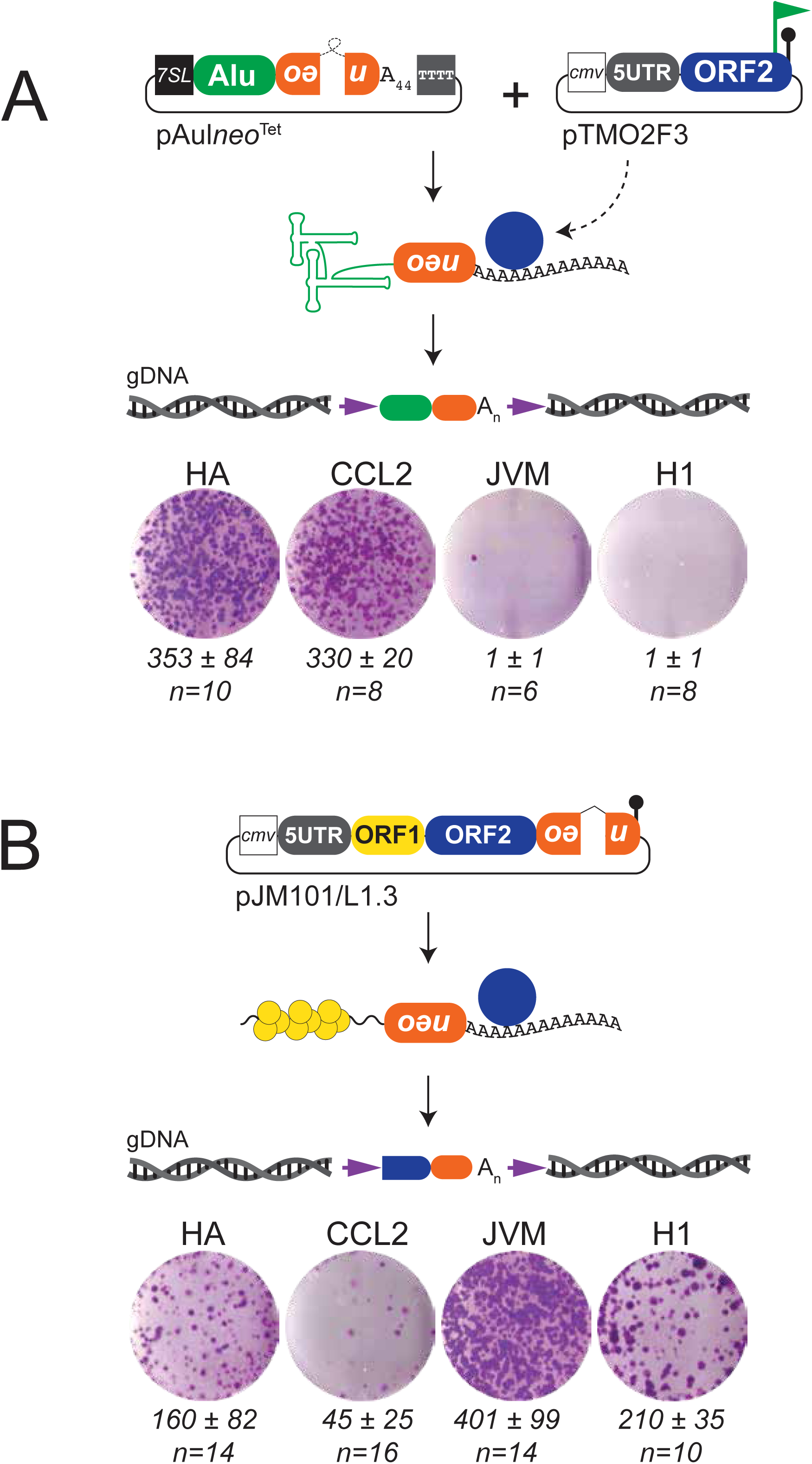
HeLa strains differ in their capacity to support *Alu* retrotransposition. *(A) Top Panel: Schematic of the Alu retrotransposition assay.* HeLa cells were co-transfected with pTMO2F3 and p*Aluneo*^Tet^. pTMO2F3 expresses ORF2p with a carboxyl terminus 3XFLAG tag (green flag). ORF2p expression is augmented by a CMV promoter (white square), the native L1 5′ UTR (grey oval), and an SV40 polyadenylation signal sequence (black lollipop). The p*Aluneo*^Tet^ retrotransposition marker (*neo*^Tet^; orange oval) contains a backwards copy of the neomycin phosphotransferase gene interrupted by a self-splicing group I intron (loop) that is in the same transcriptional orientation as the *Alu* sequence. A 44 bp poly(A) tract follows the *neo*^Tet^ sequence. *Aluneo*^Tet^ expression is augmented by a 7SL gene enhancer (black square) and a sequence of four consecutive thymidine residues (grey square) located downstream of the 44 bp poly(A) sequence. The neomycin phosphotransferase gene can only be expressed when the *Alu* transcript is spliced and subsequently is inserted into genomic DNA by L1 ORF2p (Blue circle). The resultant insertions are flanked by target site duplications (purple arrows). *Bottom Panel: Results of Alu retrotransposition assays*. Displayed are single wells of a representative six-well tissue culture plate from *Alu* retrotransposition assays. The HeLa strain is denoted above each image. Below each image is the average number of G418-resistant colonies per well ± standard deviation (n=number independent transfections). *(B) Top panel: Schematic of the L1 retrotransposition assay*. HeLa cells were transfected with an engineered human L1.3 construct (pJM101/L1.3) marked with a *mneoI* retrotransposition indicator cassette (orange oval) that consists of a backwards copy of the neomycin phosphotransferase gene interrupted by the gamma globin intron 2 sequence (inverted “V”) that is in the same transcriptional orientation as the L1 sequence. A CMV promoter (white square) and an SV40 polyadenylation signal sequence (black lollipop) augment the expression of pJM101/L1.3. The neomycin phosphotransferase gene can only be expressed when the L1 transcript is spliced, the L1 proteins (ORF1p [yellow circle] and ORF2p [blue circle]) associate with their encoding L1 RNA in *cis*, and the L1 RNA is inserted into genomic DNA. The resultant retrotransposition insertions are flanked by target site duplications (purple arrows). *Bottom panel: Results of L1 retrotransposition assays*. Displayed are single wells of a representative six-well tissue culture plate from L1 retrotransposition assays. The HeLa strain is denoted above each image. Below each image is the average number of G418-resistant colonies per well ± standard deviation (n=number independent transfections).

We initially screened for L1 and *Alu* retrotransposition in a panel of HeLa strains (HeLa-HA, HeLa-CCL2, HeLa-JVM and HeLa-H1; see Methods). We found that human *Alu*Y (p*Aluneo*^Tet^) retrotransposes efficiently in HeLa-HA and HeLa-CCL2 (herein called *Alu*-permissive strains), but not in HeLa-JVM or HeLa-H1 (herein called *Alu*-nonpermissive strains) (Fig 1A). By comparison a retrotransposition-competent human L1 (L1.3) could retrotranspose in all four HeLa strains (Fig 1B, see below). Additional controls demonstrated that despite using different cell plating densities, transfection reagents, “driver”/“reporter” plasmid ratios, concentrations of G418 during selection, and cell culture media, *Alu*Y retrotransposition is restricted to the HeLa-HA and HeLa-CCL2 strains and is greatly enhanced by the co-expression of a “driver” construct expressing a functional version of *L1 ORF2p* (data not shown).

Subsequent experiments further revealed: (**1**) an engineered *Alu*Ya5 “reporter” could retrotranspose in the HeLa-HA strain when it was included in the same plasmid backbone as an *L1 ORF2* “driver” gene (Fig S1A, plasmid pA2); (**2**) a mutagenic *Alu*Ya5 insertion isolated from the neurofibromatosis I (*NF1*) gene (6) could undergo retrotransposition in the HeLa-HA strain when a different RNA pol III terminator (BC1) signal was included in the plasmid (Fig S1B, pQ19-33-*Alu*Ya5); (**3**) PCR-based assays could detect evidence of robust *Alu* retrotransposition (*i.e.*, spliced neo gene products) in genomic DNA derived from HeLa-HA cells that were co-transfected with p*Aluneo*^Tet^ “reporter” and an L1 ORF2p “driver” (pTMO2F3), but markedly less *Alu* retrotransposition was observed in HeLa-JVM cells using the same assay (Figs S1C and S1D); (**4**) co-transfection of p*Aluneo*^Tet^ with a reverse transcriptase missense mutant version of pTMO2F3 (pTMO2F3^D702A^) did not give rise to G418-resistant colonies in either the HeLa-HA or HeLa-JVM strains (Fig S1E); and (**5**) co-transfection of p*Aluneo*^Tet^ with a full-length L1.3 “driver” plasmid that expresses both *L1 ORF1p* and *L1 ORF2p* (pTMF3Δ*neo*) promoted *Alu* retrotransposition in HeLa-HA, but not in HeLa-JVM cells (data not shown).

We next confirmed that *Alu*-permissive and *Alu*-nonpermissive strains are *bona fide* HeLa cell derivatives by conducting: (**1**) blinded low-resolution HLA-typing of Class I (HLA-A: A*03:19,68:02; HLA-B: B*15:03,15:03; HLA-C: C*12:03,12:03) and Class II (HLA-DQ: DQA1*01:02,01:02 and DQ*B1*\*05:01,05:01; HLA-DR: DR*B1*\*01:02,01:02) markers (61) (Fig S2A; see Methods); (**2**) blinded Short Tandem Repeat (STR) analyses of thirteen markers located on chromosomes 6 and 15 (62) (Fig S2B and Sup Table 1, [eight chromosome 6 and five chromosome 15 markers, respectively]); and (**3**) STR analyses through a commercial vendor (Fig S2C and Sup Table 2).

Spectral karyotyping (SKY) (54) and G-banding experiments (Figs 2A, 2B, 2C and Sup Table 3), further revealed that the HeLa-HA and HeLa-JVM strains exhibit near triploid (3n) karyotypes, with an average of 62 and 54 chromosomes per cell, respectively (Sup Table 3). Finally, 8X-15X low coverage Illumina whole genome sequencing and copy number (CN) analyses (Fig 2D, Sup Fig S2D; see Methods) showed that HeLa-HA and HeLa-CCL2 exhibited similar CN content across individual chromosomes that was distinct from that observed in the HeLa-JVM and HeLa-H1 strains (Fig 2D, Sup Fig S2D). Taken together, the above data suggest we identified two closely related *Alu*-permissive (HeLa-HA and HeLa-CCL2) and two closely related *Alu*-nonpermissive (HeLa-JVM, HeLa-H1) strains.

**Figure 2:**
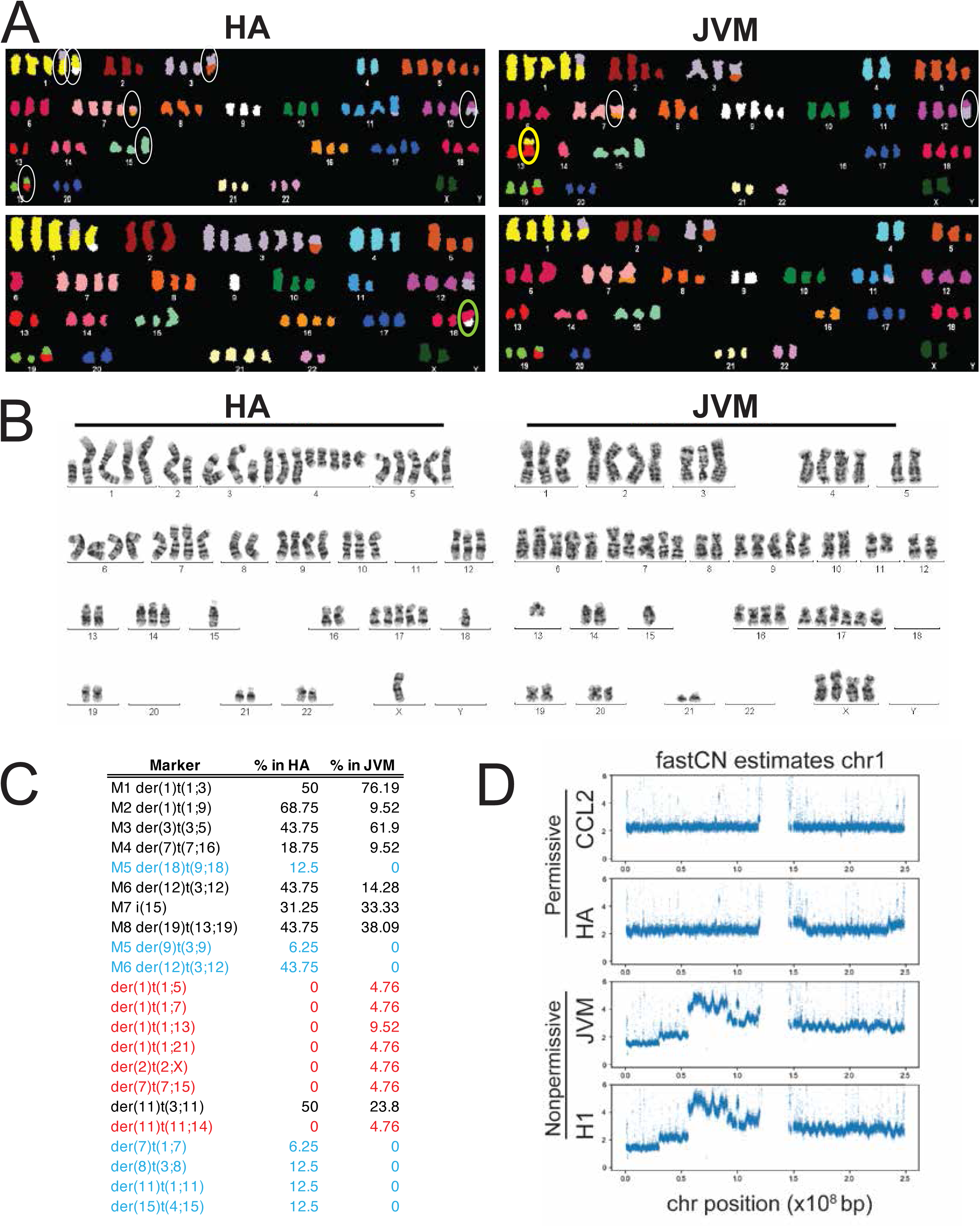
Genetic characterization of *Alu*-permissive and *Alu-*nonpermissive HeLa strains. *(A) SKY FISH.* Mitotic spreads showing representative cells from HeLa-HA (left panels) and HeLa-JVM (right panels). White ovals indicate examples of common marker chromosomes shared between HeLa-JVM and HeLa-HA. Some marker chromosomes were unique to HeLa-HA (green oval) or HeLa-JVM (yellow oval) (see also Fig 2C). *(B) Karyotypes.* Shown are two representative G-banding karyotypes of HeLa-HA (left) and HeLa-JVM (right) cells; markers are not included in the panel. Twenty-five metaphase spreads were scored for each strain (see Sup Table 3). (*C) Summary chart of HeLa cell SKY-FISH analyses.* Consistent with karyotype analyses, both HeLa-HA and HeLa-JVM share a common origin. There are several markers that have been characterized previously (54) that are common to both HeLa-HA and HeLa-JVM (*e.g.*, der (1)t(1;3)). Some markers were only observed in the HeLa-JVM (red text) or HeLa-HA strains (blue text). *(D) Copy number (CN) estimates for the four HeLa strains.* Graphs depict CN estimates for chromosome 1 of each HeLa cell strain (indicated to the left of each plot). The X-axis indicates chromosome position; the Y-axis indicates CN (also see Sup Fig S2D).

### *Alu*-permissive and *Alu*-nonpermissive HeLa strains support the retrotransposition of human and non-human LINEs

We next analyzed the retrotransposition efficiency of engineered LINEs in the *Alu*-permissive (HeLa-HA) and *Alu*-nonpermissive (HeLa-JVM) strains. Retrotransposition assays (22,45,46) revealed that a wild type human LINE-1 (L1.3 (32), plasmid pJM101/L1.3 (26)) retrotransposed efficiently in both HeLa strains (Fig 1B). Consistently, additional retrotransposition assays revealed that engineered versions of a natural mouse L1 element from the G_F_ subfamily (pTG_F_21) (44), a highly active synthetic codon-optimized mouse L1 from the T_F_ subfamily (pCEPsmL1) (40), and a natural zebrafish LINE-2 element (pZfL2-2*mneoI*) (43) retrotransposed at comparable levels in both the HeLa-HA and HeLa-JVM strains (Figs 3A and S3A). Controls revealed that a missense mutation in the L1.3 reverse transcriptase active site (pJM105/L1.3; D702A) (22,26) severely reduced L1 retrotransposition (Figs 3A, 3B, S3A, and S3B).

**Figure 3:**
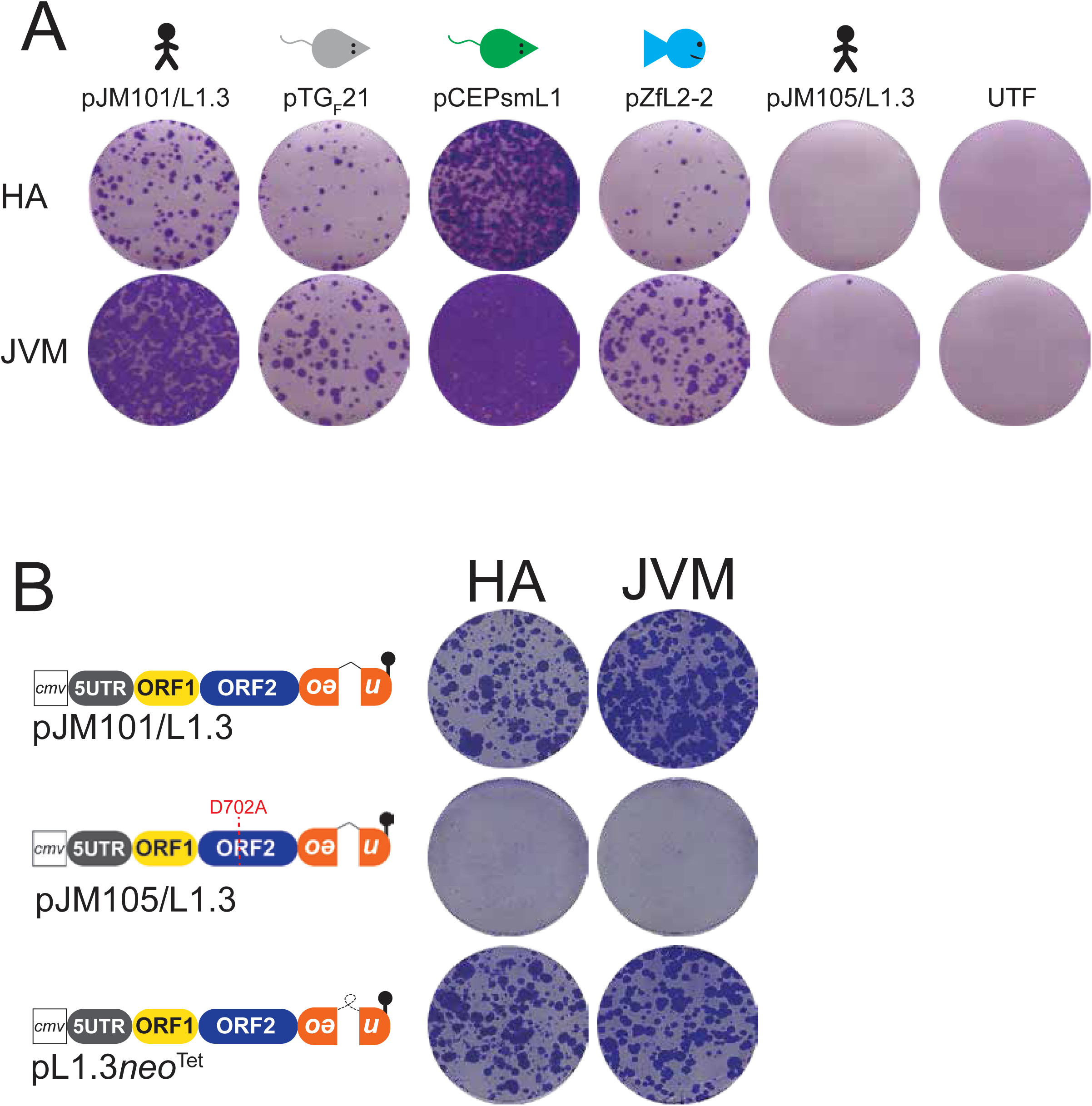
The retrotransposition of vertebrate LINEs in HeLa strains. *(A) HeLa strains support the retrotransposition of mouse and zebrafish LINEs*. HeLa cells were transfected in 6-well plates at ∼2×10^5^ cells/well with human L1 (pJM101/L1.3), mouse L1 (pTG_F_21), synthetic mouse L1 (pCEPsmL1), zebrafish LINE-2 (pZfL2-2), or a human L1 containing a D702A mutation in ORF2 (pJM105/L1.3). Displayed are single wells of a representative six-well tissue culture plate from qualitative retrotransposition assays. The experiments were repeated three times with similar results. The LINE plasmid is indicated above each image and the HeLa strain is indicated to the left of each row of images. *(B) Results of pL1.3neo^Tet^ retrotransposition assay*. HeLa cells were transfected in 6-well plates with pJM101/L1.3, pL1.3*neo*^Tet^ (which contains the *neo*^Tet^ reporter), or pJM105/L1.3. A CMV promoter, native L1 5′ UTR, and an SV40 polyadenylation signal sequence augment LINE expression. The right panel displays retrotransposition results from single wells of a representative six-well tissue culture plate. The HeLa strain is denoted above each column of images. The assay was done three times yielding similar results.

Our data revealed that engineered vertebrate LINEs can retrotranspose efficiently in both *Alu*-permissive and *Alu*-nonpermissive HeLa strains (Figs 3 and S3). However, the LINE cultured cell retrotransposition assay relies on the use of a spliceosomal intron within the *mneoI* retrotransposition indicator cassette (22,63) (Fig 1B), whereas the *Alu* retrotransposition assay relies on the use of an autocatalytic self-splicing group I intron within the *neo*^Tet^ retrotransposition indicator cassette (4,34) (Fig 1A). To examine whether the use of a self-splicing of group I intron might be responsible for lack of *Alu* retrotransposition in nonpermissive HeLa strains, we tagged a human L1.3 element with the *neo*^Tet^ cassette used in the *Alu* retrotransposition assay (Fig 3B, plasmid pL1.3*neo*^Tet^). We readily detected retrotransposition events derived from both pJM101/L1.3 and pL1.3*neo*^Tet^ in the HeLa-HA and HeLa-JVM strains (Figs 3B and S3B). Thus, the inability to detect *Alu* retrotransposition in the HeLa-JVM strain is not likely caused by misfunctioning of the *neo*^Tet^ indicator cassette.

### *Alu* insertions in the HeLa-HA and HeLa-JVM strains exhibit TPRT structural hallmarks

To confirm that G418-resistant foci result from *bona fide Alu* retrotransposition events, we transfected the HeLa-HA and HeLa-JVM strains with the pA2 vector (Fig. S1A) and selected for G418-resistant foci. Genomic DNA then was isolated from clonally expanded G418-resistant cells and was used as a template in inverse PCR reactions to characterize the *de novo Alu* integration events (see Methods). Because of the low level of *Alu* retrotransposition (*i.e.,* G418-resistant foci) detected in HeLa-JVM assays (see Figs 1A, S1A, and S1B), we used larger tissue culture plates (*i.e.*, 100 mm dishes) to conduct analyses in the HeLa-JVM strain.

The analysis of 26 *de novo Alu*Ya5 insertions from the HeLa-HA strain (Figs 4A and 4B; Sup Table 4) revealed typical TPRT hallmarks (*i.e*., the insertions ended in a 3′ poly(A) tract, were flanked by variable length target site duplications [TSDs], and generally integrated into an L1 ORF2p endonuclease consensus cleavage sequence [5′-TTTTT/AA-3′, where the “/” indicates the cleavage site]) (64,65). The *Alu*Ya5 insertions were of full-length and ended in 3′ poly(A) tracts that averaged 71 nts in size (ranging from 26-113 nts) and were flanked by TSDs that ranged from 7-18 bp (Sup Table 4). One HeLa-HA insertion (HA-27) lacked the first *Alu* “G” nucleotide, whereas another insertion (HA-41) contained an untemplated “CA” dinucleotide insertion that preceded the first *Alu*Ya5 “G” nucleotide (Sup Table 4). Two insertions (HA-4 and HA-36) contained a single mismatch in the 5′ TSD (Sup Table 4). Notably, the mismatch in the TSD of HA-4 (*i.e.*, a C>T change) is consistent with previously reported APOBEC3A mutational signatures of single stranded genomic DNA at L1 integration sites (66). The *Alu*Ya5 insertions in the HeLa-HA strain also were dispersed across the genome at different chromosomal locations (Fig 4C).

**Figure 4:**
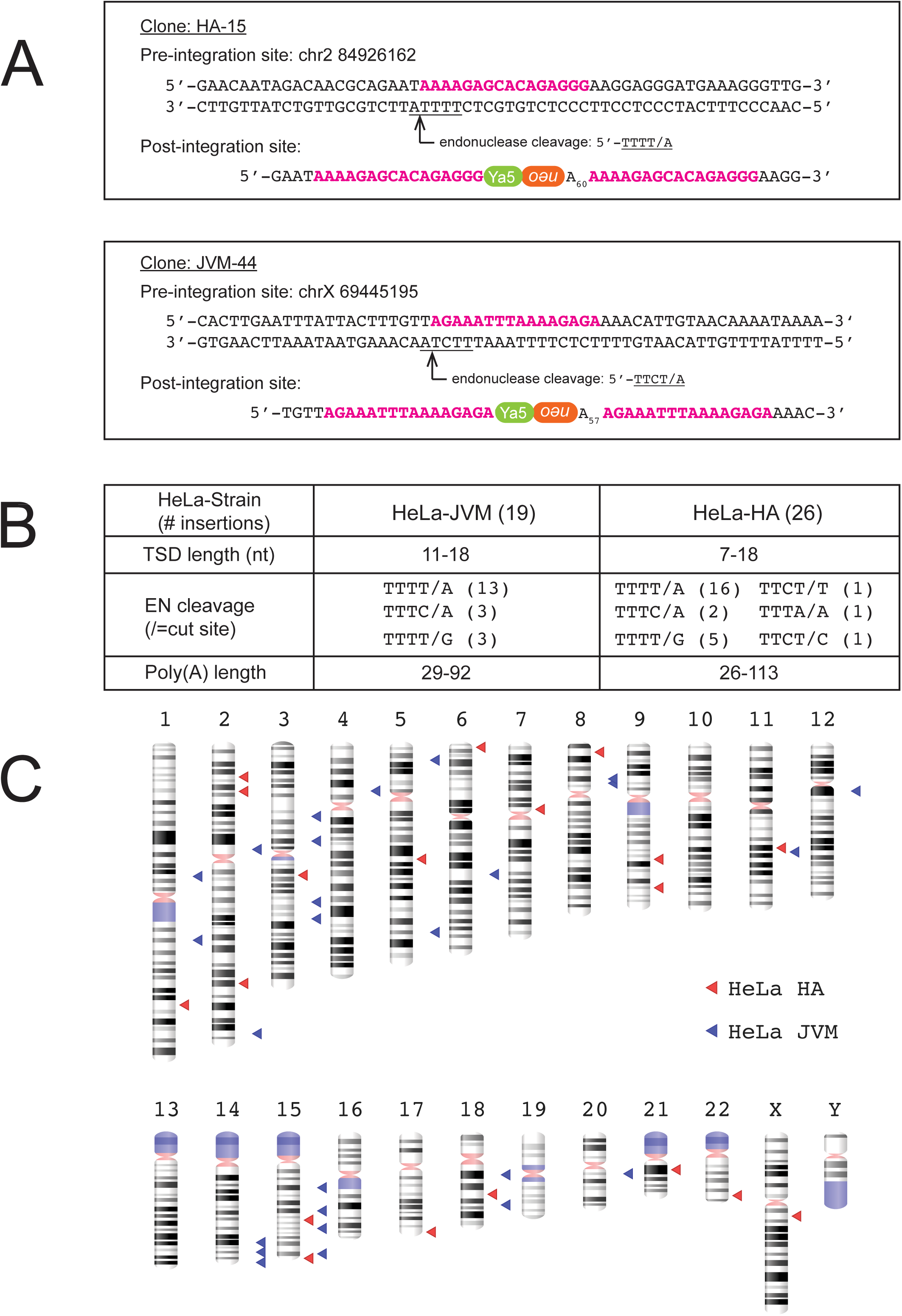
Structure of *Alu* insertions in HeLa cells. *(A) Structure of pA2 insertions.* Representative structure of *de novo Alu*Ya5 insertions identified in HeLa-HA (Clone 15, top), and HeLa-JVM (Clone-44, bottom) cells. Genomic insertion site locations are based on human reference genome sequence GRCh37/hg19. The *Alu*Ya5 insertions contain typical TPRT structural hallmarks indicative of L1-mediated retrotransposition (see main text). Target site duplications are noted in bold magenta characters. The L1 ORF2p EN target site is underlined; the arrow and “/” indicates the cleavage site. *(B) Summary table of Alu events characterized in HeLa-HA and HeLa-JVM strains.* Column 1: TPRT-mediated structural hallmarks; Columns 2 and 3: number of insertions characterized from the HeLa-JVM (column 2) and HeLa-HA (column 3) strains, respectively. Row 1, total number of insertions characterized in HeLa-JVM and HeLa-HA cells. Row 2, ranges of TSD length. Row 3, L1 ORF2p EN cleavage sequences; Row 4, estimated poly(A) tail lengths in nucleotides. *(C)* Ideograms depicting the approximate genomic locations (GRCh37/hg19) of *Alu*Ya5 insertions in HeLa-JVM (blue triangles) and HeLa-HA (red triangles).

The analysis of 19 *de novo Alu*Ya5 insertions from the HeLa-JVM strain (Figs 4A and 4B; Sup Table 5) also revealed typical TPRT hallmarks (see above). The *Alu*Ya5 insertions were of full-length and ended in 3′ poly(A) tracts that averaged 62 nts in size (ranging from 29-92 nts) and were flanked by TSDs that ranged from 11-18 bp. One insertion (JVM-21) included 20 nts of reporter plasmid sequence preceding the first *Alu*Ya5 nucleotide, which could have resulted from transcription initiating upstream of the *Alu*Ya5 sequence, as well as an untemplated “G” residue at the 5′ end of the insertion. Another insertion (JVM-57) contained a 1 bp deletion of the first *Alu*Ya5 “G” nucleotide (Sup Table 5). The *Alu*Ya5 insertions in HeLa-JVM were dispersed across the genome at different chromosomal locations (Fig 4C). Thus, the structures of the *Alu*Ya5 insertions in the HeLa-HA and HeLa-JVM strains represent *bona fide* L1-mediated retrotransposition events. These data indicate that the differences in *Alu* retrotransposition efficiency between HeLa strains appear to be due to a reduction in *Alu* insertion frequency.

### HeLa strains differ in their ability to support the retrotransposition of older resurrected *Alu* elements and other *7SL RNA*-derived SINE RNAs

To further define the retrotransposition defect in *Alu-*nonpermissive HeLa strains, we next tested whether *L1 ORF2p* could drive the retrotransposition of consensus sequences derived from an older *Alu* subfamily element (*Alu*Sx3) (67), an ancient consensus *Alu*J left-hand monomer sequence (p*Alu*J) (12), or a consensus *BC200* RNA, which is a non-coding RNA expressed in the brains of primates that arose from an ancient monomeric *Alu* element (37,68). Each *Alu* element, as well as the consensus *BC200* sequence (*BC200*con1 (37)), was tagged with the *neo*^Tet^ indicator cassette and assayed for their ability to undergo L1 ORF2p-dependent retrotransposition in the HeLa-HA and HeLa-JVM strains (Figs 5A, 5B, and 5C; see Methods). Consistent with our analyses of evolutionarily younger *Alu*Y elements (see Fig 1), the consensus *Alu*Sx3 and *Alu*J elements, and the consensus *BC200* RNA, efficiently retrotransposed in the HeLa-HA, but not the HeLa-JVM strain (Figs 5A, 5B, and 5C).

**Figure 5:**
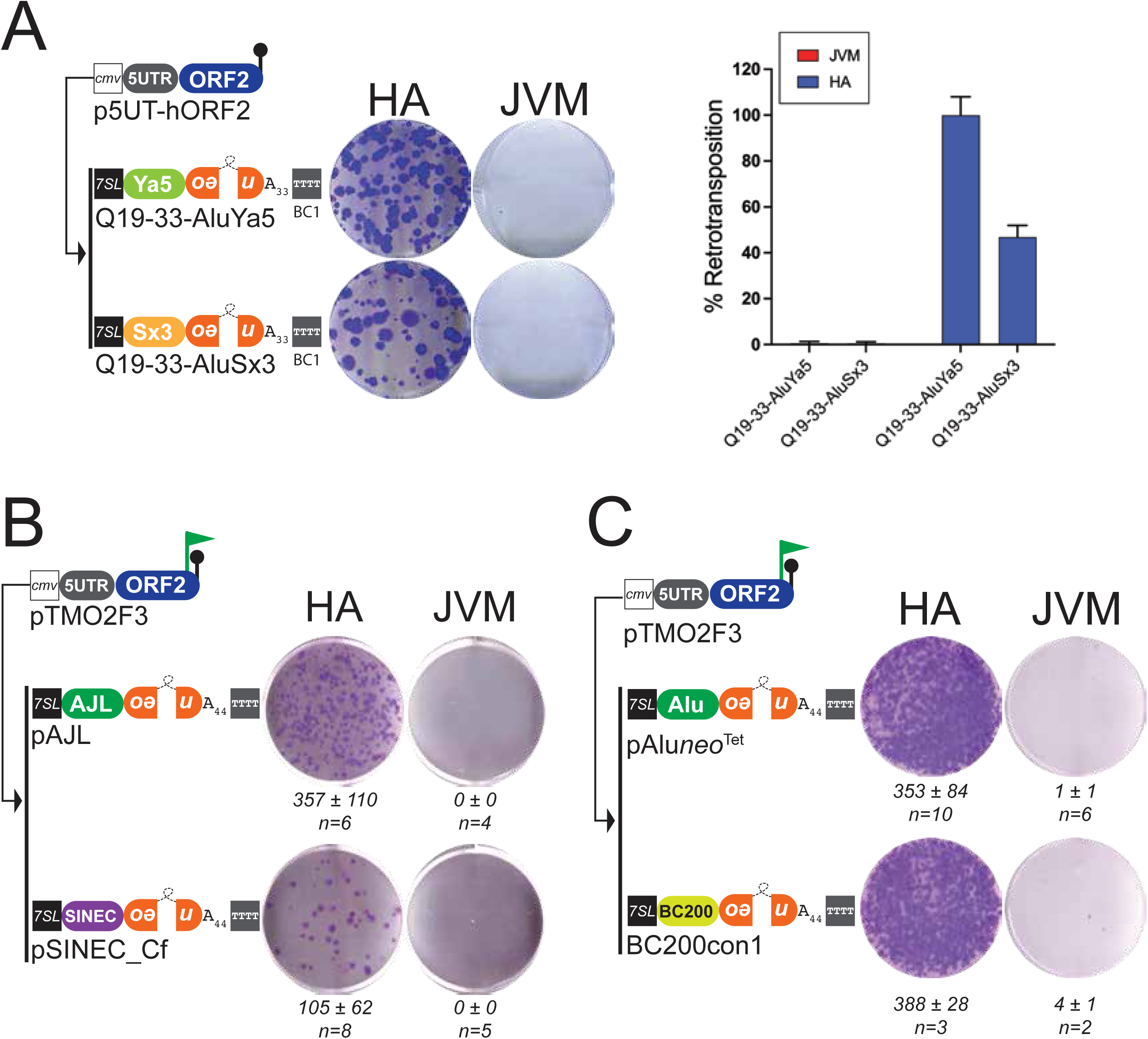
The retrotransposition of *7SL*- and *tRNA*-derived SINES in different HeLa strains. *(A) Retrotransposition results for AluSx3*. *Right panel:* HeLa cells were co-transfected with p5UT-hORF2 (see Fig 1D) and Q19-33-*Alu*Ya5 or Q19-33-*Alu*Sx3, which express an *Alu*Sx3 element (see main text). Displayed are single wells of a representative six-well tissue culture plate from qualitative retrotransposition assays. The HeLa strain is denoted above each column of images. The experiment was done three times with similar results. Left panel: quantitation of retrotransposition results. The X-axis indicates the *Alu* element co-transfected with p5UT-hORF2 and the Y-axis indicates the percent (%) retrotransposition efficiency normalized to Q19-33-*Alu*Ya5. Error bars indicate standard deviations. *(B) Retrotransposition results for AluJ and SINEC_Cf*. HeLa cells were co-transfected with pTMO2F3 (see Fig 1) and either pAJL (*Alu*J left hand monomer) or p*SINEC_Cf* (canine consensus *tRNA*-derived SINE). Displayed are single wells of a representative six-well tissue culture plate from the retrotransposition assays. The HeLa strain is denoted above each column of images. Below each well is the average number of G418-resistant colonies per well ± standard deviation (n=number independent transfections). *(C) BC200 retrotransposes in HeLa-HA*. HeLa cells were co-transfected with pTMO2F3 (see Fig 1) and either p*Aluneo*^Tet^ or *BC200*con1 (*BC200* consensus sequence (37)). Displayed are single wells of a representative six-well tissue culture plate from the retrotransposition assays. The HeLa strain is denoted above each column of images. Below each well is the average number of G418-resistant colonies per well ± standard deviation (n=number independent transfections).

We next tested whether another mammalian *7SL RNA*-derived SINE could undergo retrotransposition in an *Alu-*nonpermissive HeLa strain. Mouse *B1* elements comprise ∼2.7% of mouse genomic DNA and are the most abundant SINE in the mouse genome (69). They are derived from the *7SL RNA* gene but have a monomeric structure (70,71). Using the same co-transfection approach described above (see Fig 1A), we demonstrated that a recent mutagenic *7SL RNA*-derived mouse *B1* SINE (*B1*_Mur4) (36) could undergo efficient retrotransposition in the HeLa HA, but not the HeLa-JVM, strain (Figs S4A,and S4B; also see ref. (27)). Thus, the above data suggest that the *Alu* retrotransposition defect in *Alu-*nonpermissive HeLa cell lines probably extends to all primate *Alu* subfamilies and other *7SL-RNA* derived SINEs.

### HeLa strains differ in their ability to support the retrotransposition of *tRNA*-derived SINEs

In addition to *7SL RNA*-derived SINEs, transfer RNA (*tRNA*)-derived SINEs also are highly abundant in mammalian genomes. For example, *tRNA*-derived mouse *B2* elements (72,73) are present at approximately 350,000 copies and comprise ∼2.4% of mouse genomic DNA (69). Similarly, *tRNA*-derived SINEs (*SINE-C* sequences) are abundant in canine genomes and are highly polymorphic among different dog breeds (28).

To determine whether the retrotransposition defect in *Alu-*nonpermissive HeLa cell lines extends to other SINEs, we next tested whether L1 ORF2p could drive the retrotransposition of consensus sequences derived from either a canine *tRNA*-derived SINE (*pSINEC_Cf*) (28) or mouse *tRNA*-derived SINE (*B2* element; *B2*_Mm1a). Retrotransposition assays revealed that the canine *tRNA*-derived SINE and the mouse *B2* element efficiently retrotranspose in the HeLa-HA, but not the HeLa-JVM strain (Figs 5B, S4A, and S4B; also see ref. (27)). Thus, *7SL-* and *tRNA*-derived SINEs exhibit similar retrotransposition dynamics in *Alu-*permissive and *Alu-*nonpermissive HeLa strains, suggesting that they retrotranspose by similar molecular mechanisms.

### HeLa strains support the retrotransposition of an engineered *SVA* element

*SVA* elements are hominid specific non-autonomous non-LTR retrotransposons (74). Recent estimates indicate that an average human genome contains >5,000 *SVAs* (comprising ∼0.5% of human genomic DNA) and can be subclassified into six different subfamilies (*SVA*_A to *SVA*_F) (75). In contrast to mammalian *7SL RNA*- and *tRNA*-derived SINEs, *SVA* elements are longer in size (ranging from 700-4000 bp in length) (76), are transcribed by RNA pol II, and end in a post-transcriptionally added 3′ poly(A) tail (74). Moreover, whereas *Alu* retrotransposition only requires L1 ORF2p (4), previous reports suggested that *SVA* retrotransposition is dependent on the expression of both the *L1 ORF1*-encoded protein (ORF1p) and L1 ORF2p (29,30).

To determine whether the retrotransposition defect in *Alu-*nonpermissive HeLa strains extends to *SVA* elements, we co-transfected the HeLa-HA and HeLa-JVM strains with a full-length L1.3 “driver” plasmid that expresses both L1 ORF1p and L1 ORF2p (pTMF3Δ*neo*) and a “reporter” plasmid that expresses an active human *SVA_E* subfamily element tagged with a modified version of the *mneoI* retrotransposition reporter that prevents mis-splicing between the *SVA* element and *neo* gene (38) (Fig 6A). Retrotransposition assays revealed that *SVA* is active in both the HeLa-HA and HeLa-JVM strains (Fig 6A). By comparison, *SVA* assays conducted in HeLa strains co-transfected with a “driver” plasmid that only expresses L1.3 ORF2p (pTMO2F3) and the pAD26 “reporter” plasmid revealed *SVA* retrotransposition was reduced by ∼75% in both HeLa-HA and HeLa-JVM cells when compared to retrotransposition assays conducted with the L1 ORF1-containing pTMF3Δ*neo* “driver” plasmid (Fig 6A). Thus, *SVA* retrotransposition is more efficient when both L1 ORF1p and ORF2p are ectopically over-expressed from a “driver” plasmid in *Alu-*permissive and *Alu-*nonpermissive HeLa cell lines.

**Figure 6:**
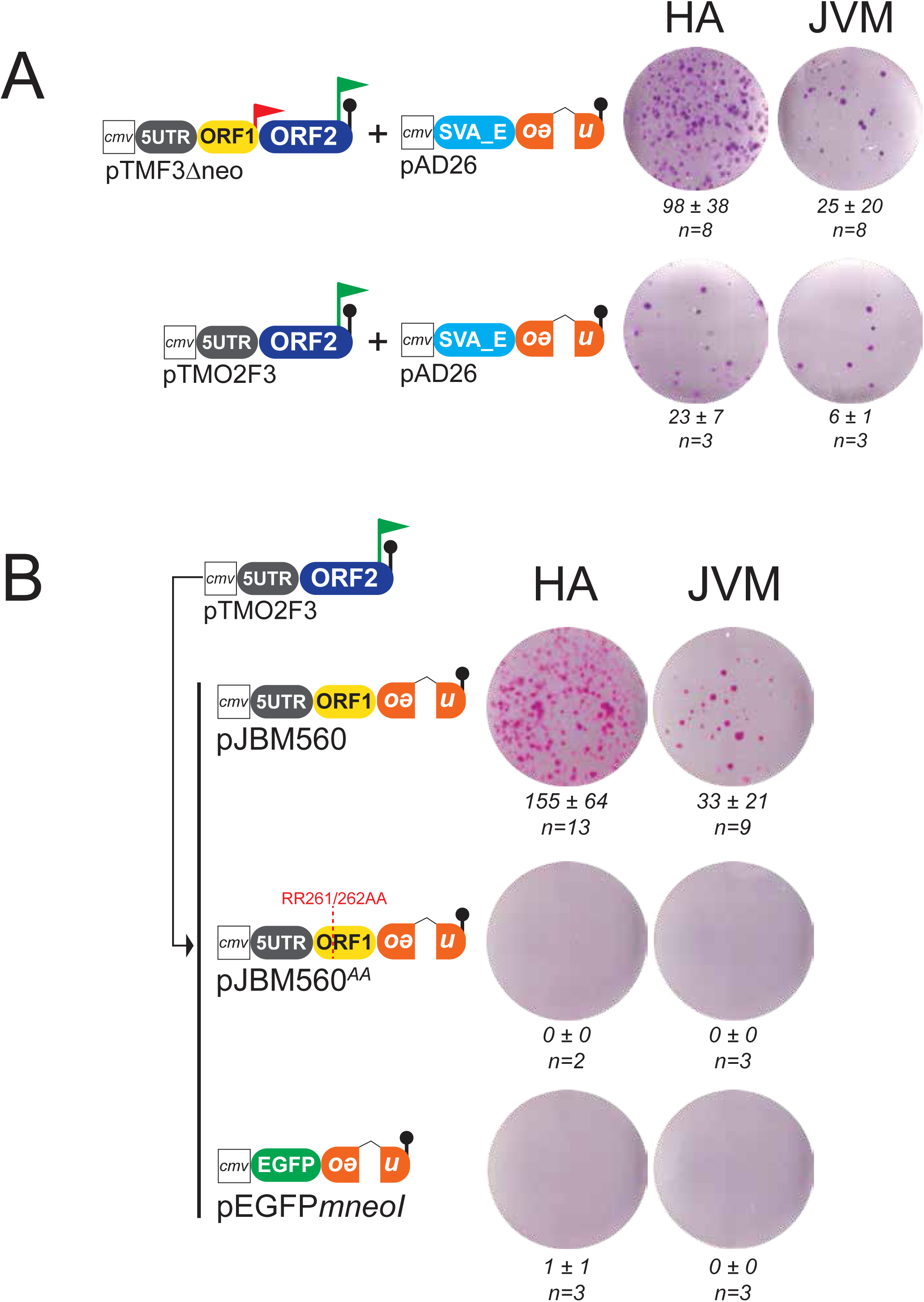
*SVA element* retrotransposition and processed pseudogene formation in different HeLa strains. *(A) SVA Retrotransposition results. Left panel:* HeLa cells were co-transfected with the “driver” plasmid (either pTMF3Δ*neo* or pTMO2F3) and the *SVA* retrotransposition “reporter” construct (pAD26). A CMV promoter and an SV40 polyadenylation signal sequence augment *SVA* expression. *Right panel:* Displayed are single wells of a representative six-well tissue culture plate from the *SVA* retrotransposition assays. The HeLa strain is denoted above each column of images. Below each well is the average number of G418-resistant colonies per well ± standard deviation (n=number independent transfections); *SVA* experiments were conducted at least three times. *(B) ORF1mneoI retrotransposition results.* HeLa cells were co-transfected with the “driver” plasmid pTMO2F3 and either pJBM560 (*L1 ORF1mneoI*), pJBM560^AA^ (a R261A and R262A missense mutation in *L1 ORF1*), or a processed pseudogene construct (p*EGFPmneoI*). A CMV promoter and an SV40 polyadenylation signal sequence augment expression of the “driver” and *mneoI*-tagged plasmids. Displayed are single wells of a representative six-well tissue culture plate from the retrotransposition assays. The HeLa strain is denoted above each column of images. Below each well is the average number of G418-resistant colonies per well ± standard deviation (n=number independent transfections).

### HeLa strains support the retrotransposition of functional *L1 ORF1* containing mRNAs

In addition to RNA polymerase III transcribed SINEs, the precursors of certain non-autonomous non-LTR retrotransposons are derived from RNA pol II transcripts that end in long 3′ poly(A) tails. For example, we previously demonstrated that L1 ORF2p could act *in trans* to mediate the retrotransposition of a non-autonomous non-LTR sequence (*ORF1mneoI*), which resembles the structure of a rat “Half of an L1,” (*HAL1*) SINE (31), to new genomic locations (26). The efficient retrotransposition of *ORF1mneoI* mRNAs in cultured cells required the co-transfection of a “driver” plasmid that produces L1 ORF2p and an *mneoI*-based “reporter” plasmid that contains the L1 5′ UTR and produces a functional version of L1 ORF1p (26,48).

Given the above data, we next examined whether ORF1*mneoI* mRNAs could retrotranspose in HeLa strains by co-transfecting either the HeLa-HA or the HeLa-JVM strains with a L1.3 ORF2p “driver” plasmid (pTMO2F3) and an *ORF1meoI* “reporter” plasmid (Fig 6B, pJBM560). We observed readily detectable G418-resistant foci in both the HeLa-HA and HeLa-JVM strains, although we observed ∼4.6-fold fewer foci in the Hela-JVM strain (Figs 6B and S4A). Consistent with our previous analyses (48), co-transfection experiments conducted with pTMO2F3 and an RNA-binding mutant L1 ORF1 derivative of pJBM560 (pJBM560^AA^; R261A, R262A) (22,42) did not give rise to G418-resistant foci in either HeLa strain (Fig 6B). As an additional control, we demonstrated that co-transfection of pTMO2F3 with an expression plasmid containing an *EGFP* cDNA tagged with an *mneoI* retrotransposition indicator cassette (p*EGFPmneoI*) only rarely gave rise to G418-resistant foci in either HeLa strain (Fig 6B). Together, the above data suggest that *SVA*_E and *ORF1mneoI* RNAs can retrotranspose in both *Alu*-permissive and *Alu*-nonpermissive HeLa strains.

### *Alu* RNAs are reverse transcribed by L1 ORF2p in *Alu-*permissive and *Alu*-nonpermissive HeLa cells

The “ribosome association model” suggests that *Alu* RNA localizes to a ribosome, which allows the encoded *Alu* RNA poly(A) tract to compete with L1 mRNA poly(A) tail for the co-translation binding of L1 ORF2p (4,12,18,20). Thus, we next asked whether *Alu* RNA binds to ORF2p in the different HeLa strains. To address this question, we used a modified version of the LINE-1 element amplification protocol (LEAP) (56,57) to determine whether L1 ORF2p can reverse transcribe *Alu* RNAs in different HeLa strains. Briefly, the HeLa-HA or HeLa-JVM strains were co-transfected with pTMO2F3 and p*Aluneo*^Tet^ (see Methods). Forty-eight hours later, anti-FLAG antibodies were used to immunoprecipitate ORF2p-3XFLAG RNPs from whole cell lysates; the resultant ORF2p-3XFLAG RNPs then were subjected to LEAP assays using an oligonucleotide primer (RACE12T) that contained a unique 5′ sequence followed by 12 thymidine (T) residues (Fig 7A).

**Figure 7:**
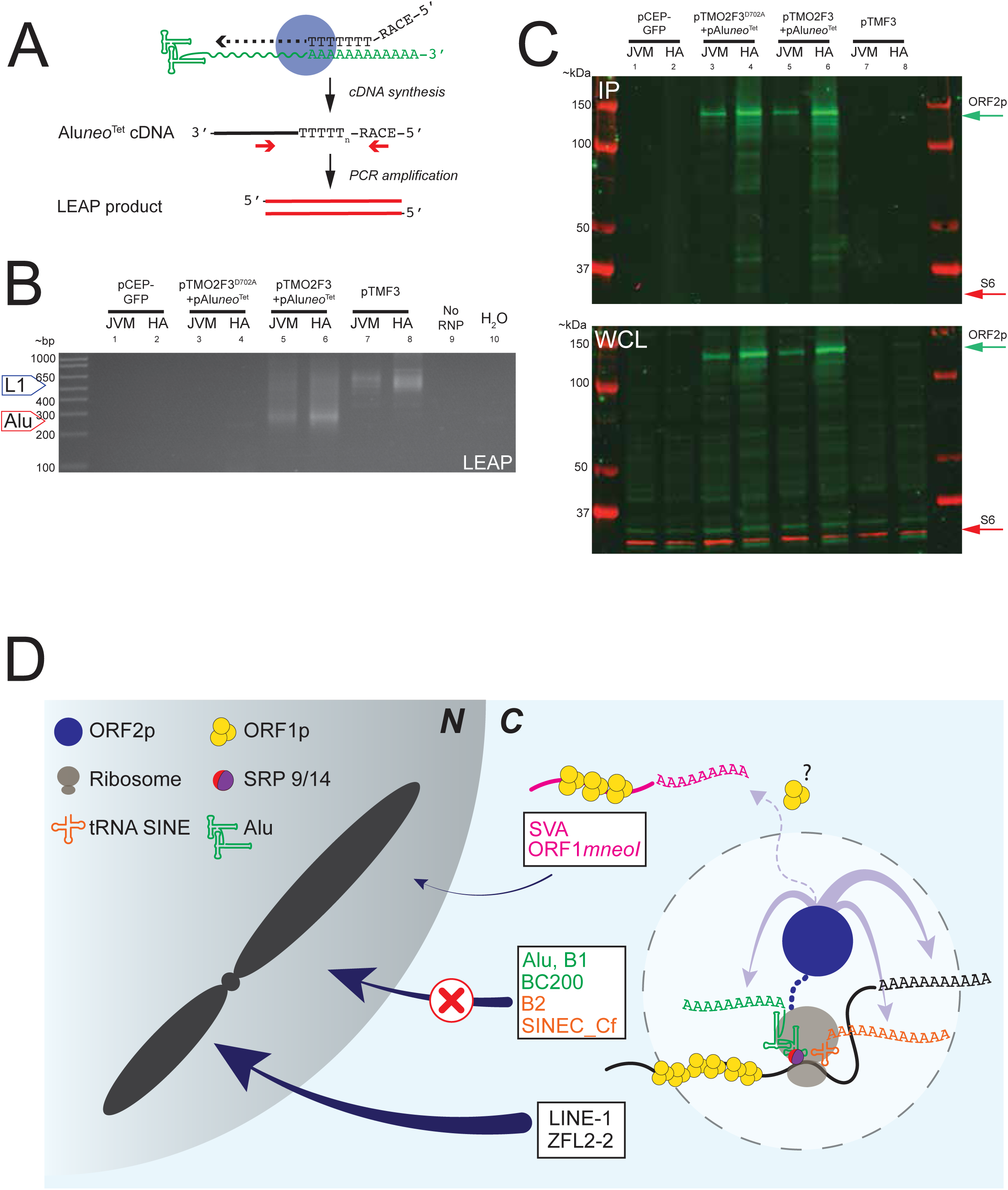
L1 ORF2p reverse transcribes *Alu* RNA in HeLa cells. *(A) LEAP assay:* HeLa cells were co-transfected with pTMO2F3 and p*Aluneo*^Tet^. Anti-FLAG antibodies then were used to immunoprecipitate ORF2p-3XFLAG (blue circle) and associated *Aluneo*^Tet^ RNAs. For the LEAP reaction, the ORF2p-3XFLAG eluates are combined with the RACE12T primer to synthesize *Aluneo*^Tet^ cDNAs. The resultant *Aluneo*^Tet^ cDNAs then were amplified using PCR with primers (red arrows) complementary to sequences at the 5′ end of the RACE12T oligo and within the 3′ end of the p*Aluneo*^Tet^ cDNA, yielding a LEAP product of ∼240 bps. *(B) LEAP results.* LEAP reactions were analyzed by 2% agarose gel electrophoresis. Transfection conditions and HeLa cell lines are indicated at the top of the gel image (No RNP = LEAP reaction control; H_2_O = PCR control). DNA size markers (in bp) are shown to the left of the gel. The predicted LEAP product sizes of *Alu* LEAP (∼240 bp; red arrow) and L1 LEAP (∼420 bp; blue arrow) reactions are indicated on the left of the gel image. The L1 LEAP products are larger due to the presence of the pCEP4 SV40 polyadenylation signal sequence at the end of the *mneoI* indicator in pTMF3. *(C) L1 ORF2p western blot results.* ORF2p-3XFLAG steady state levels were determined by Western blot using anti-FLAG IP eluates (top panel) and whole cell extracts (WCL) (bottom panel) derived from transfected HeLa-JVM and HeLa-HA cells. The lanes correspond to the LEAP gel image in Figure 7B. Anti-Flag antibodies were used to detect ORF2p-3XFLAG (green arrows). Anti-S6 antibodies (red arrows) were used to detect the S6 protein loading control. Protein size standards (in kDa) are indicated to the left of the blot images. *(D) L1 ORF2p promotes the retrotransposition of Alu and other cellular polyadenylated RNAs by distinct mechanisms. Alu* RNA (green structure) associates with ribosomes by interacting with SRP9/14 (red and purple circle) to gain access to ORF2p. The *tRNA*-derived SINE (*i.e.*, *B2* or *SINEC_Cf*; orange cruciform) localizes to ribosomes by an unknown mechanism to gain access to ORF2p. At the ribosome, newly synthesized ORF2p (blue oval) can associate (solid purple arrows) with the poly(A) tail of L1 RNA (black wavy line) or poly(A) tracts of SINE RNAs (*i.e., 7SL RNA*-derived and *tRNA* derived SINEs) to promote their retrotransposition. L1 ORF2p can also promote the retrotransposition of *SVA* RNA and/or other cellular poly(A)+ RNAs (*e.g.*, *ORF1mneoI* and mRNAs) that may not directly associate with the ribosome (dashed purple arrow) by an ORF1p-dependent mechanism that requires elucidation. *7SL-* and *tRNA*-derived SINE retrotransposition appears to be inhibited in nonpermissive HeLa cell lines after ORF2p associates with *Alu* RNA (indicated by the red “X”).

Positive control LEAP reactions, using RNPs derived from the HeLa-HA or HeLa-JVM strains transfected with an engineered human L1.3 element containing an *mneoI* reporter (pTMF3), gave rise to LEAP products of the expected size (Fig 7B, ∼430-500 bp). In contrast, negative control LEAP reactions, using RNPs derived from the HeLa-HA or HeLa-JVM strains transfected with either a GFP expressing plasmid (pCEP/GFP) or co-transfected with a reverse transcriptase missense mutant version of pTMO2F3 (pTMO2F3^D702A^) and p*Aluneo*^Tet^ and did not give rise to visible LEAP products (Fig 7B). By comparison, LEAP reactions conducted using RNPs derived from HeLa strains co-transfected with pTMO2F3 and p*Aluneo*^Tet^ yielded *Alu* reverse transcribed products of the expected size (∼240-300 bps) in both the HeLa-HA and HeLa-JVM strains (Fig 7B; note that the L1 LEAP products are predicted to be larger than the *Alu* LEAP products due to the presence the SV40 late polyadenylation signal sequence [∼192 bp] located downstream of the *mneoI* reporter in the pTMF3 plasmid (56)).

Sanger sequencing of twenty independent LEAP product clones from each HeLa strain revealed the predicted 3′ end of the *Aluneo*^Tet^ RNA followed by variable length poly(A) tracts ranging from either 10-70 or 12-74 adenosine residues in the HeLa-HA and HeLa-JVM strains, respectively. Additional M-MLV RT-PCR control assays verified the presence of *Aluneo*^Tet^ RNA in the anti-FLAG eluates from both the HeLa-HA and HeLa-JVM strain cells co-transfected with p*Aluneo*^Tet^ and either pTMO2F3 or pTMO2F3^D702A^, confirming *Aluneo*^Tet^ RNA is present at comparable levels in the ORF2p-3XFLAG RNP complexes (Fig S5A). Control western blot experiments demonstrated that ORF2p-3XFLAG is present in the whole cell lysates and eluted RNP fractions (Figs 7C and S5C) and that similar levels of SRP9/14 were present in both the HeLa-HA and HeLa-JVM strains (Fig S5B). Thus, the data indicate that *Alu* RNA associates with and is reverse transcribed by L1 ORF2p in both *Alu-*permissive and *Alu-*nonpermissive HeLa cell lines.

## DISCUSSION

We have identified two permissive HeLa cell strains (HeLa-HA, HeLa-CCL2) that support *Alu* retrotransposition and two nonpermissive HeLa cell strains (HeLa-JVM, HeLa-H1) that are severely compromised for *Alu* retrotransposition. Retrotransposition assays further revealed that additional *7SL RNA*-derived SINEs (engineered primate *Alu*Sx3, *Alu*J, and *BC200* elements as well as an engineered mouse *B1* element) and *tRNA*-derived SINEs (engineered mouse *B2* and canine *SINEC_Cf* elements) retrotranspose at high efficiencies in *Alu*-permissive, but not *Alu*-nonpermissive, HeLa strains. By comparison, both *Alu*-permissive and *Alu*-nonpermissive HeLa cell strains support LINE, *SVA* element, and *ORF1mneoI* retrotransposition. These data suggest that *7SL RNA*-derived SINEs and *tRNA*-derived SINEs retrotranspose by a similar mechanism that likely depends upon the ribosomal co-localization of these SINE RNAs and L1 ORF2p (see Fig 7D). In contrast, the retrotransposition of ORF1*mneoI*-derived mRNAs appear to require the formation of a functional ORF1p-containing ribonucleoprotein particle (RNP), which might facilitate interaction with L1 ORF2p. The mechanism by which *SVA* elements hijack L1 ORF1p and L1 ORF2p requires further elucidation (Fig 7D, see below).

The SINE retrotransposition defect in *Alu*-nonpermissive HeLa strains applies to modern dimeric human-specific *Alu*Y elements, older primate-specific *Alu*Sx elements, a reconstructed ancient monomeric *Alu*J precursor consensus element, as well as a consensus *BC200* RNA. Previous studies have demonstrated that the interaction of *Alu* RNA with SRP9/14 is dependent upon precise *Alu* RNA folding (19) and that *Alu* RNA structure plays a critical role in ribosomal localization and *Alu* retrotransposition (12). Thus, these results suggest that a conserved structural motif(s) shared between currently active and ancient reconstructed retrotransposition-competent *Alu* and *BC200* RNAs may mediate *Alu* ribosomal localization to promote retrotransposition.

Mouse *7SL RNA*-derived SINEs and both mouse and canine *tRNA*-derived SINEs exhibit similar retrotransposition dynamics as *Alu* elements in the *Alu-*permissive and *Alu*-nonpermissive HeLa strains. The RNAs encoded by both *7SL*- and *tRNA*-derived SINEs are highly structured short non-coding RNA pol III transcripts that encode 3′ poly(A) tracts that might have evolved from ribosomal associated RNAs. Current models posit that *Alu* RNA localizes to the ribosome by forming an RNP with SRP 9/14, which may allow it to effectively compete with L1 RNA for the co-translational binding of ORF2p (4,12,18,20), suggesting ribosomal localization may serve as a key regulatory step in the *Alu* retrotransposition mechanism. Based upon this model, we speculate that *tRNA*-derived SINE RNAs also localize to ribosomes to gain access to L1 ORF2p. However, how *tRNA*-derived SINE RNAs localize to ribosomes and whether this localization requires additional host factor(s) requires further investigation.

In contrast to *7SL RNA*-derived and *tRNA*-derived SINEs, an *SVA_E* element as well as human and mouse ORF1*mneoI* RNAs retrotranspose in both *Alu-*permissive and *Alu*-nonpermissive HeLa cell lines (Fig 6). Although *Alu* and *tRNA*-derived SINE retrotransposition only require L1 ORF2p (4), our data agrees with previous studies suggesting that *SVA* and ORF1*mneoI* retrotransposition requires both L1 ORF1p and L1 ORF2p (26,29,30). Intriguingly, previous studies have shown that *SVA* RNA can be detected in L1 RNPs with other SINE RNAs and mRNAs known to form processed pseudogenes (77). However, whether the association of these RNAs with L1 RNPs is an active process that facilitates their retrotransposition, whether *SVA* and/or other cellular RNAs (*e.g.*, mRNAs that are used as substrates for processed pseudogenes) need to transit to the ribosome to acquire L1 ORF2p, and how *SVA* and other polyadenylated cellular RNAs compete for L1 protein binding requires elucidation.

Our LEAP, western blot, and RT-PCR data (Figs 7 and S5) demonstrate that: (**1**) *Alu-*permissive and *Alu-*nonpermissive HeLa cell lines express similar levels of SRP9 and SRP14, indicating sufficient SRP9/14 is available for *Alu* RNP formation; (**2**) *Alu* RNA associates with ORF2p in both *Alu*-permissive and *Alu*-nonpermissive HeLa strains; and (**3**) *Alu* RNAs are reverse transcribed into *Alu* cDNAs in *Alu*-permissive and *Alu*-nonpermissive HeLa strains. Together, these data indicate that L1 ORF2p binds to the 3′ poly(A) tract of *Alu* RNA in both *Alu*-permissive and *Alu*-nonpermissive HeLa strains and suggests that *Alu* retrotransposition may be blocked in *Alu*-nonpermissive HeLa cell lines after the association of L1 ORF2p with *Alu* RNA (Fig 7D).

DNA sequencing and karyotype analyses indicate that *Alu*-permissive HeLa strains are genetically distinct from *Alu*-nonpermissive HeLa strains, suggesting that genetic differences likely underlie the *Alu* and tRNA-derived SINE retrotransposition differences observed among these HeLa strains (Fig 2). Previous studies have shown that different HeLa cell strains exhibit distinct genetic and phenotypic differences (54,78–81). Our data suggest that the *Alu*-permissive HeLa-HA is genetically and phenotypically related to the *Alu*-permissive HeLa-CCL2 strain, which was the first HeLa cell line deposited to the ATCC (82,83). In contrast, the *Alu*-nonpermissive HeLa-JVM strain is more closely related to the *Alu*-nonpermissive HeLa-H1 (CRL-1958) strain, which was derived from the original HeLa cell line established by George Gey and colleagues (82,83) and deposited to the ATCC years after the HeLa-CCL2 strain. Thus, it appears that the HeLa-H1, and by proxy HeLa-JVM, strains could be derived from the original HeLa-CCL2 strain.

Our findings raise the following question: why do some SINEs show markedly different retrotransposition efficiencies in different HeLa isolates? We speculate that differences in host factors between *Alu-*nonpermissive and *Alu-*permissive HeLa isolates could account for these differences. Possible models to account for these differences in SINE retrotransposition include: (1) *Alu*-nonpermissive HeLa strains lack necessary host cell factor(s) that promote *Alu* retrotransposition; or (**2**) *Alu*-nonpermissive HeLa strains express dominant host cell factor(s) that suppress *Alu* retrotransposition. It is notable that previous studies have exploited differences between permissive and nonpermissive cell lines to identify host cell factors that influence retrovirus and retrotransposon activity (84,85). Our future studies will seek to identify host factors responsible for the differences in SINE retrotransposition among the permissive and non-permissive HeLa strains.

### CONCLUSIONS

The differences of LINE, *7SL RNA*-derived SINE, *tRNA*-derived SINE, *SVA* element, and a proxy for a *HAL1*-like SINE (*ORF1mneoI*) retrotransposition dynamics in different HeLa strains support the idea that distinct mechanisms exist for the L1-mediated retrotransposition of different cellular RNAs (25,29,30,48,64,86,87) (Fig 7D). A critical requirement common to all these retrotransposition mechanisms is their dependence on L1 ORF2p. L1 *cis*-preference and RNA pol III SINE (*i.e.*, *Alu*, *SINEC_Cf*) retrotransposition are likely dependent on ribosomal localization, which allows the L1 3′ poly(A) tail or the encoded *7SL RNA*-derived/*tRNA*-derived SINE encoded poly(A) tract to compete for binding L1 ORF2p (4,12,18,20). However, it remains unclear whether *SVA*, *ORF1mneoI*, and/or processed pseudogene mRNA precursors also need to transit the ribosome to gain access to L1 ORF2p. Notably, *U6 small nuclear RNA* (snRNA), and likely *U6_ATAC_ snRNA*, are proposed to utilize a different ligation-based mechanism to mediate their retrotransposition, where an RNP complex containing *U6 snRNA*, a putative endo-ribonuclease, and the RtcB RNA ligase, act in concert to ligate *U6 snRNA* directly to L1 RNA after L1 RNP formation in the nucleus; retrotransposition of the ligated U6/L1 fusion transcript then leads to the creation of *U6/L1* chimeric pseudogenes (87–89). Thus, our analyses shed additional light on the myriad of mechanisms by which the L1 ORF1p and/or L1 ORF2p-mediated *trans*-mobilization of cellular RNAs continue to diversify the human and likely other mammalian genomes.

## ACKNOWLEDGEMENTS

We thank Dr. Annette Damert for providing the *SVA* element construct (pAD26), Dr. Thierry Heidmann for providing the p*Aluneo*^Tet^ and *neo*^Tet^ constructs, Dr. Astrid Engel for providing the HeLa-HA strain, Dr. Thomas Widmann and members of the Moran and Garcia-Perez laboratories for helpful advice and helpful critiques throughout the study, and the University of Michigan Advanced Genomics Core for assistance in generating next generation sequencing data. We also thank Isabel Maleno and Federico Garrido (Hospital Universitario Virgen de las Nieves, Granada, Spain) for conducting blinded HLA typing and STR analyses. We are grateful to Henrietta Lacks, now deceased, and to her surviving family members for their contributions to biomedical research. The HeLa cell line that was established from her tumor cells without her knowledge or consent in 1951 has made significant contributions to scientific progress and advances in human health.

## AUTHOR CONTRIBUTIONS

JBM, HCK, JLGP, and JVM designed the study. JBM, HCK, YL, MG-C, PC, PEL, LS, JOK, and JMK conducted wet bench experiments and/or computational analyses. JBM, HCK, YL, MG-C, PC, PEL, LS, JOK, JMK, JLGP, and JVM analyzed the data. JBM wrote the initial version of the manuscript. JBM, JLGP, and JVM edited the manuscript. All authors had a chance to comment on the manuscript.

## DATA DEPOSITION

Low passage Illumina whole genome sequencing (WGS) data are deposited in the HeLa WGS resource under study phs003397.

## SUPPLEMENTARY FIGURE LEGENDS

**Supplementary Figure S1, Related to Figure 1:**
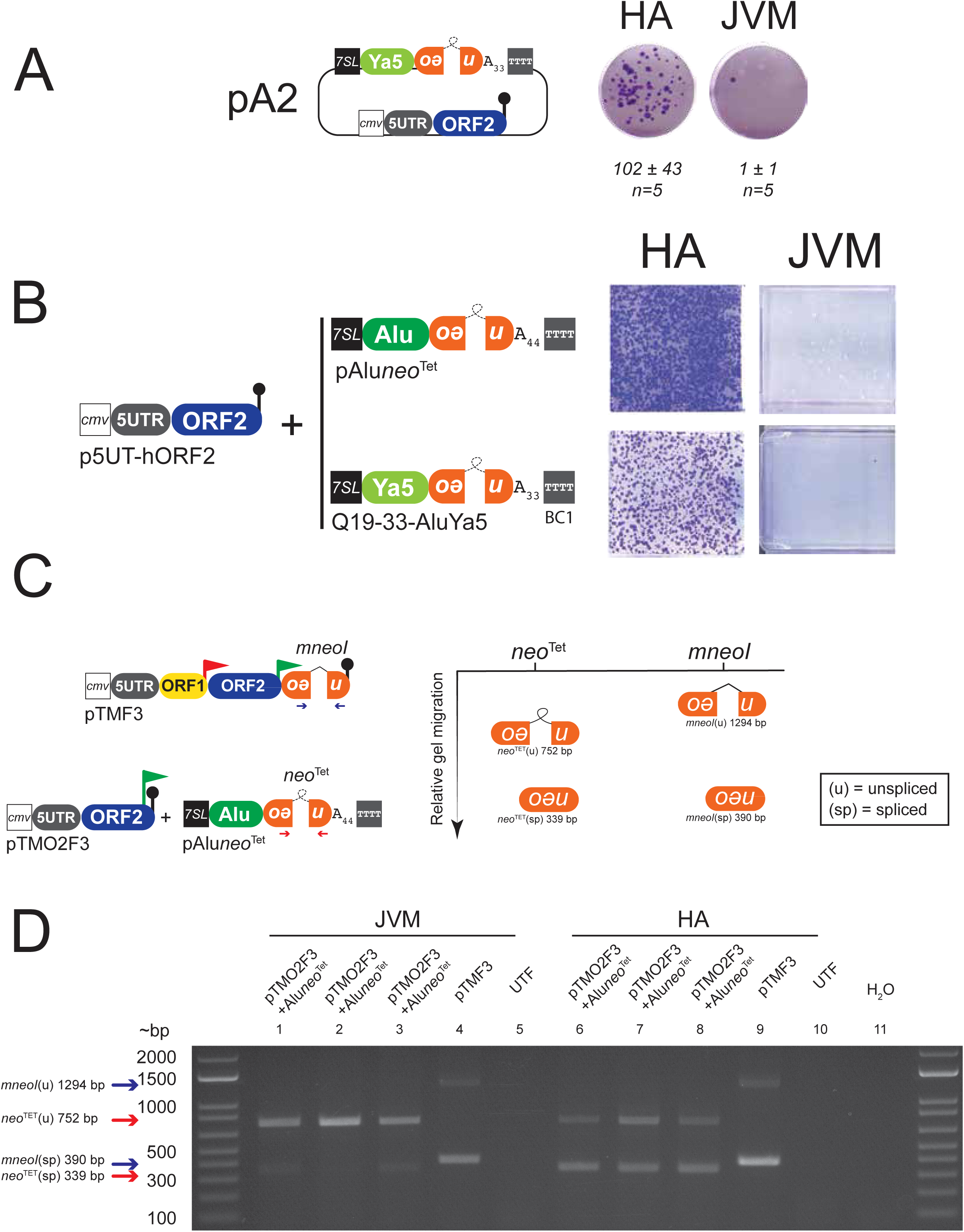
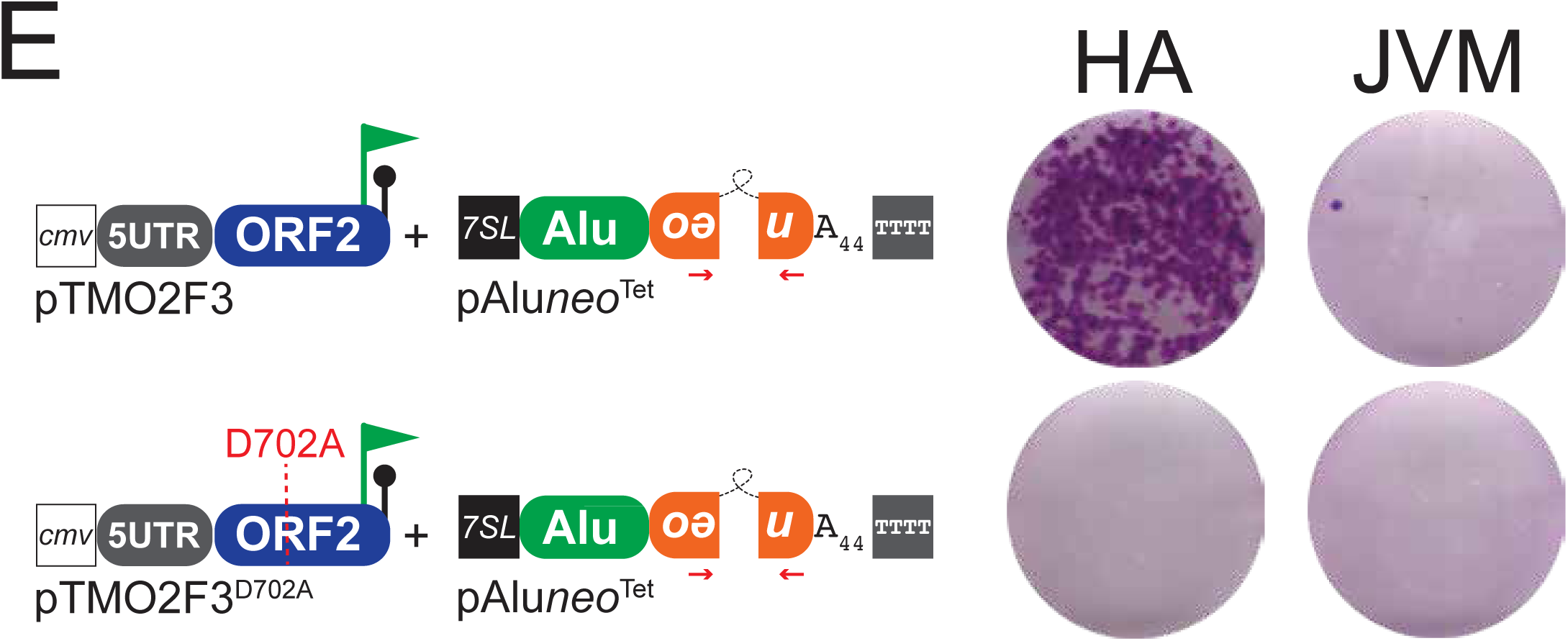
*(A) Retrotransposition results for pA2. Left panel:* Diagram of the pA2 plasmid. *Right panel:* pA2 retrotransposition results. Displayed are single wells of a representative six-well tissue culture plate from pA2 retrotransposition assays (where both the L1 ORF2 “driver” and *Alu* “reporter” genes are expressed from the same plasmid). The HeLa strain is denoted above each well. Below each well is the average number of G418-resistant colonies per well ± standard deviation (n=number independent transfections). *(B) Retrotransposition results for AluYa5. Left panel:* Diagram of plasmids used in the assay. Q19-33-*Alu*Ya5 expresses a human *Alu*Ya5 element marked with a *neo*^Tet^ marker (see main text for details) and p5UT-hORF2 expresses an untagged version of human ORF2p. A CMV promoter (white square), the native L1 5′ UTR (grey oval), and an SV40 polyadenylation signal sequence (black lollipop) augment p5UT-hORF2 expression. *Right panel:* Qualitative *Alu*Ya5 retrotransposition results. Displayed are images of a representative T-75 flask. *(C) Left panel: Diagram of constructs used in the PCR-based retrotransposition assay.* TMF3 is identical to pJM101/L1.3 (see Fig 1 legend) except that TMF3 contains a *T7 gene 10* epitope tag (red flag) at the 3’ end of *L1 ORF1* and a 3XFLAG tag at the 3’ end of *L1 ORF2* (green flag). The relative positions of forward and reverse PCR primers (arrows) are indicated underneath the *neo* retrotransposition indicator cassettes. *Right panel:* Relative sizes of expected PCR products from unspliced (u) or spliced (sp) retrotransposition indicator cassettes. *(D) Results from PCR assays for intron removal (retrotransposition) in HeLa-JVM (lanes 1-5) and HeLa-HA (lanes 6-10).* Transfection conditions are indicated above each lane. Each lane represents an independent transfection. The top band in each lane corresponds to unspliced (u) PCR products derived from the transfected plasmid DNA; the lower band corresponds to spliced (sp) PCR products derived from the integrated (*i.e.*, retrotransposed) retrotransposition indicator cassette. UTF=untransfected cells; H_2_O=PCR negative control. The experiment was repeated twice with similar results. *(E) Retrotransposition results for pAluneo^Tet^ co-transfected with an L1 ORF2p reverse transcriptase mutant. Left panel:* Diagram of plasmids used in the assay. p*Aluneo*^Tet^ was co-transfected with an ORF2p “driver” plasmid that expresses wild type ORF2p (pTMO2F3) or a reverse transcriptase mutant version of ORF2p (pTMO2F3^D702A^) *Right panel:* Qualitative retrotransposition results. Displayed are images of a representative well of a 6-well plate. The HeLa strain is denoted above each image and the transfected construct is displayed to the left of the images.

**Supplementary Figure S2, Related to Figure 2:**
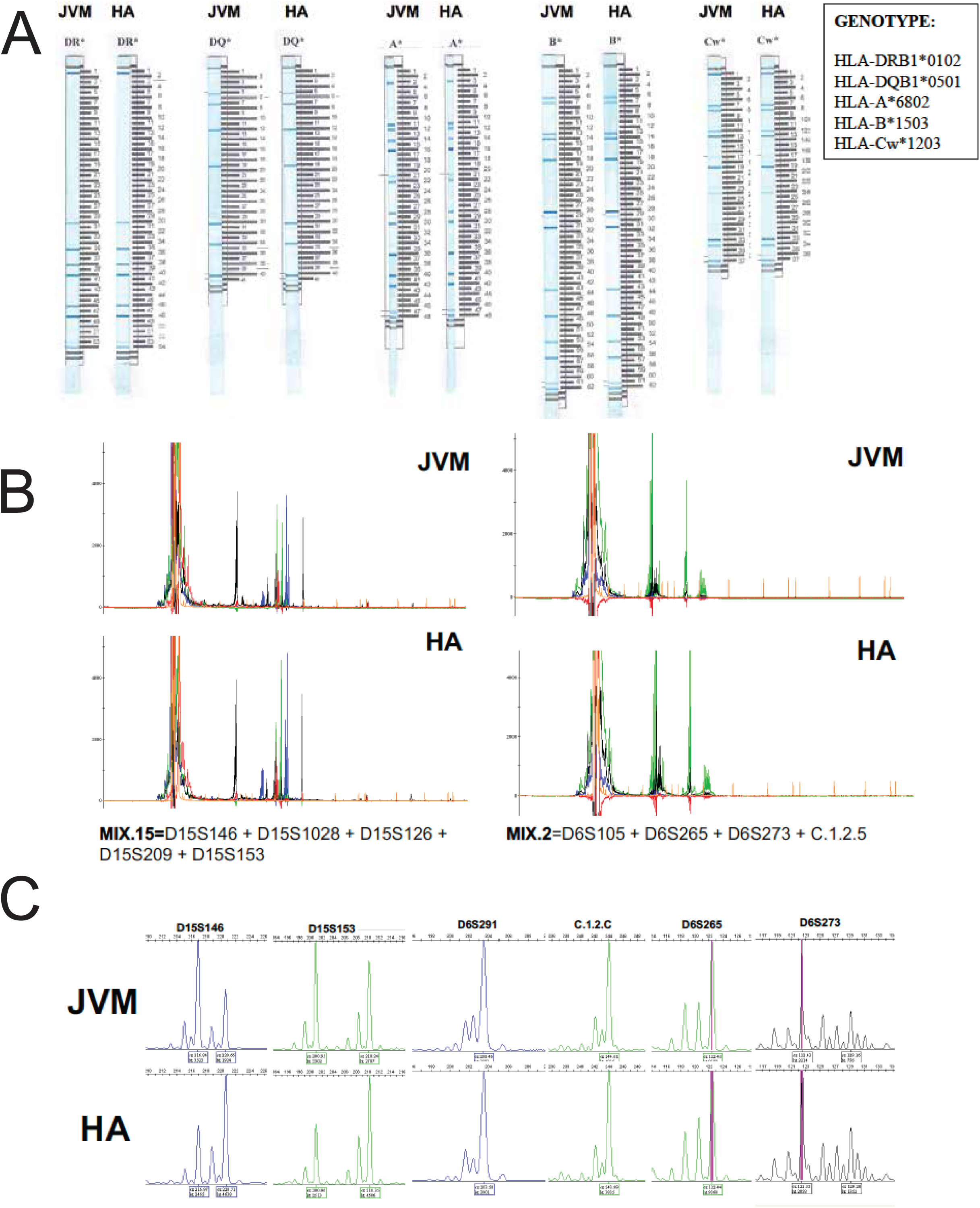
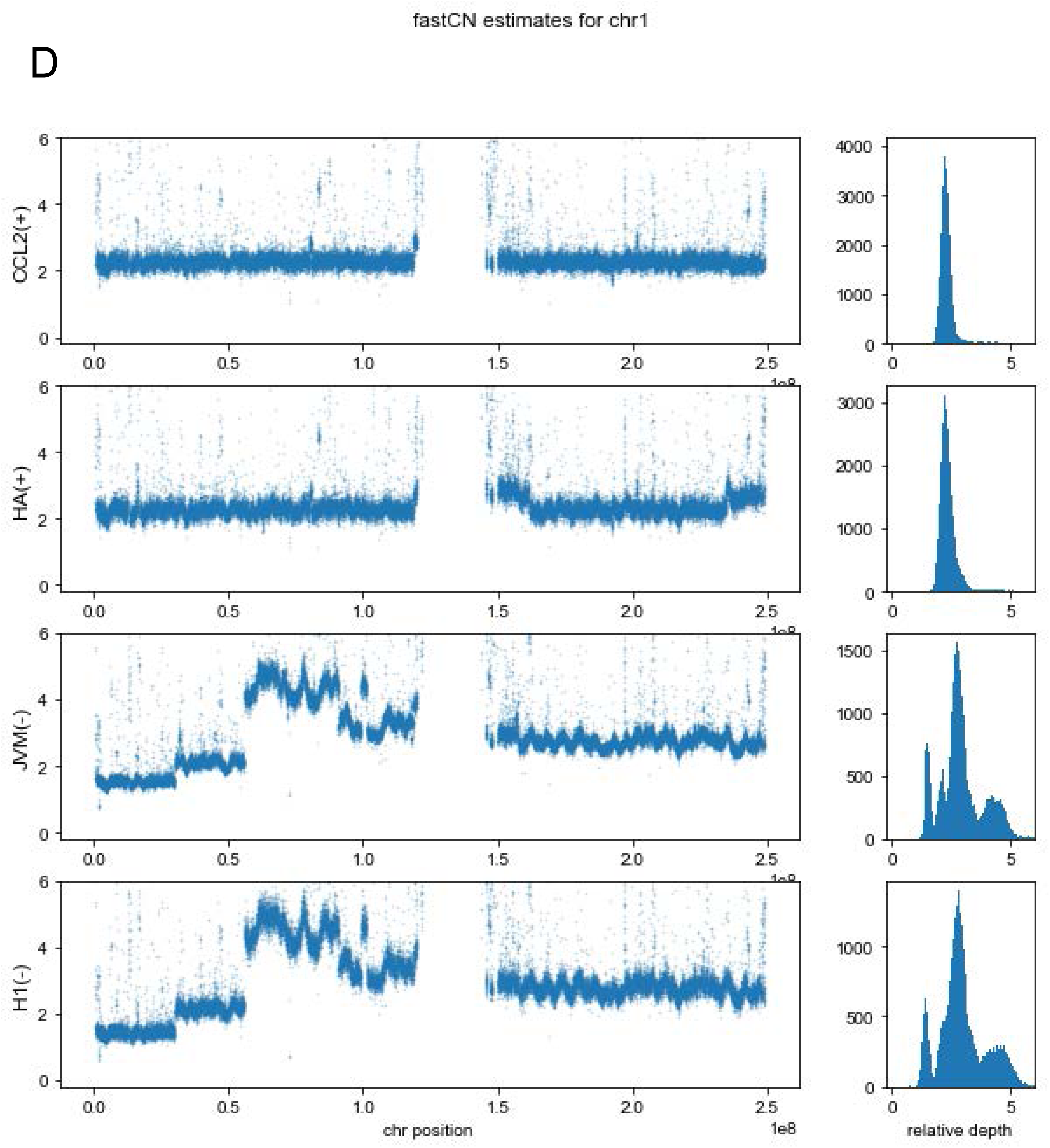
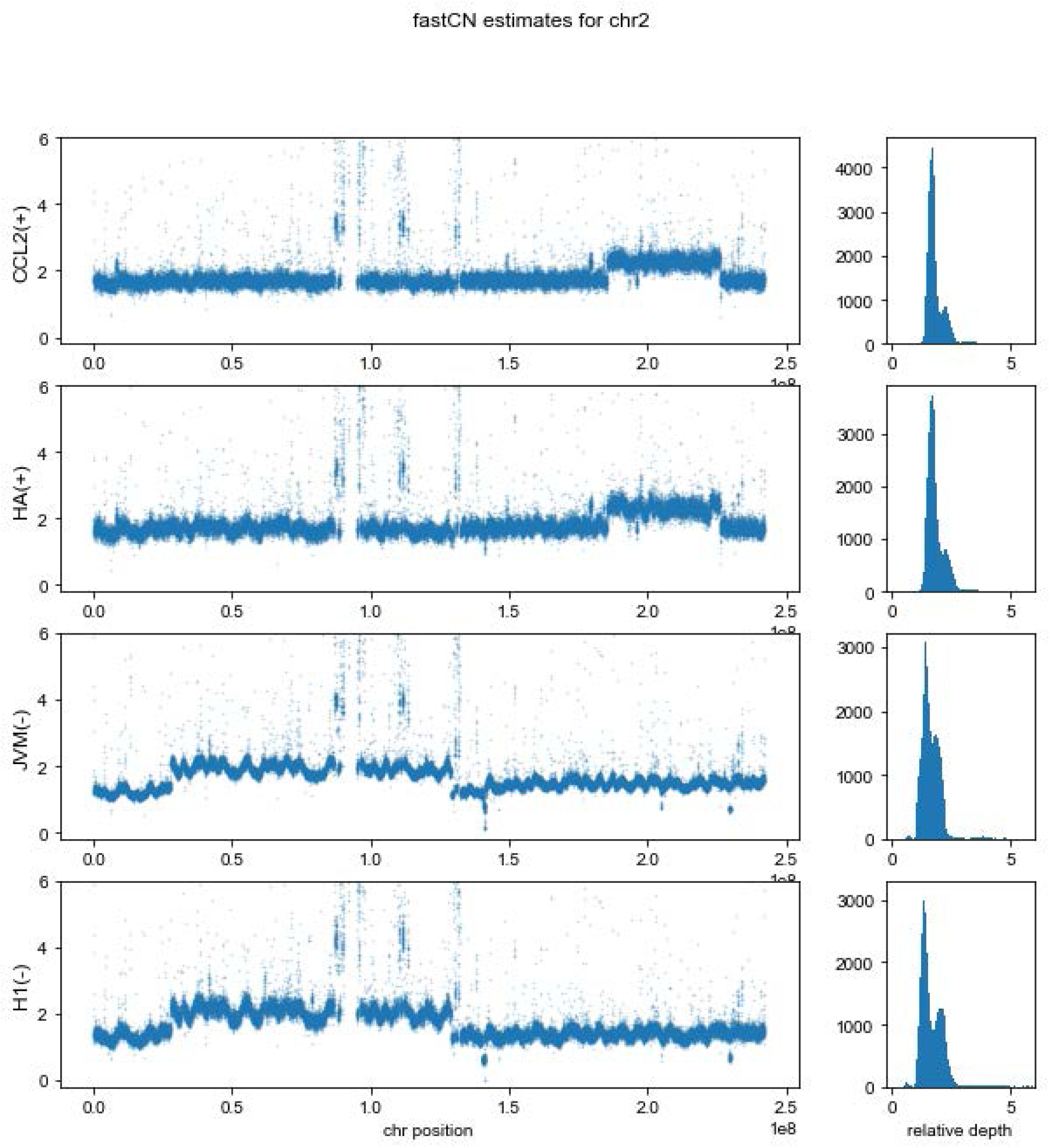
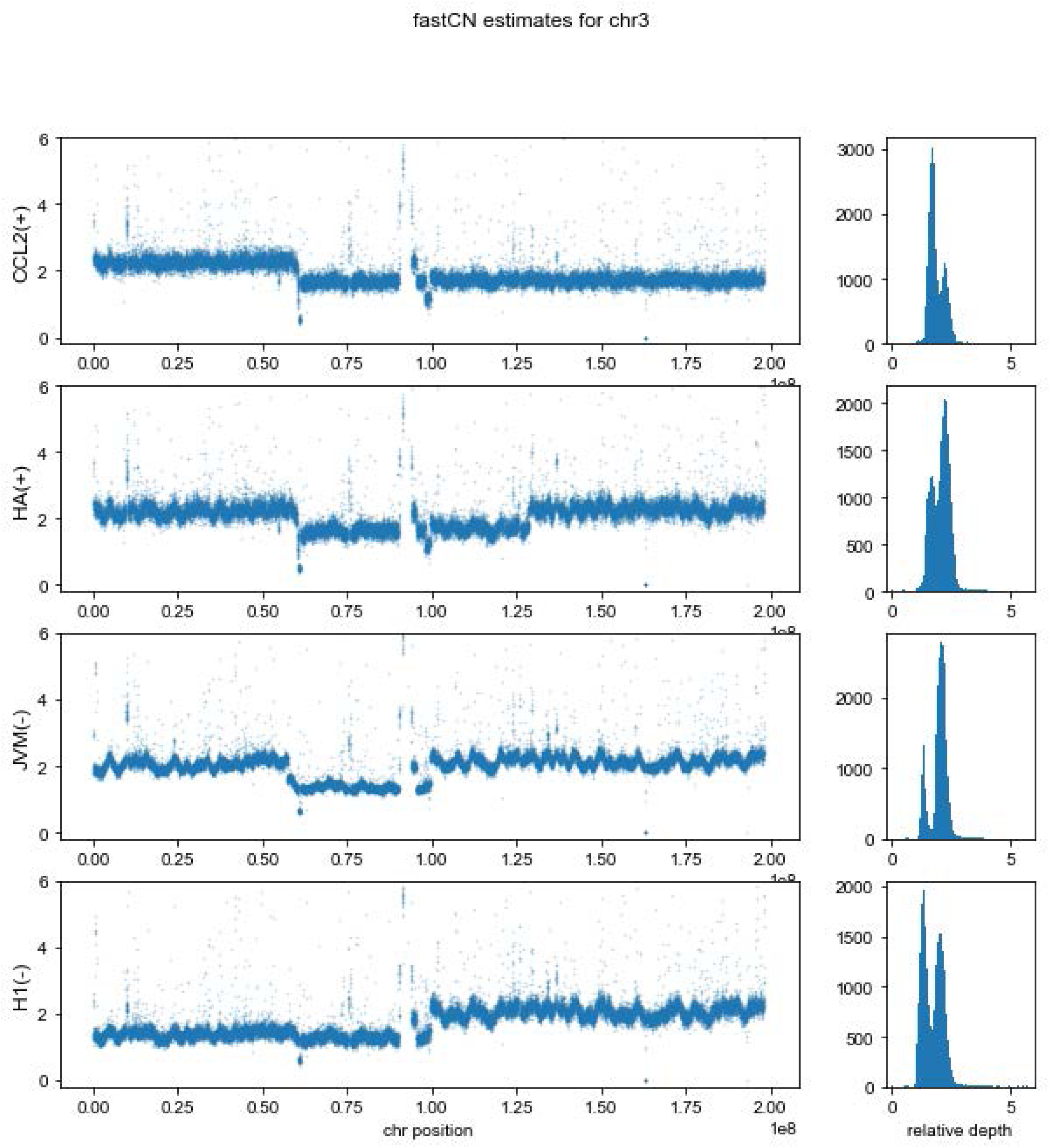
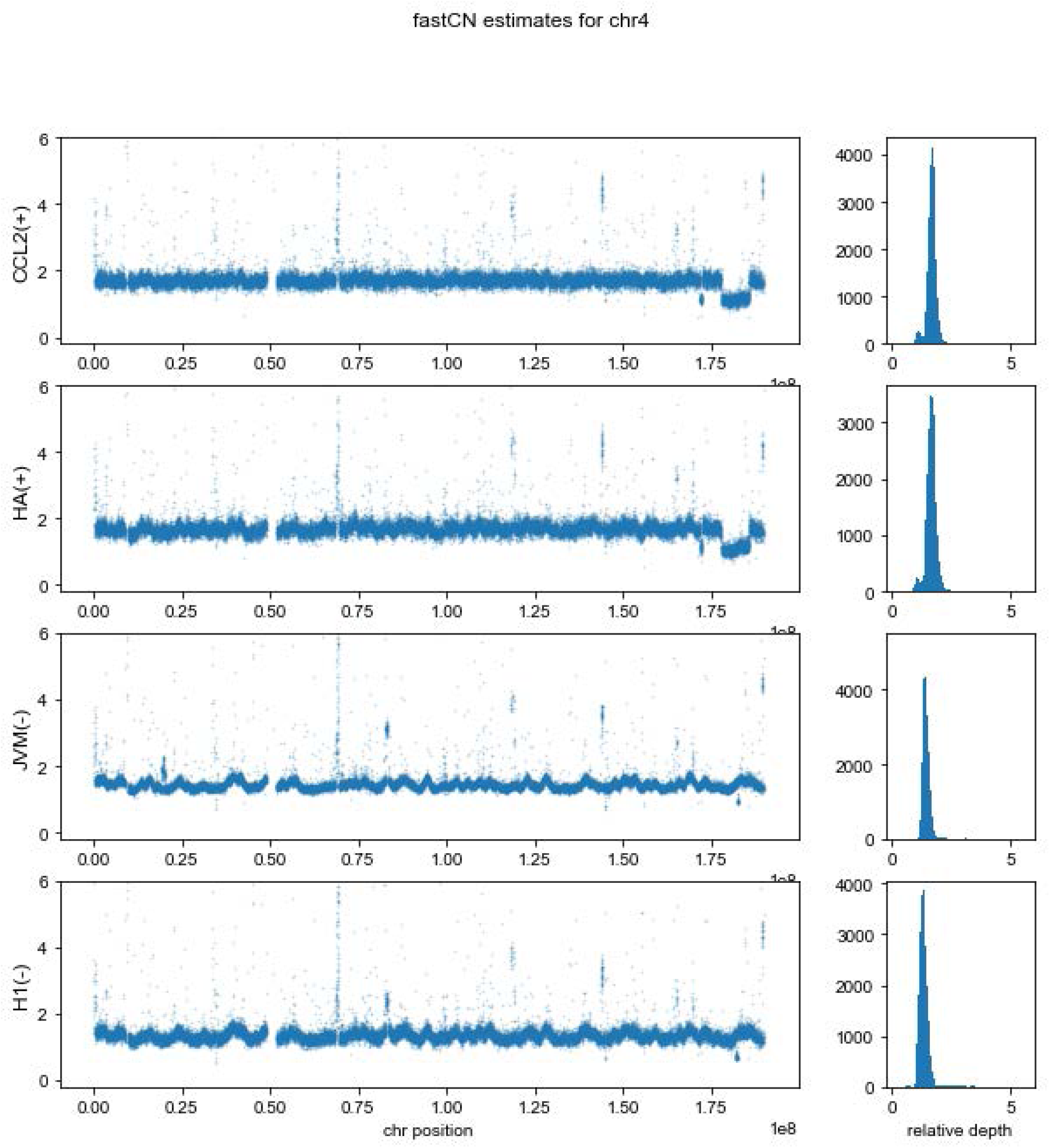
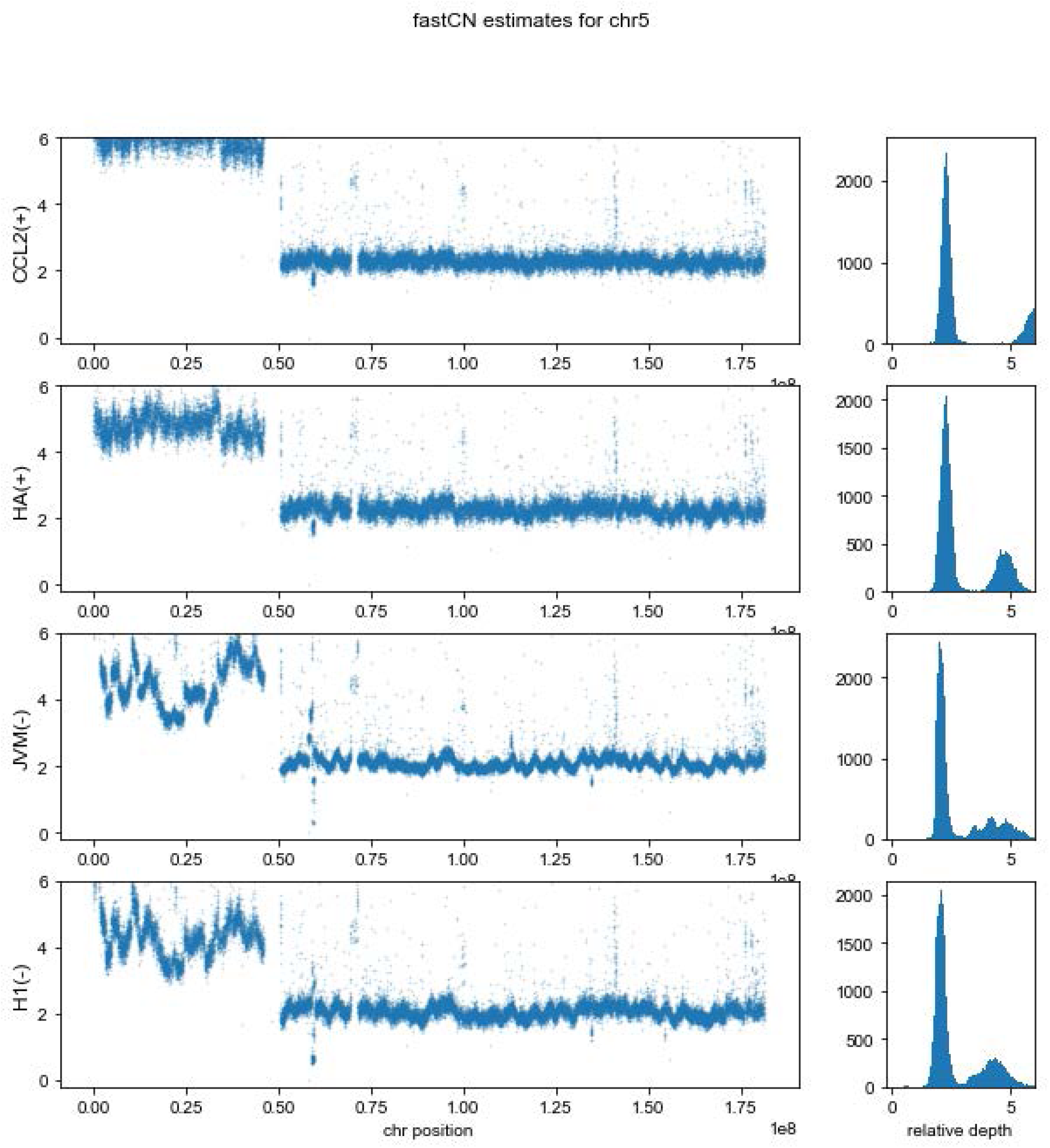
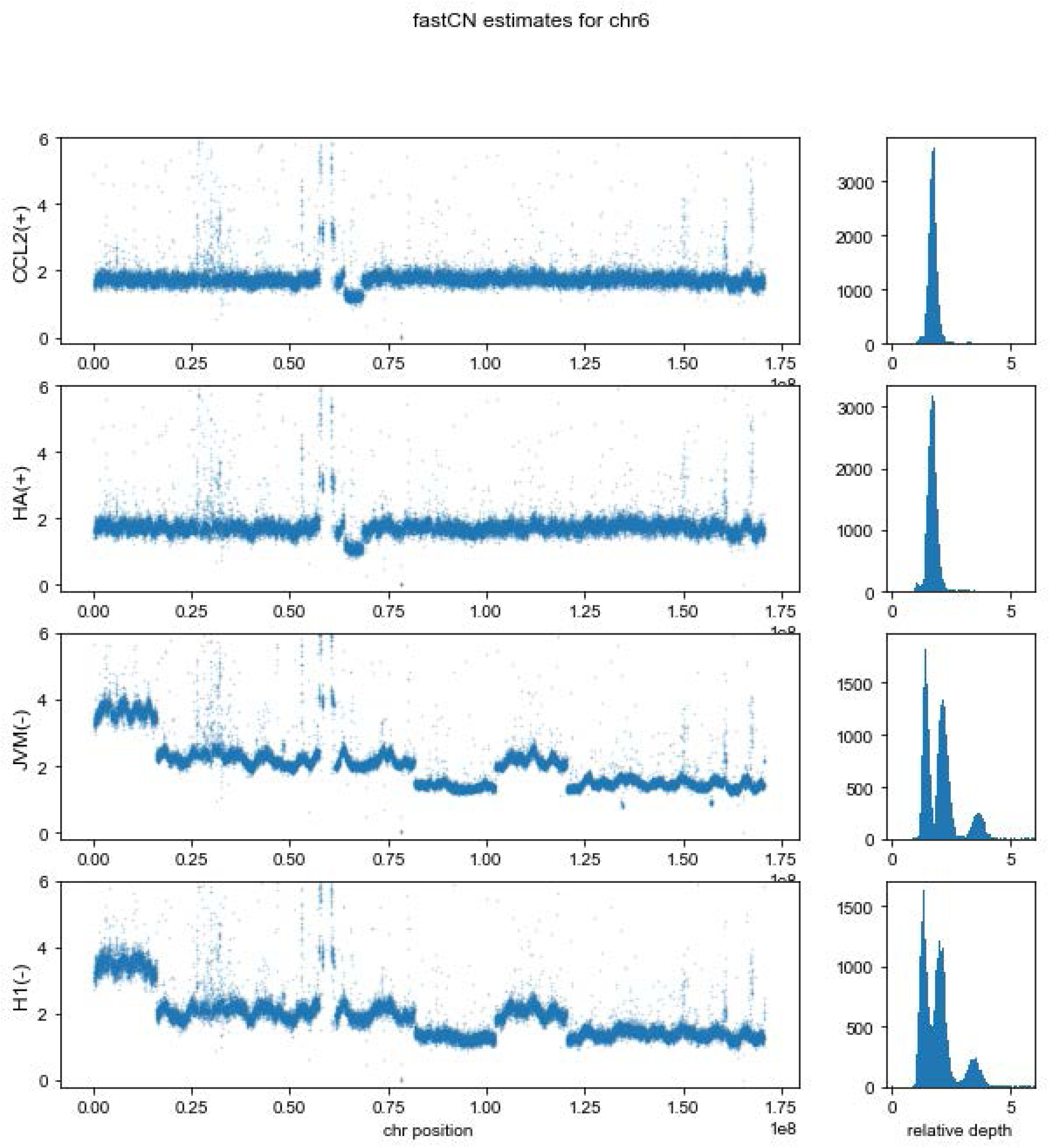
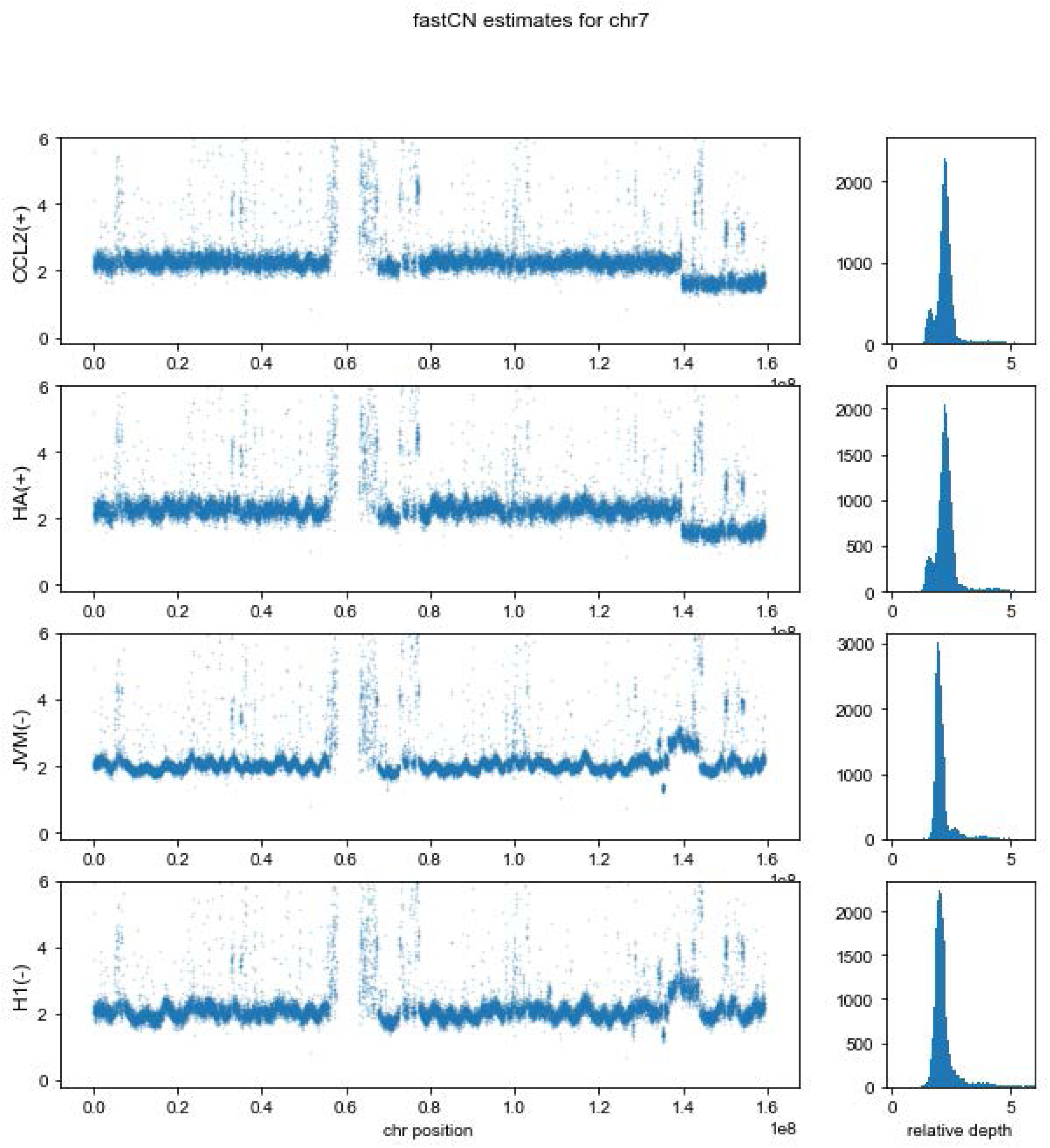
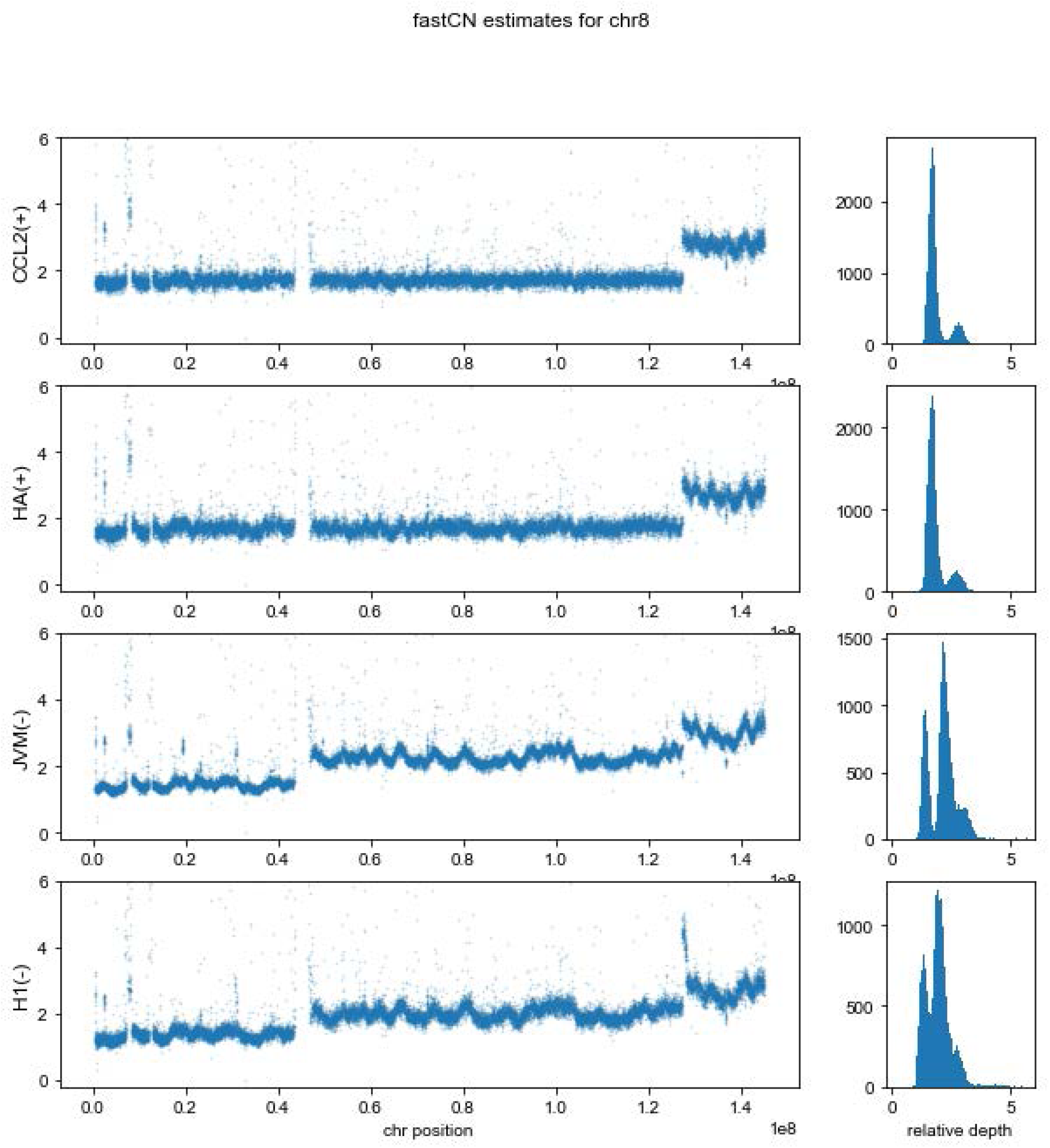
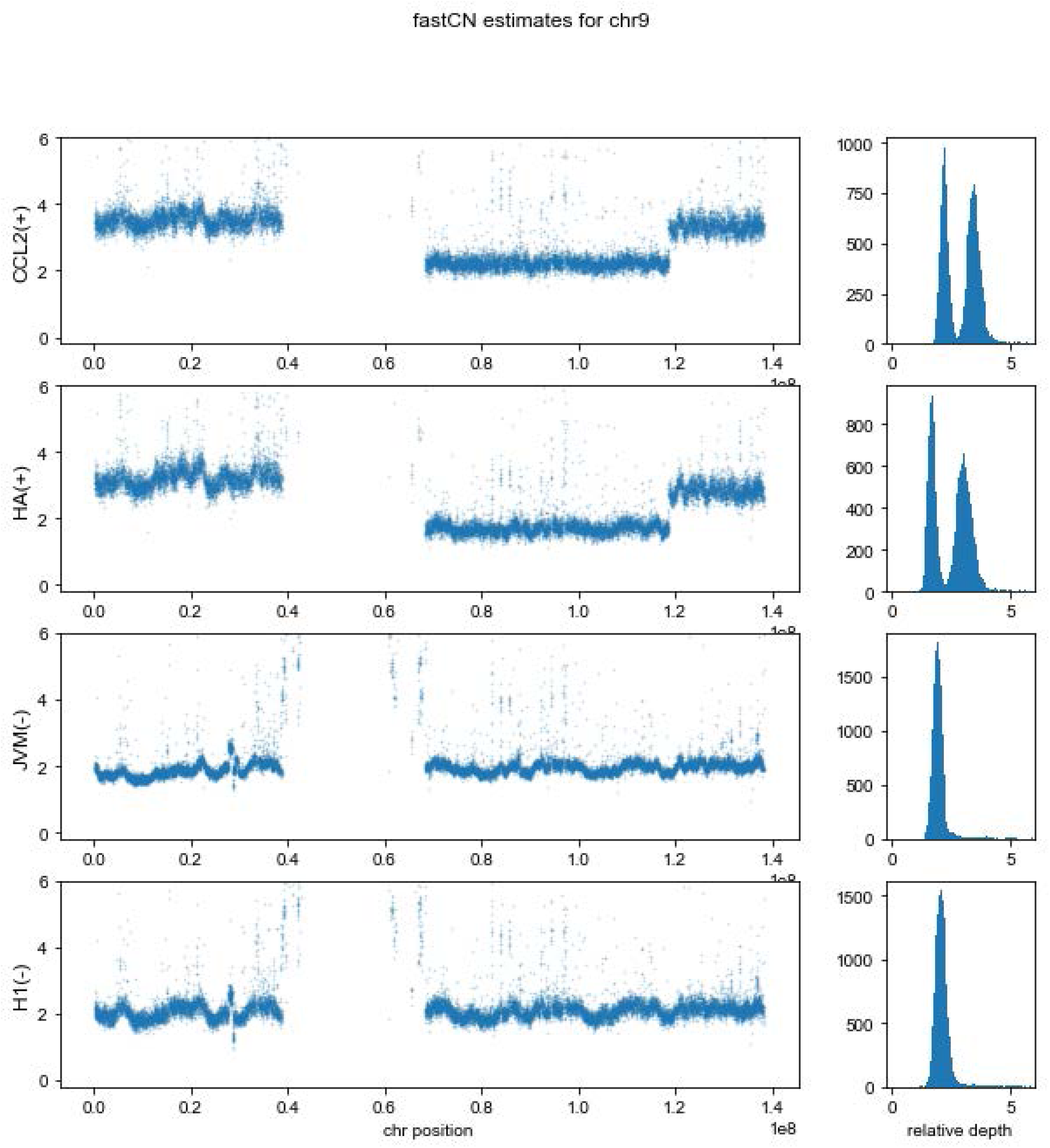
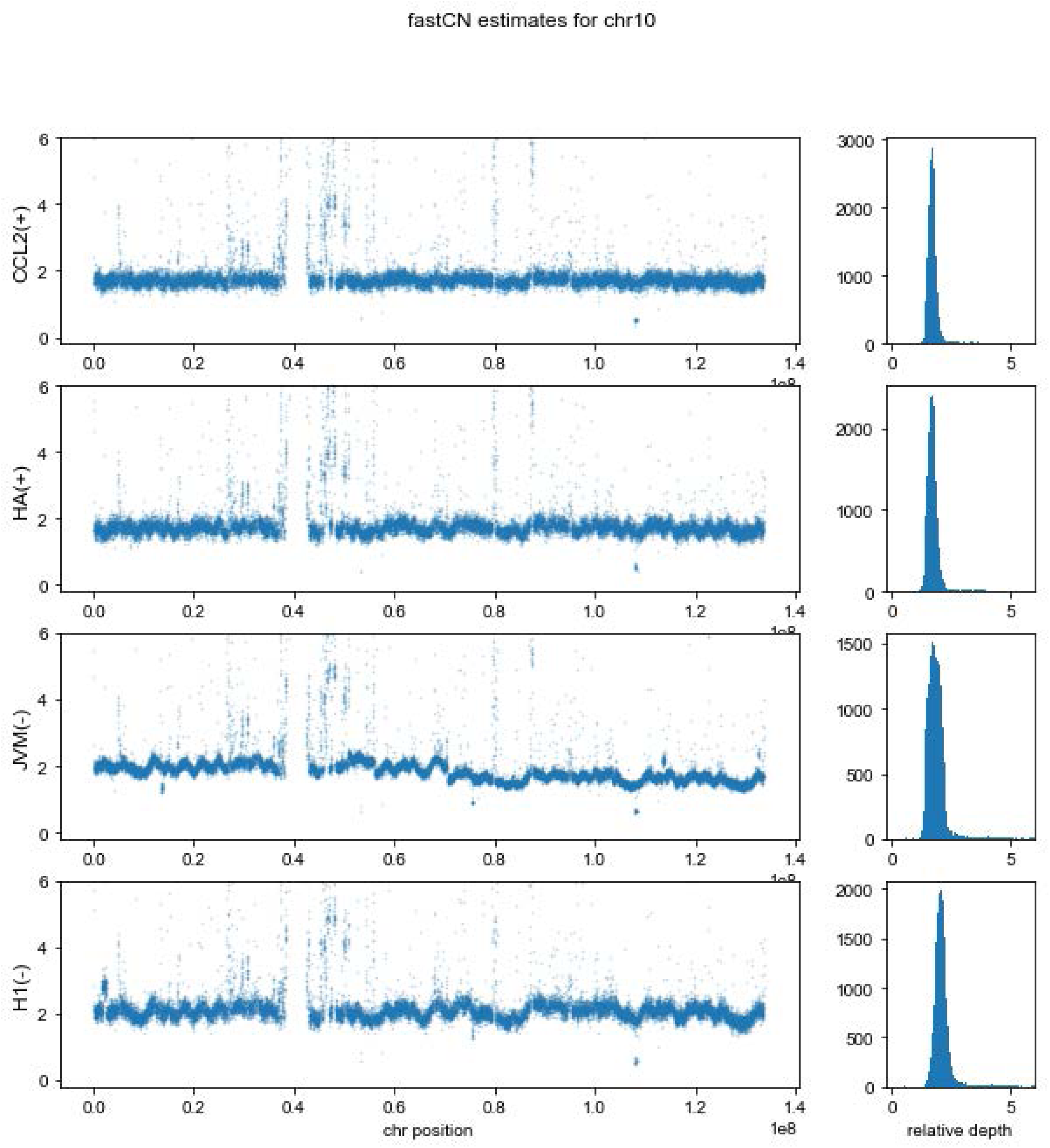
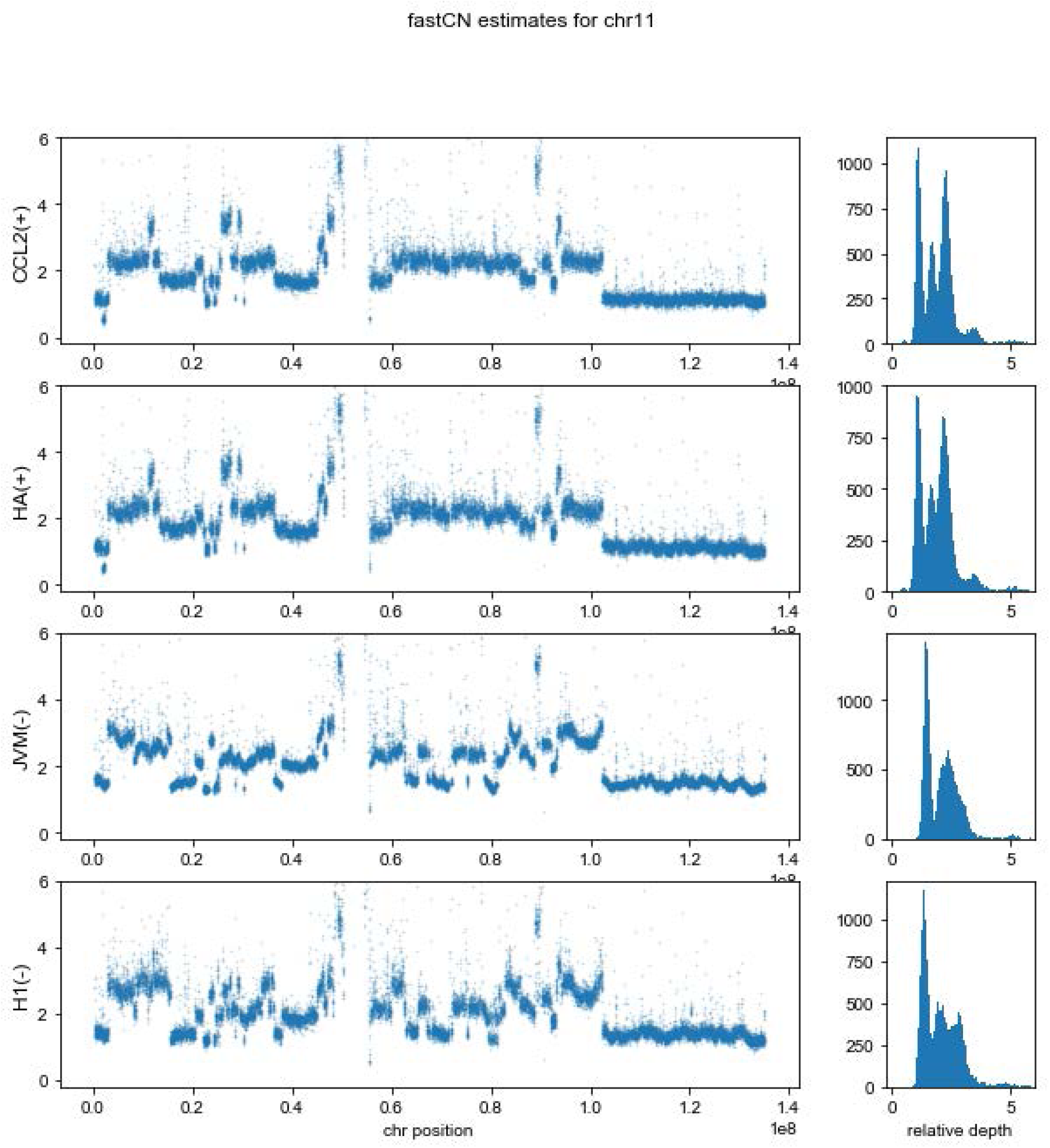
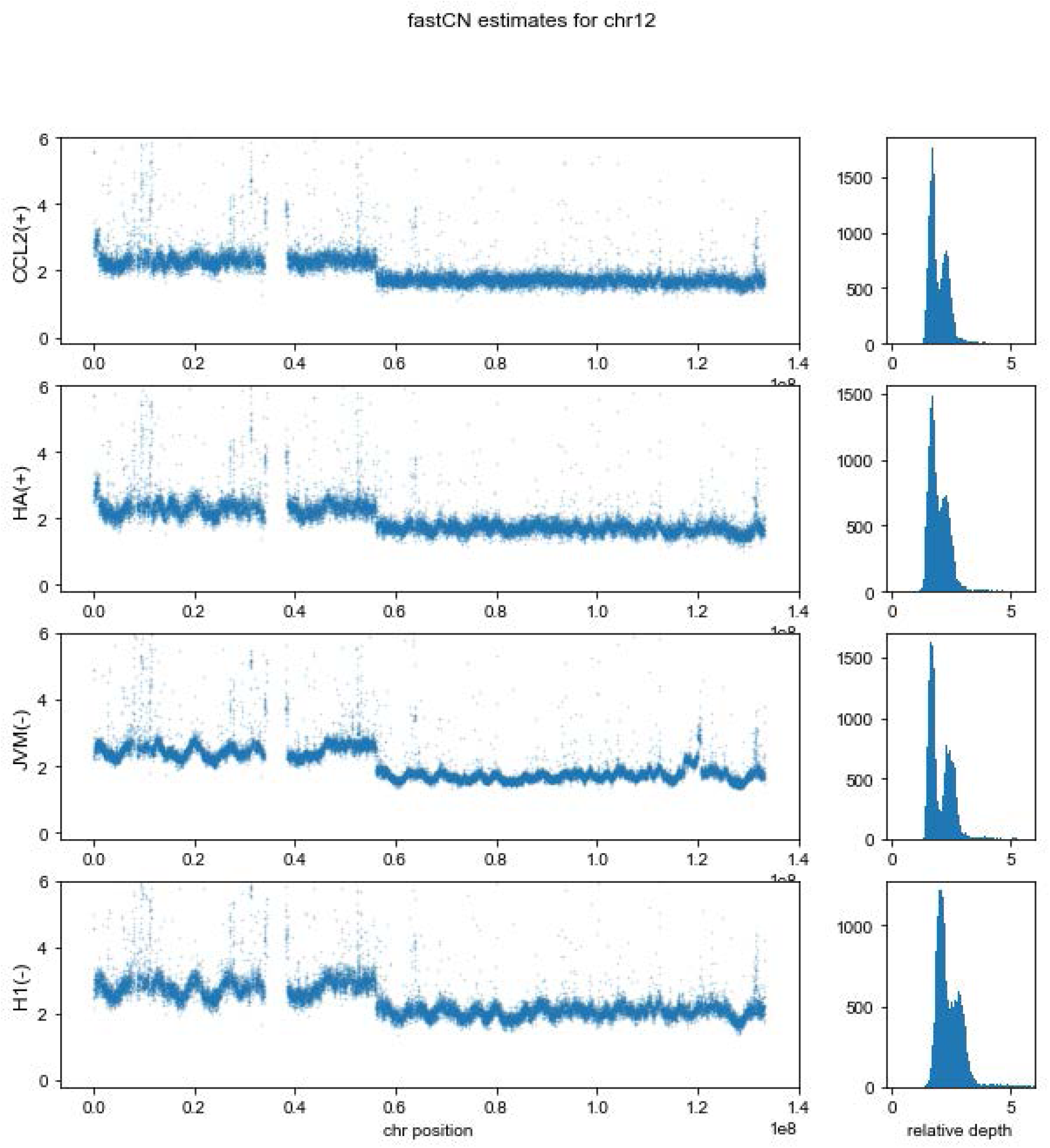
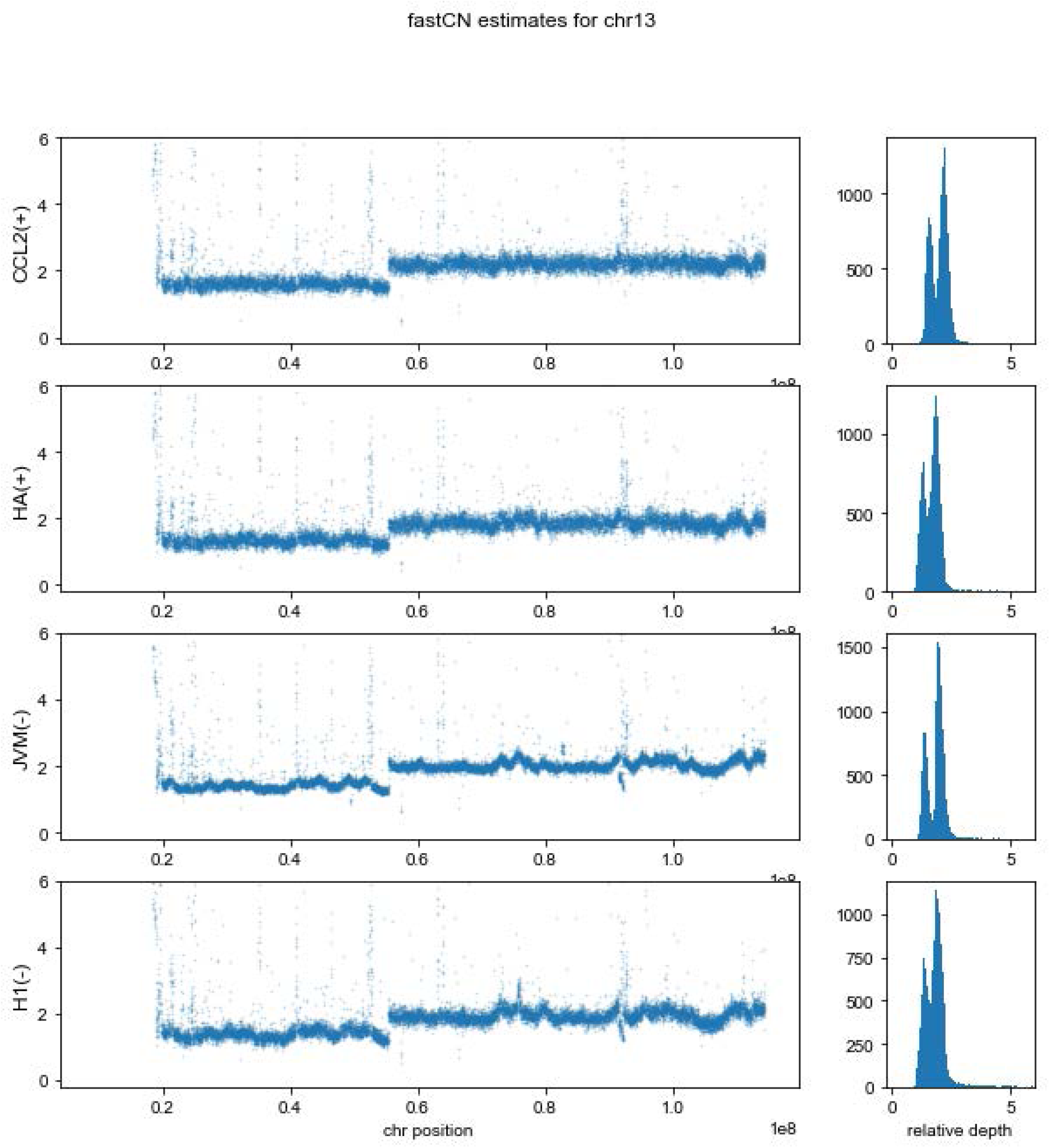
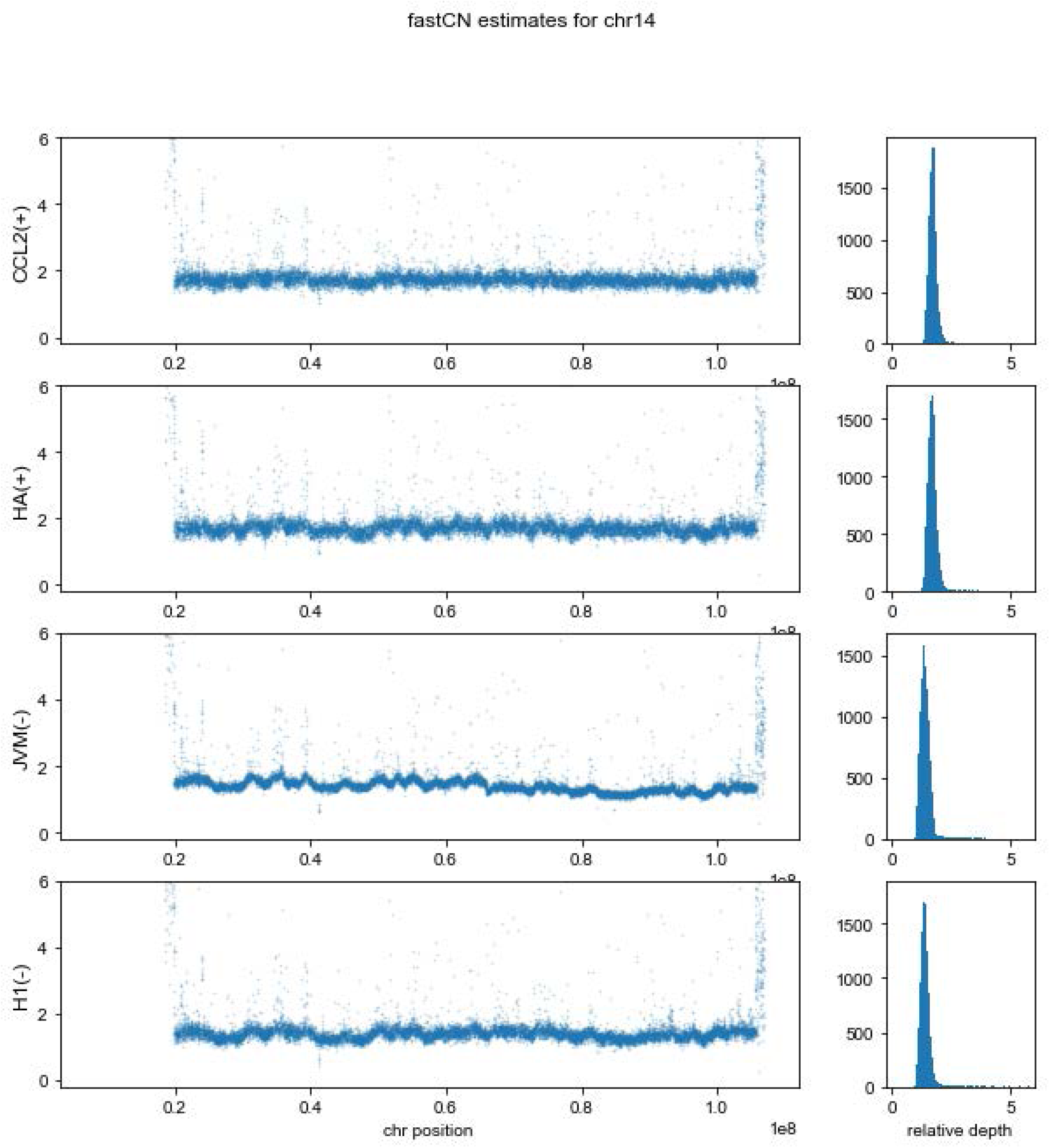
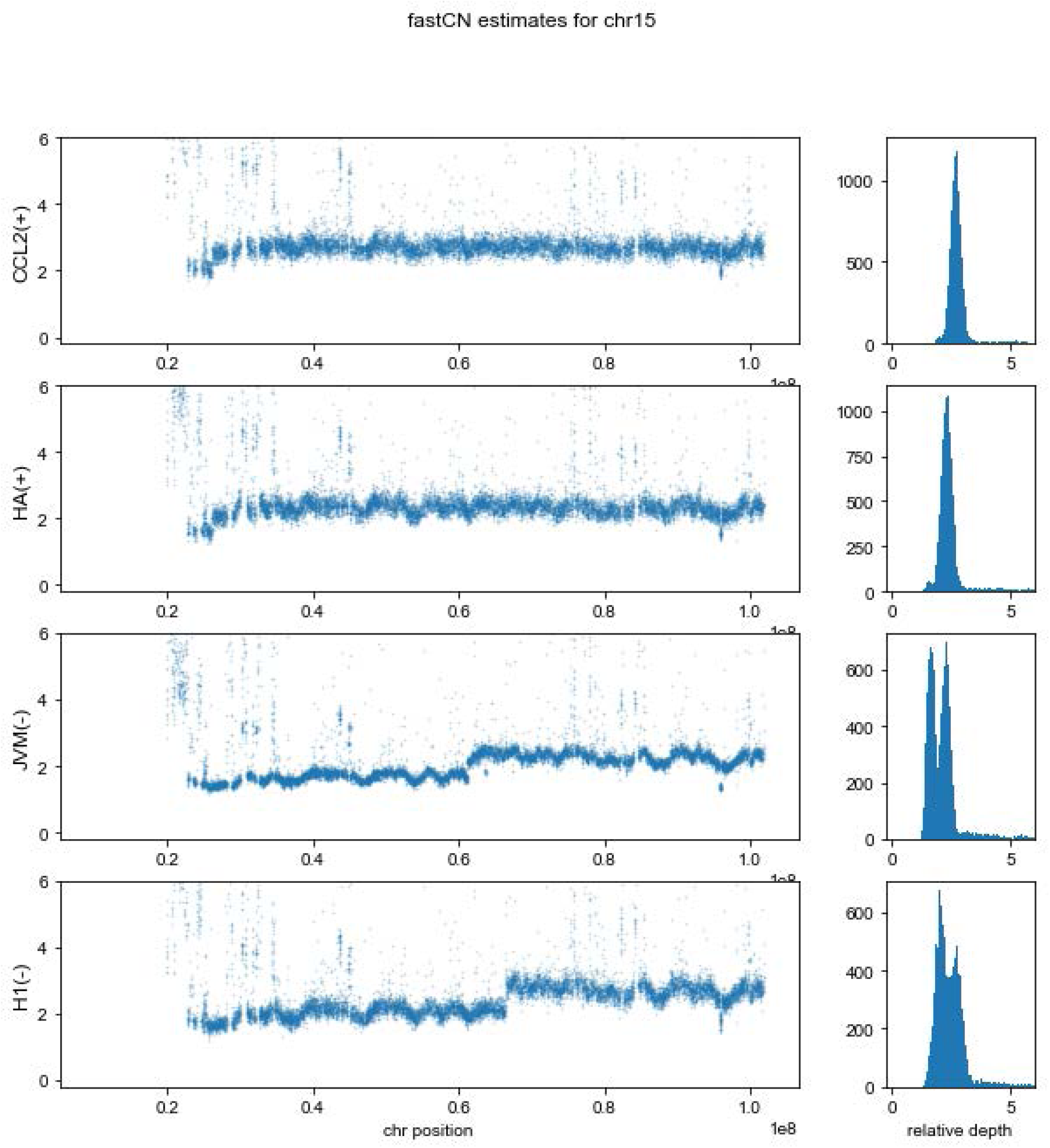
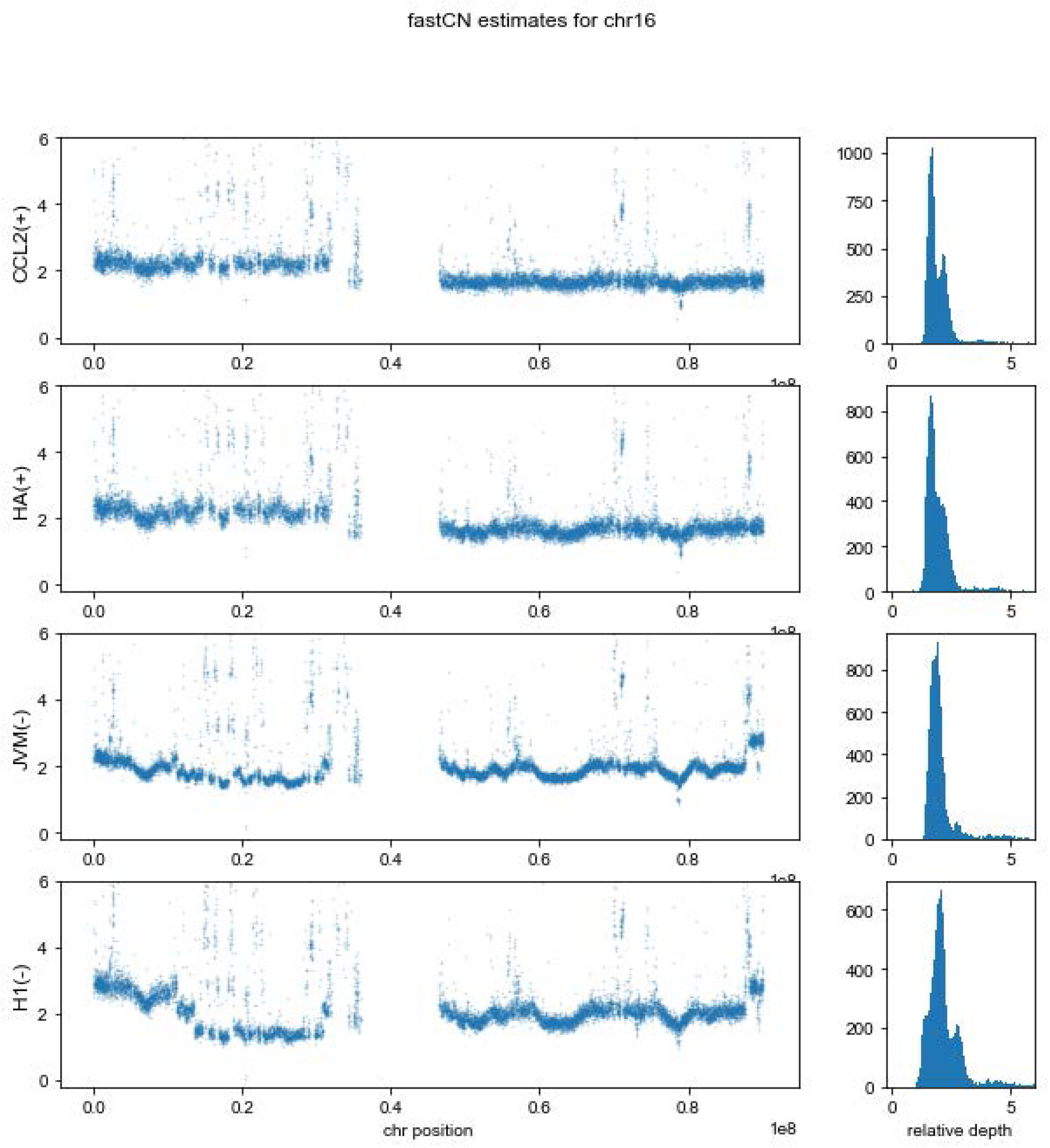
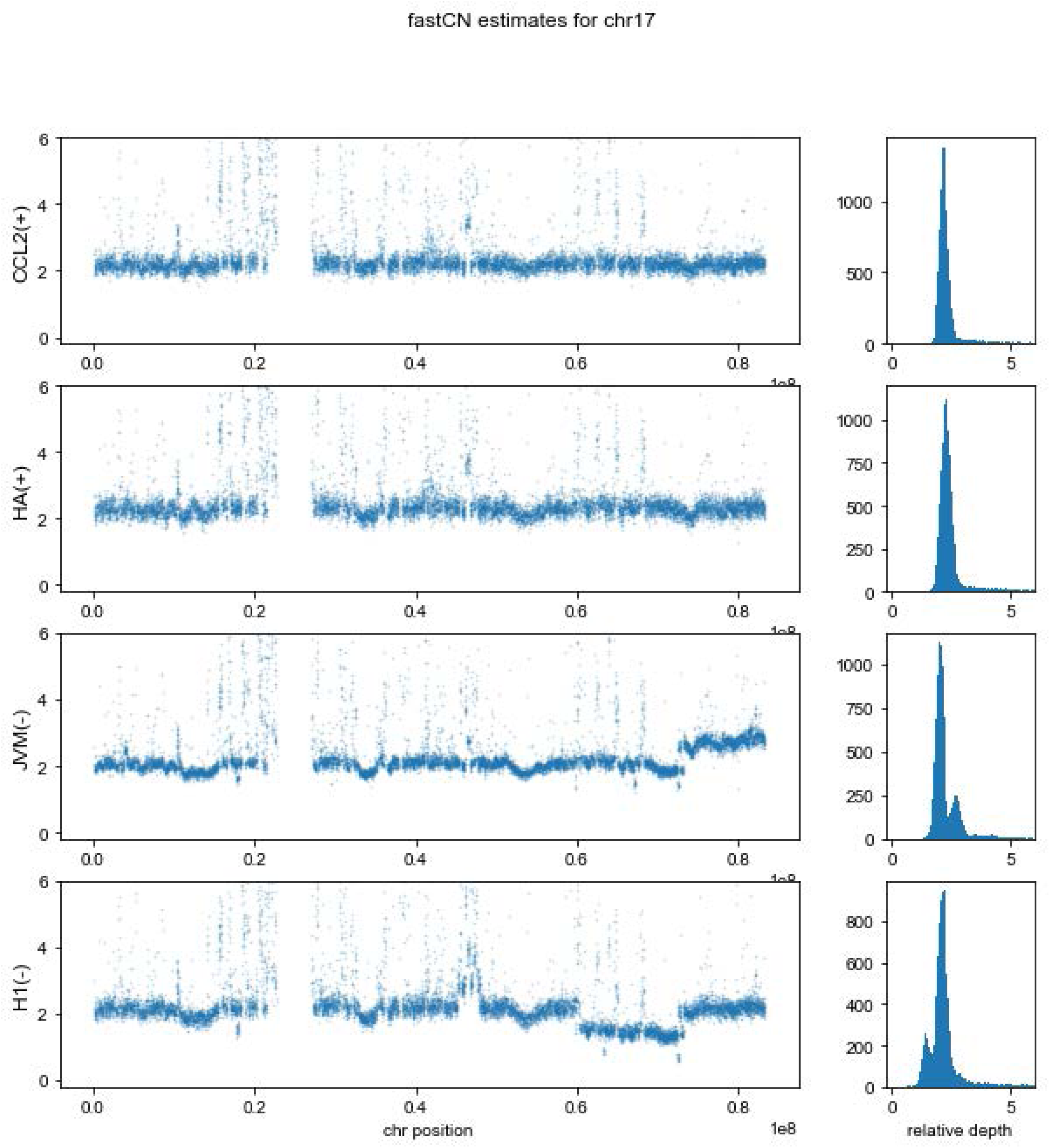
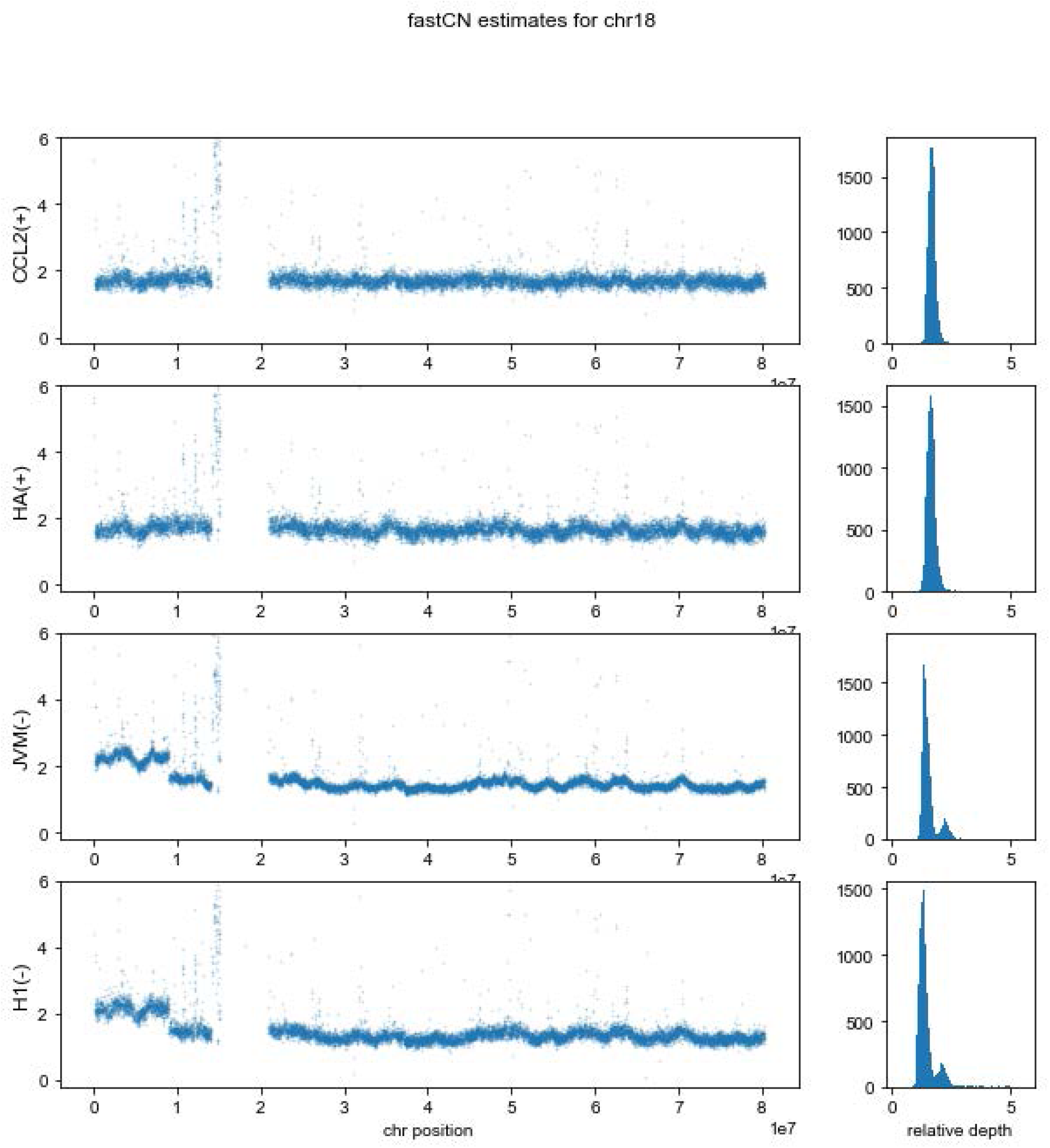
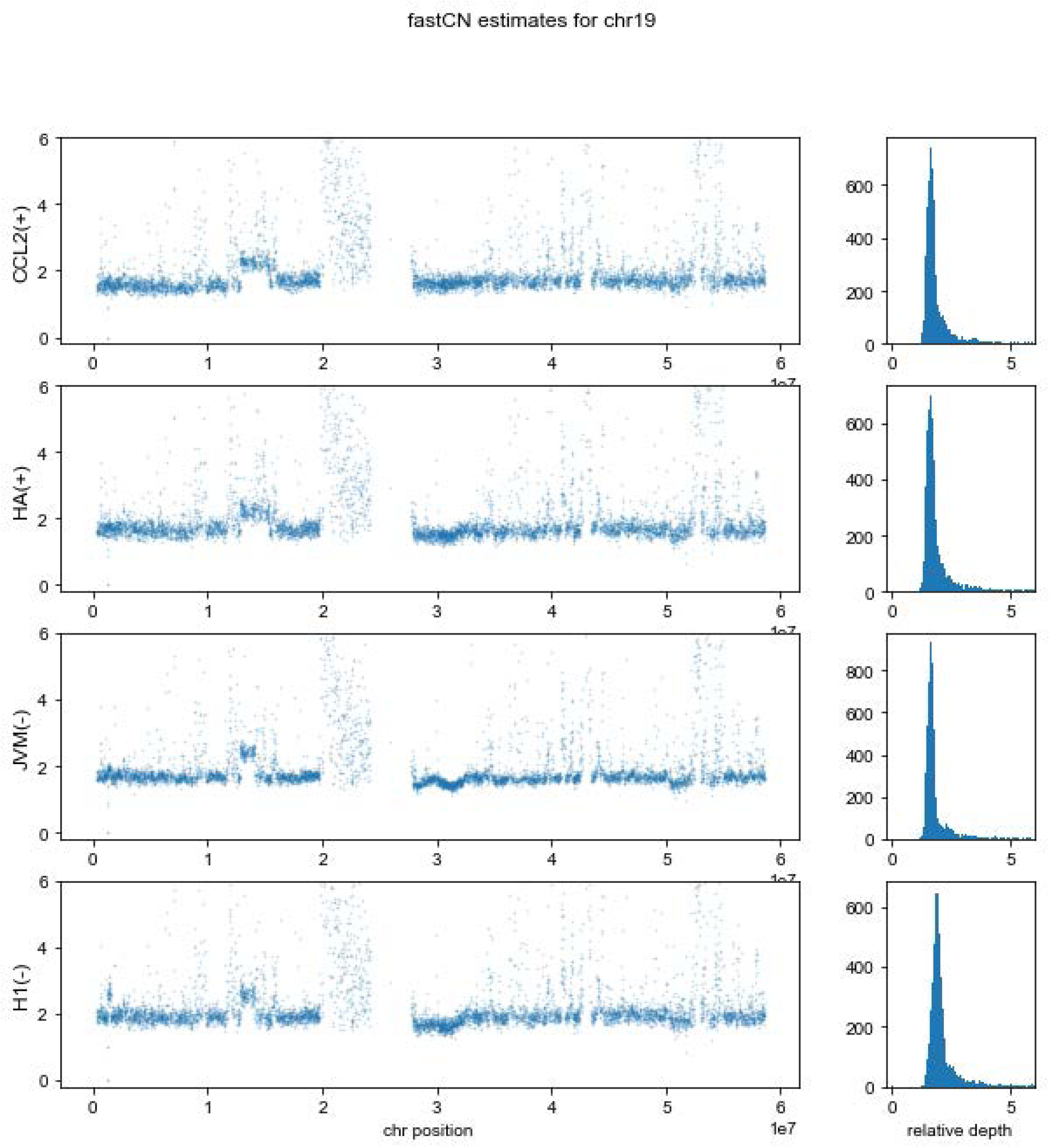
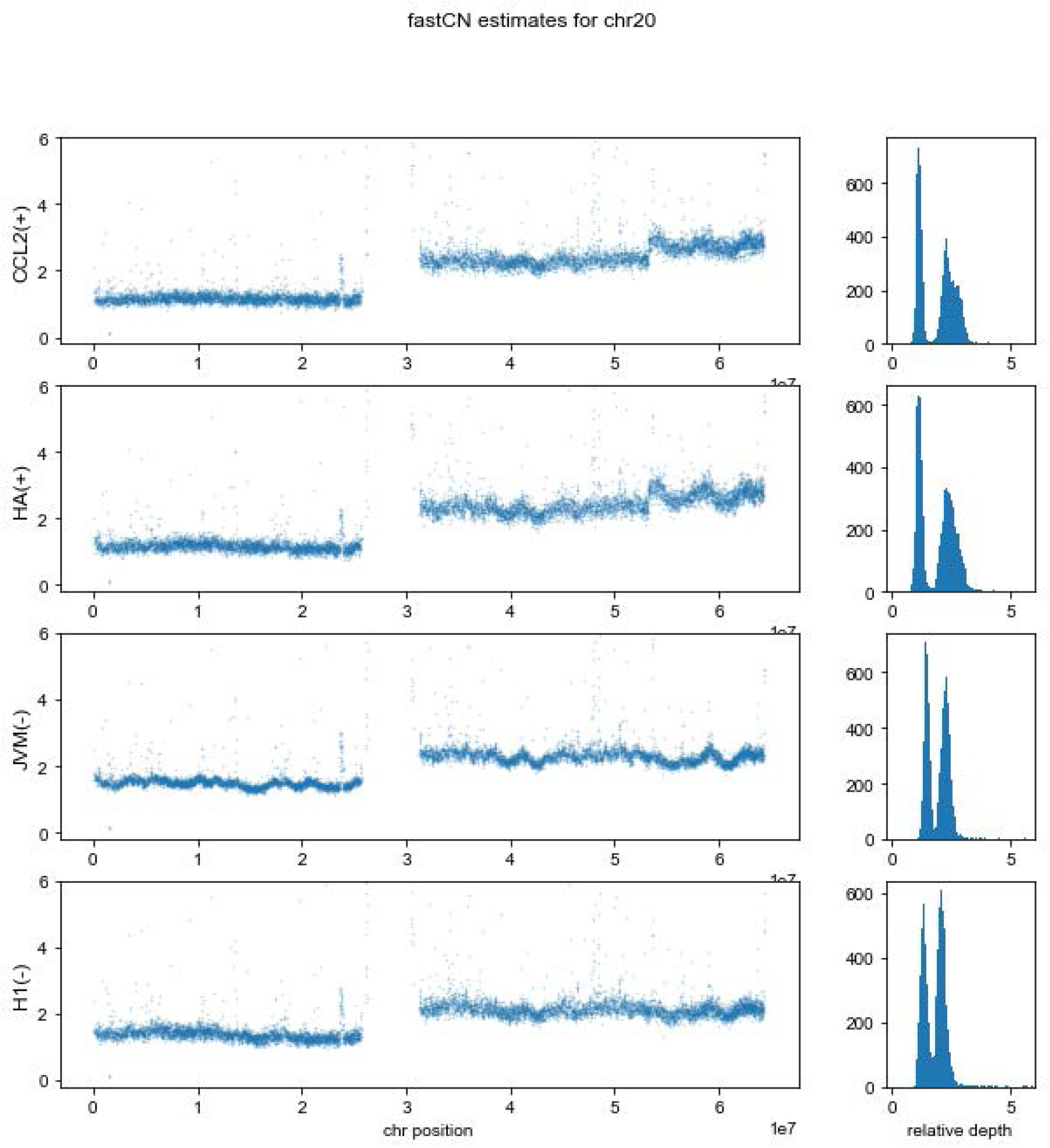
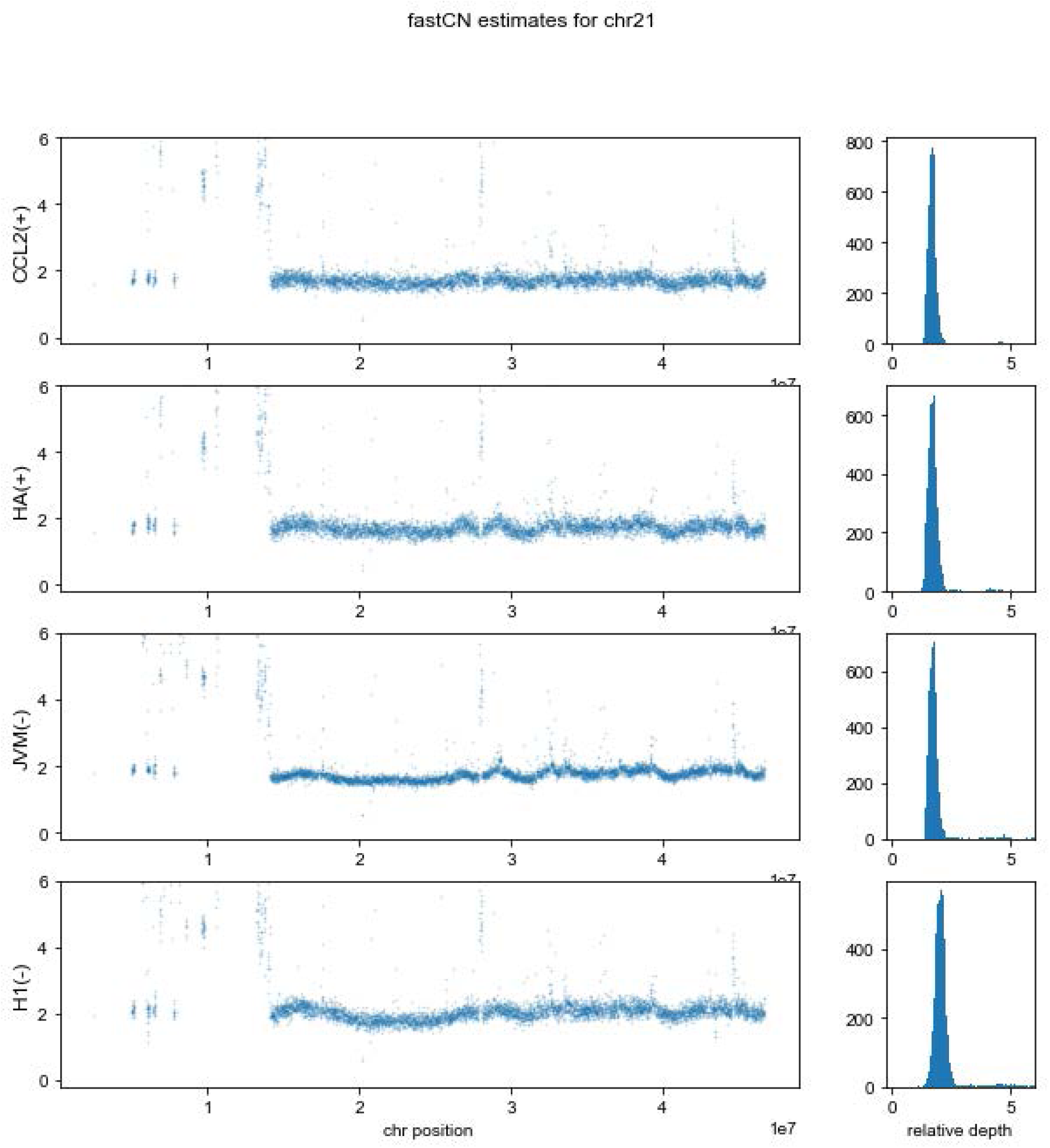
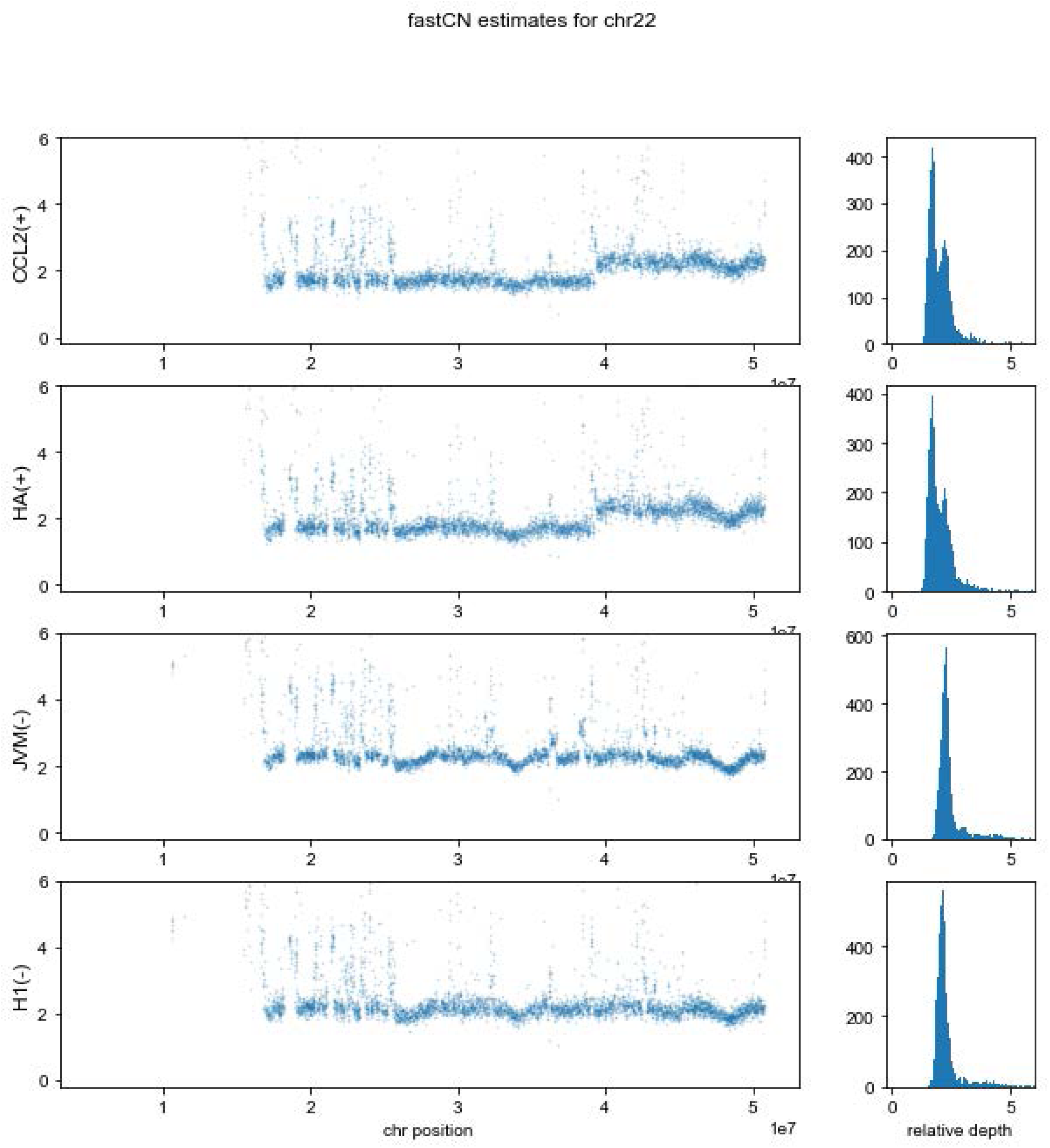
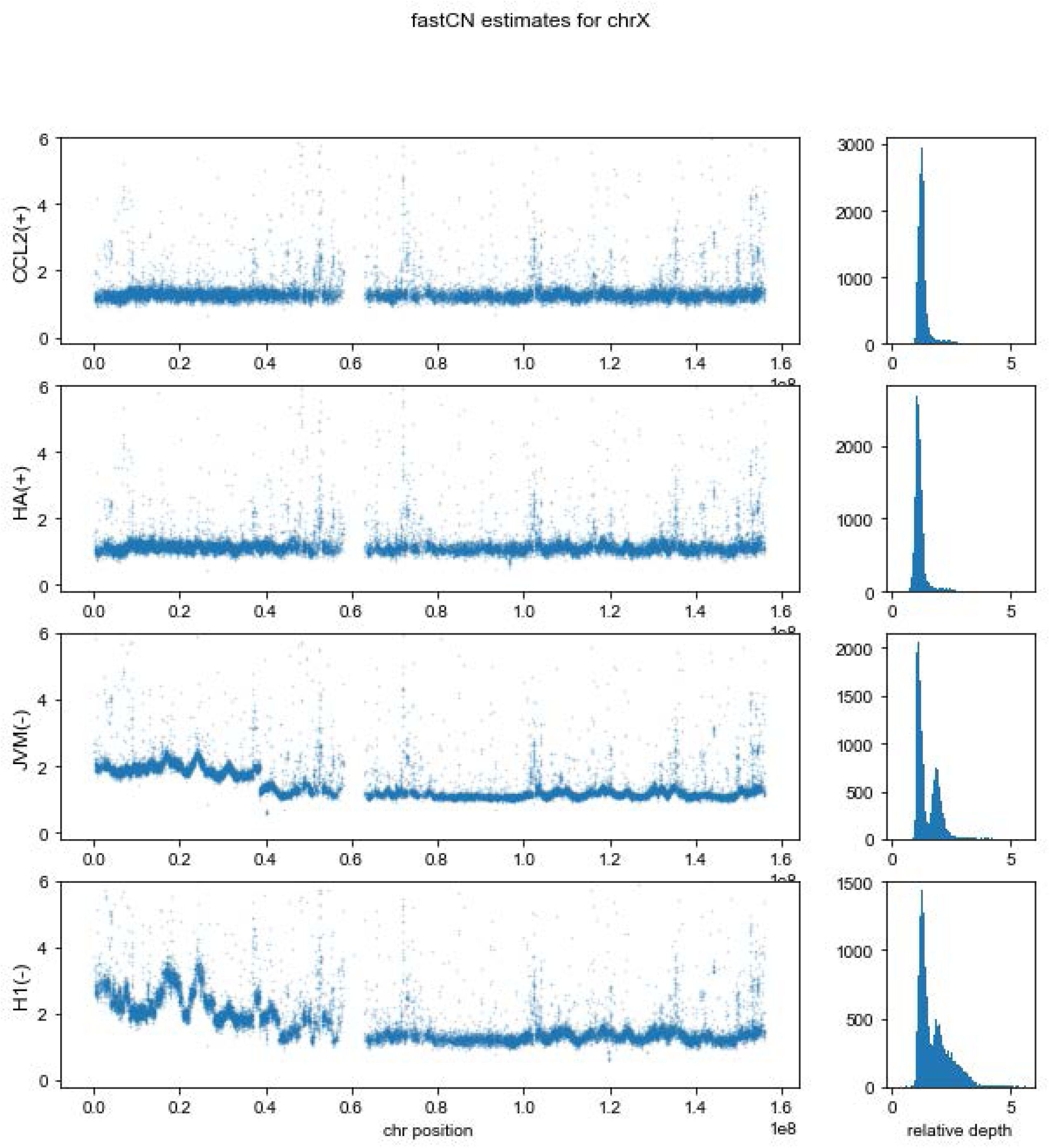
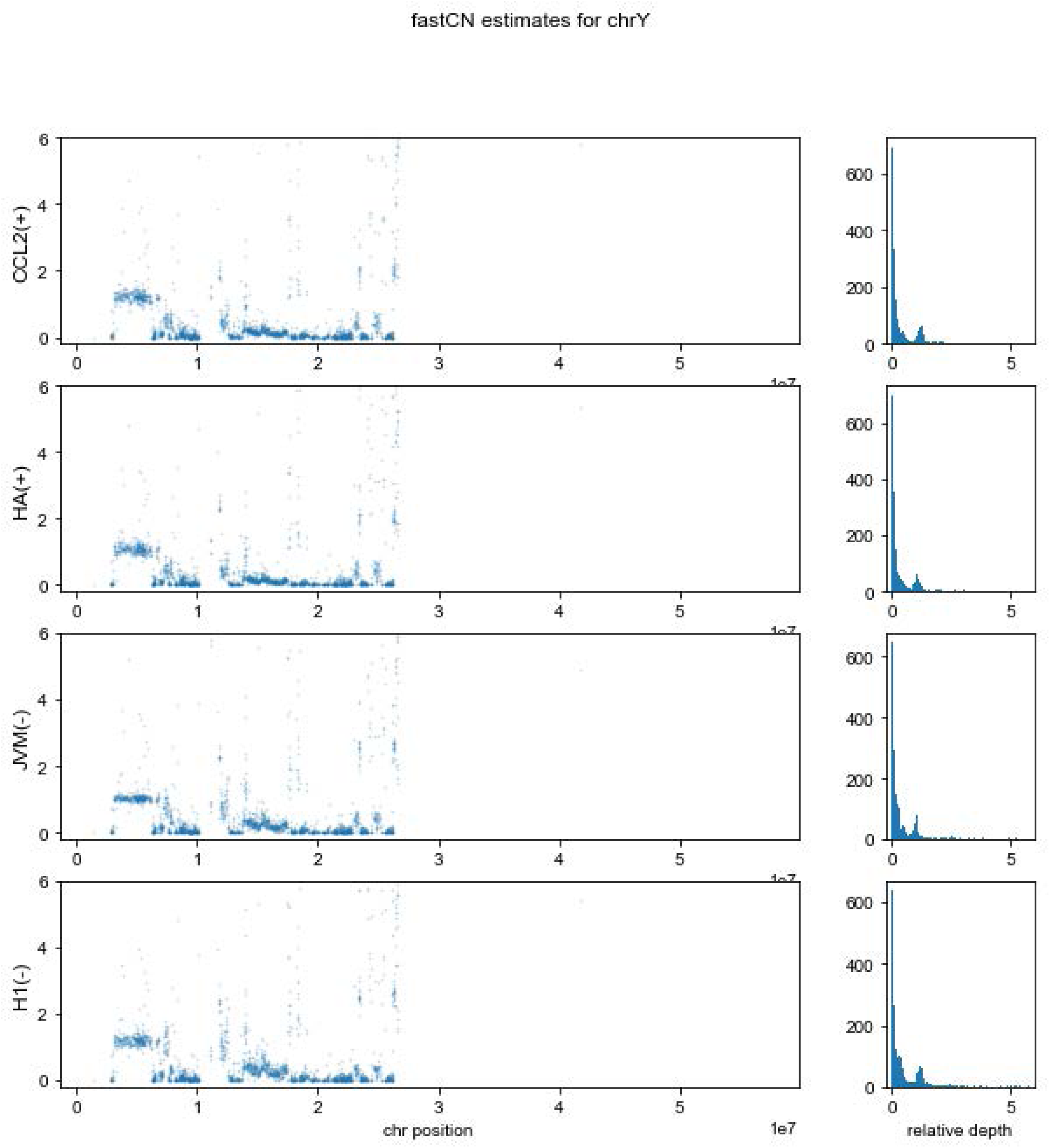
*(A) HLA typing of HeLa-HA and HeLa-JVM cell lines.* From left to right, each panel shows HLA-typing results for DR*B1*, DQ*B1*, HLA-A, HLA-B and HLA-Cw in the indicated HeLa cell lines (indicated in the top using the “JVM” and ”HA” labels). The boxed text displays the inferred genotype of both HeLa-HA and HeLa-JVM strains. *(B) Short Tandem Repeat (STR) analyses in HeLa cells.* Shown are results from multiplexed STR analyses resolved on an ABI PRISM 3130xl Genetic analyzer (PE Applied Biosystem). The left and right panels show results for MIX.15 (*e.g.*, D15S146 + D15S1028 + D15S126 + D15S209 + D15S153) and MIX.2 (*e.g.*, D6S105 + D6S265 + D6S273 + C.1.2.5), respectively, in HeLa-JVM (top) and HeLa-HA (bottom). *(C) Each panel shows the GeneMapperTM (PE Applied Biosystem) profile of the indicated marker (top) in HeLa-JVM (top row) and HeLa-HA (bottom row*) *(D) Copy number (CN) estimates for the four HeLa strains.* Graphs depict CN estimates for each chromosome of each HeLa cell strain (indicated to the left of each plot). The X-axis indicates chromosome position; the Y-axis indicates CN.

**Supplementary Figure S3, Related to Figure 3:**
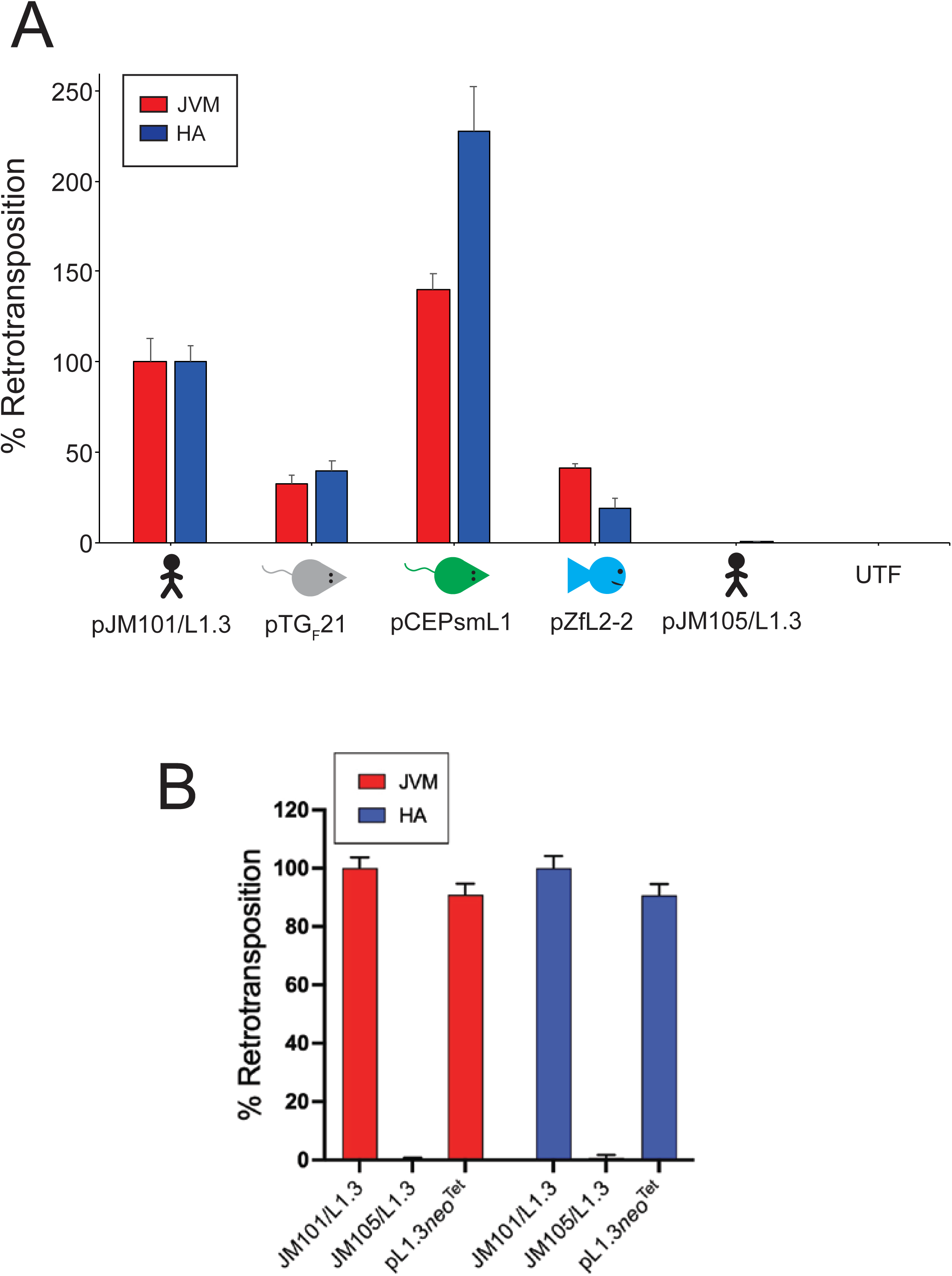
*(A) Quantification of LINE retrotransposition data from* Figure 3A: The X-axis indicates LINE construct, and the Y-axis indicates average % retrotransposition efficiency normalized to pJM101/L1.3. Each experimental transfection condition was repeated at least three times with similar results. Error bars represent standard deviation. *(B) Quantification of the pL1.3neo^Tet^ retrotransposition assay.* The X-axis indicates the L1 construct, and the Y-axis indicates average % retrotransposition efficiency normalized to pJM101/L1.3. Each experimental transfection condition was repeated at least 3 times with similar results. Error bars represent standard deviation.

**Supplementary Figure S4, Related to Figure 5:**
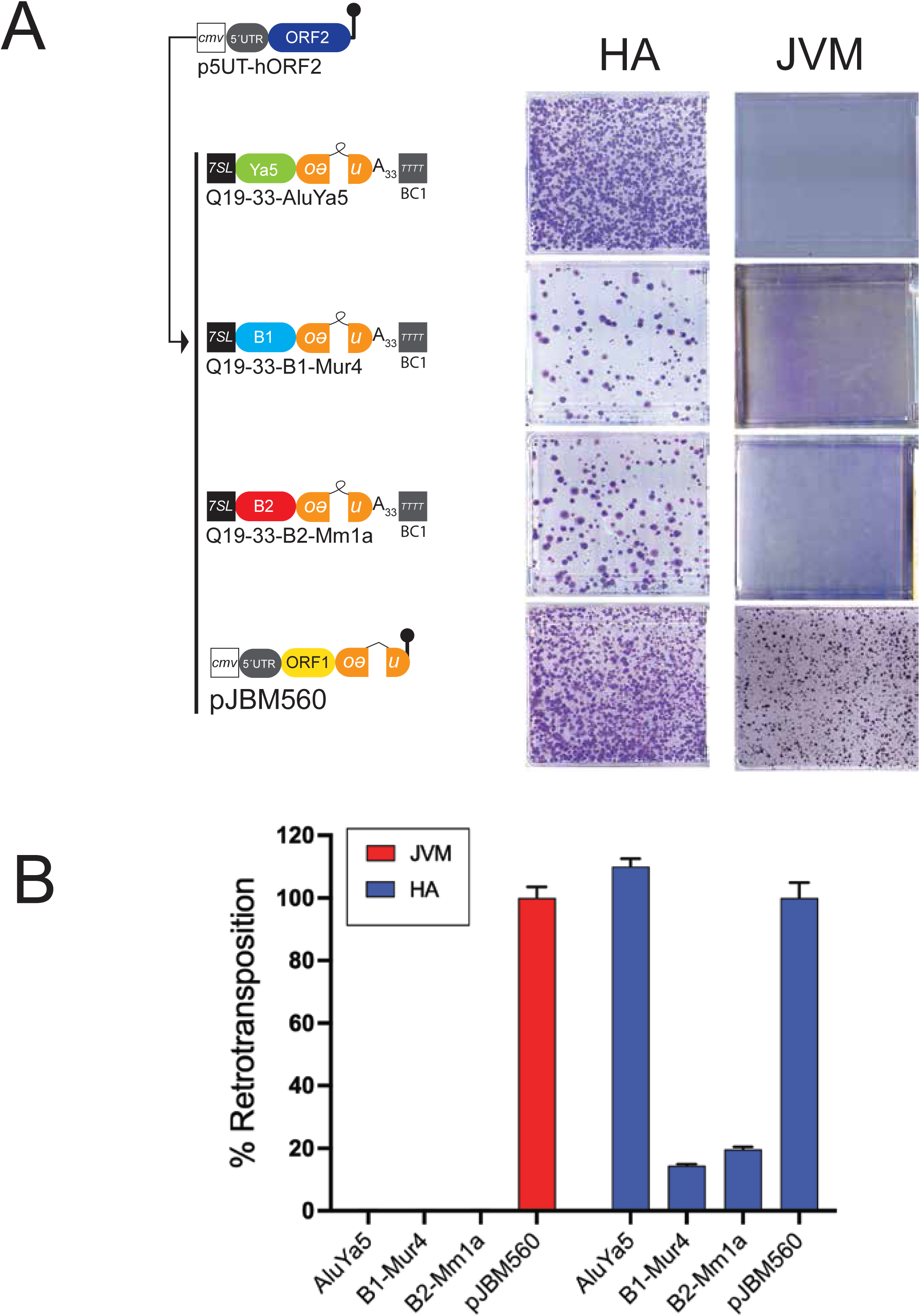
*(A) Retrotransposition results for mouse SINEs*. HeLa cells were co-transfected with a “driver” p5UT-hORF2 plasmid and either Q19-33-*Alu*Ya5, Q19-33-*B1*-Mur4, Q19-33-*B2*-Mm1a (see main text for details), or *ORF1mneoI* (pJBM560) (26) “reporter” plasmids. Displayed are images of representative T-75 flasks from the retrotransposition assays. The HeLa strain is denoted above each column of images. (B) *Quantification of mouse SINE retrotransposition assays.* The X-axis indicates SINE construct co-transfected with p5UT-hORF2 and the Y-axis indicates average % retrotransposition efficiency normalized to pJBM560. Shown is the average of three independent biological replicates; error bars represent standard deviation.

**Supplementary Figure S5, Related to Figure 7:**
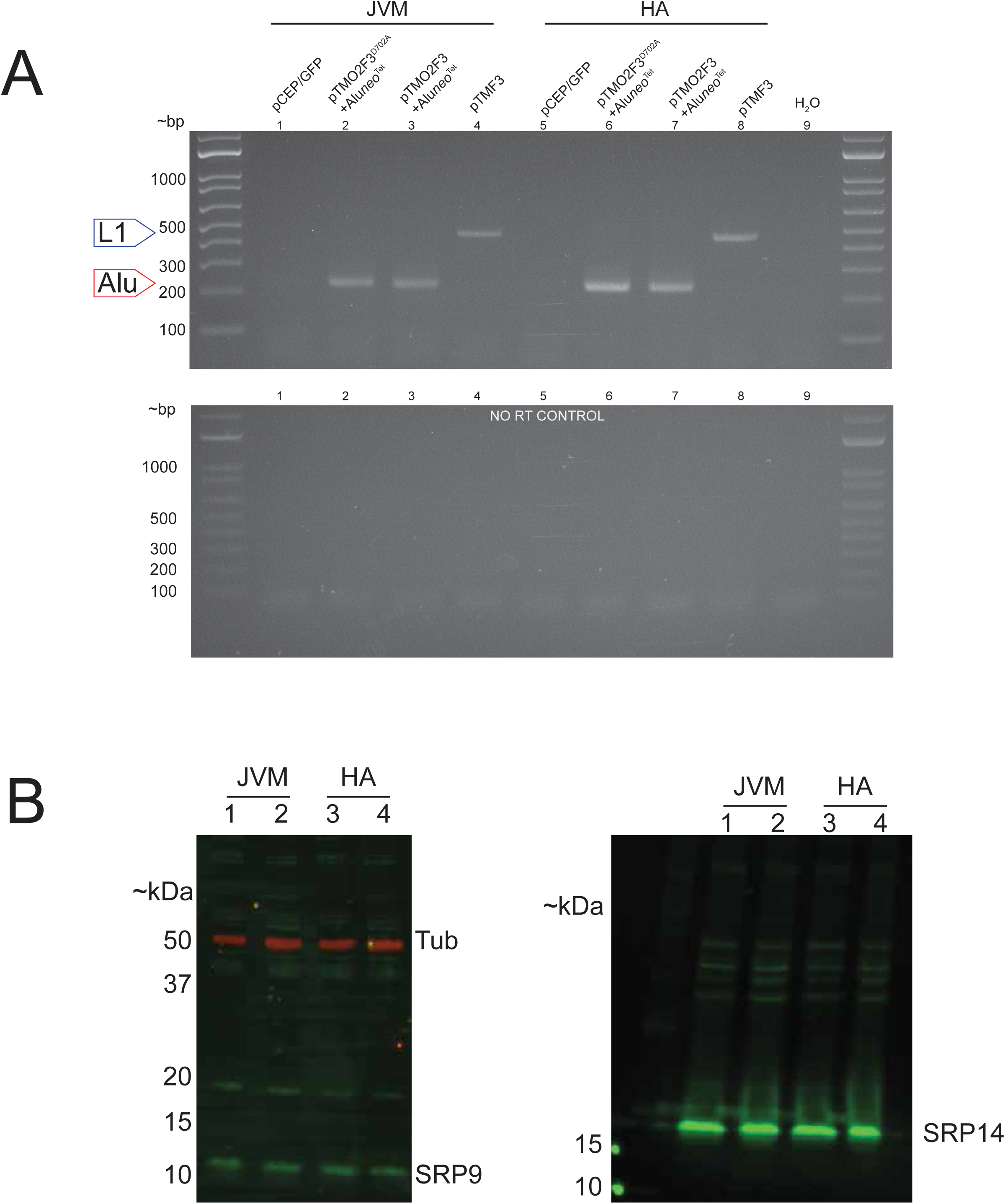
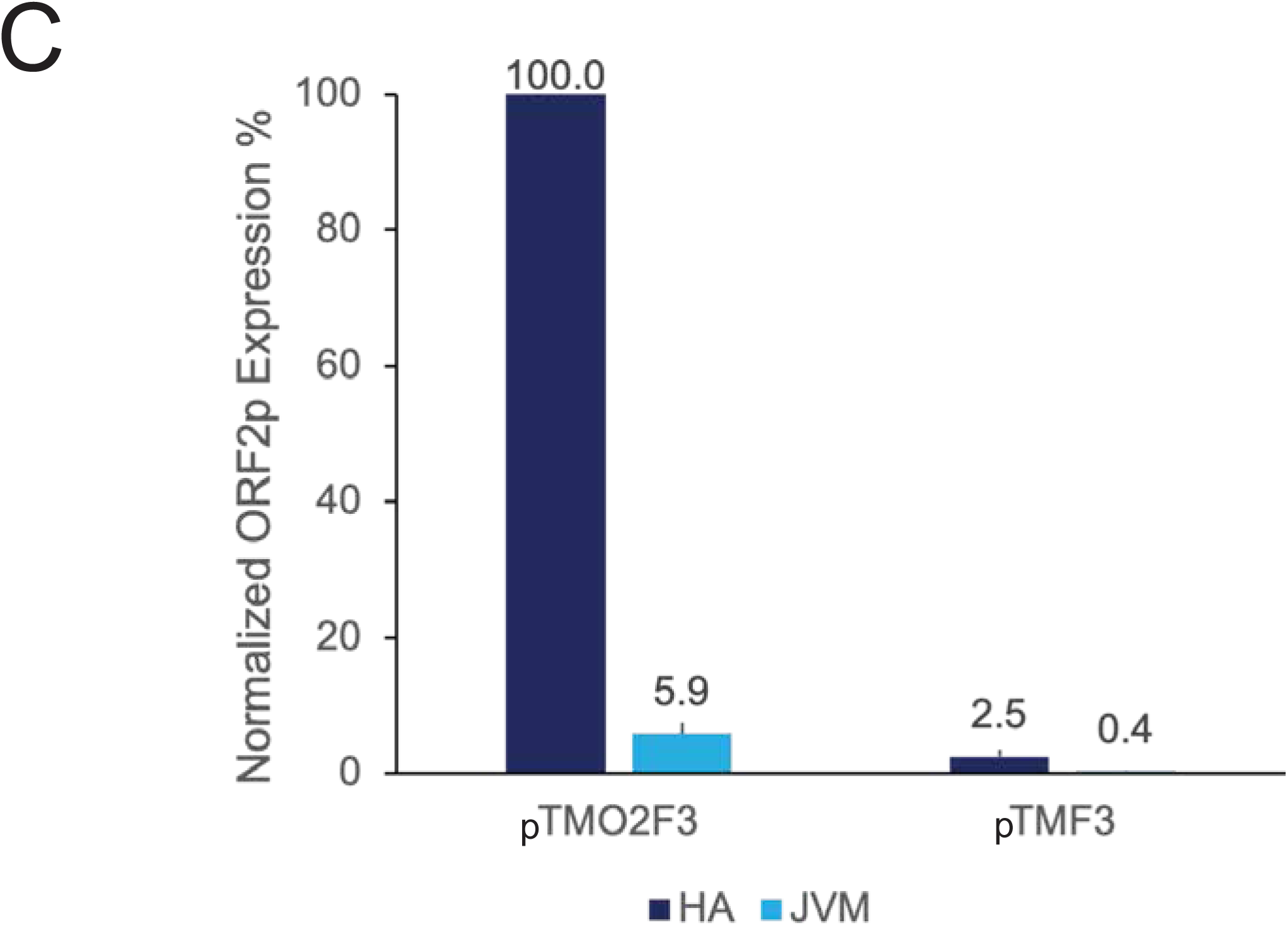
*(A) MMLV RT-PCR results:* Transfection conditions and HeLa cell lines are indicated at the top of the gel image. The approximate sizes of *Alu* RT-PCR products (∼240 bp; red arrow) and L1 RT-PCR products (∼420 bp; blue arrow) are indicated on the left of the gel image. The L1 RT-PCR products are larger due to the presence of the pCEP4 SV40 polyadenylation signal sequence at the end of the *mneoI* indicator in pTMF3. Bottom gel = No RT control reactions; H_2_O = PCR control; DNA size markers (bp) are shown at the left of the gel. *(B) SRP9/14 Western Blot results from HeLa-JVM and HeLa-HA.* Shown are Srp9 (left gel image) and Srp14 (right gel image) western blots from HeLa cell whole cell lysates. HeLa-JVM results are shown in lanes 1-2 and HeLa-HA results are shown in lanes 3-4. Lanes 1 & 3 = HeLa transfected with pCEP-GFP; Lanes 2 & 4 = HeLa co-transfected with pTMO2F3 and p*Aluneo*^Tet^. Tubulin (Tub) served as a loading control. *(C) Quantification of steady-state ORF2p expression in whole cell lysates from HeLa-HA (blue) and HeLa-JVM (light blue) from LEAP experiments.* The X-axis indicates the LINE-1 expression plasmid. The Y-axis indicates the ORF2p steady-state expression level normalized to HeLa-HA cells transfected with pTMO2F3.

## SUPPLEMENTAL TABLE LEGENDS

**Supplementary Table 1, Related to Fig S2B.** Primers used in short tandem repeat (STR) analyses. Column 1, STR name. Column 2, STR size. Column 3, STR marker type. Column 4, PCR primer sequences.

**Supplementary Table 2, Related to Fig S2C.** Summary of markers used in the *GeneMapper* STR analyses.

**Supplementary Table 3, Related to Fig 2B.** Summary of G-banding karyotype data summary for the HeLa-HA and HeLa-JVM strains.

**Supplementary Table 4, Related to Fig 4.** Structures of *Alu* insertions characterized from HeLa-HA cells. Column A: name of insertion. Column B: genomic DNA sequence flanking the *Alu* insertion; all sequences listed in the 5’ to 3’ direction. Column C: estimated poly(A) tail length. Column D: length of target site duplication (TSD). Column E: L1 endonuclease (EN) cleavage site sequence on the bottom strand. The “/” indicates the L1 EN cleavage site. Column F: the genomic location of the L1 EN cleavage site based on GRCh37/Hg19. Column G: notes regarding the insertion sequence if applicable.

**Supplementary Table 5, Related to Fig 4.** Structures of *Alu* insertions characterized from HeLa-JVM cells. Column A: name of insertion. Column B: genomic DNA sequence flanking the *Alu* insertion; all sequences listed in the 5’ to 3’ direction. Column C: estimated poly(A) tail length. Column D: length of target site duplication (TSD). Column E: L1 endonuclease (EN) cleavage site sequence on the bottom strand. The “/” indicates the cut site. Column F: the genomic location of the L1 EN cleavage site based on GRCh37/Hg19. Column G: notes regarding the insertion sequence if applicable.

